# Artificial intelligence and species distribution ensemble models inform resource interactions with offshore wind development

**DOI:** 10.1101/2024.06.10.598232

**Authors:** Evan C. Ingram, Keith J. Dunton, Michael G. Frisk, Liam Butler

**Affiliations:** School of Marine and Atmospheric Sciences, Stony Brook University, Stony Brook, New York 11790, USA; Monmouth University, West Long Branch, New Jersey 07764, USA; Wake Forest University, Winston-Salem, North Carolina 27109, USA

## Abstract

Development of offshore wind energy resources has led to growing concerns for marine wildlife. However, significant uncertainty remains regarding the technology’s potential to impact species of interest that may occupy planned development sites. This is further compounded by the difficulty of monitoring highly migratory or data-poor species in marine waters, making practical assessment of site- or species-specific threats that could require additional management intervention particularly problematic. Here, I identify a highly generalizable framework to inform species interactions in marine habitats allocated for offshore resource exploitation, using telemetry-derived artificial intelligence species distribution models. Results from a case study of the federally protected Atlantic Sturgeon (*Acipenser oxyrinchus*) demonstrate excellent discriminatory capacity (i.e., AUC ≥ 0.9) at a relatively fine scale (raster resolution = 1 km^2^), while providing critical information on predicted occurrence over a broad swath of unmonitored marine habitats (i.e., the Atlantic OCS region of the US; area > 620,000 km^2^). Furthermore, ensemble map products developed from these models are readily scalable to ongoing management needs and, when overlaid with offshore wind energy lease areas, can feed directly into management strategies to inform best practices for potential habitat influences on Atlantic Sturgeon, as well as other species of commercial or conservation interest.

## Introduction

The accelerated commercialization of offshore wind energy has prompted significant concerns for marine wildlife (Akhtar et al., 2022). With global wind farm capacity expected to increase exponentially over the coming decades, there is an urgent need to advance data-driven approaches capable of guiding management and conservation responses to commercial-scale development (Gill, 2005; Boehlert and Gill, 2010). However, our current understanding of marine autecology in habitats allocated for offshore wind exploitation is largely limited and remains unresolved for many species of concern (e.g., Drewitt and Langston, 2006; Bergstrom et al., 2013; Bailey et al., 2014; Bergstrom et al., 2014). Critical data-gaps in the literature regarding the potential for spatiotemporal overlap between species movements distributions and offshore renewable activities are only further compounded by the difficulty of monitoring cryptic or highly migratory populations across broad, spatially heterogeneous marine areas—increasing the potential for regulatory incongruities as offshore development timelines outpace the ability of resource managers to identify appropriate conservation strategies (Boehlert and Gill, 2010; Nazir et al., 2021)

In the United States, aggressive clean energy targets and favorable domestic policies have driven rapid growth in the offshore wind energy market (Sun et al., 2020; Alola and Akadiri, 2021). Wind energy areas have been identified throughout the waters of the US Outer Continental Shelf (OCS), with active commercial wind leases and near-term projects primarily concentrated in the Northeast and Mid-Atlantic regions (Table S1, Supporting Information). As domestic efforts in the OCS continue to scale up to meet the demand for renewable energy production, consideration of ecological effects across the entirety of these projects has become a necessarily complex undertaking. While stressors to marine ecosystems and fauna are expected during all phases of development, the lack of scale-appropriate information needed to properly identify and mitigate potential resource conflicts—and the challenge of obtaining such data over a large marine area on a site-by-site basis—could prevent adequate impact assessment or lead to substantial delays in the consenting process (Gill, 2005; Boehlert and Gill, 2010).

Recent studies have underscored the importance of spatial and temporal considerations when assessing the effects of offshore wind development on marine fauna (e.g., Gusatu et al., 2021; Galparsoro et al., 2022). The main impacts of operational offshore wind farms—both positive and negative—are often locally influenced and contingent on specific site dynamics. However, the geographical footprint of these developments is expected to increase alongside expected advancements in offshore wind technology that allow for the deployment of larger and denser arrays in areas favorable for development. As such, integration of ecological data across multiple scales is critical to informing planning and decision-making processes throughout the OCS (Gill, 2005). This is especially important when designing impact assessments for species of concern that may range widely or with limited or no existing information on offshore habitat use or distribution.

Species distribution modelling (SDM) are correlative-predictive models that have become critical tools for conservation (Franklin 1995; Guisan and Thuiller, 2005; Miller and Rogan, 2007; Elith and Leathwick, 2009; Franklin, 2010). Importantly, SDMs allow researchers to quantify species-biogeographic relationships at scales that are appropriate to a variety of conservation and management applications. These models are readily interpreted and can provide actionable insights into factors influencing patterns of biodiversity. Spatial products (i.e., mapped predications) generated from SDMs are useful as inputs to other analyses, particularly in conservation planning or environmental impact assessments. SDMs have high discriminatory capability and can provide predictions of occurrence outside of sampled areas, allowing extrapolation from empirical models to make inferences about novel environments (e.g., Alberto Jiménez-Valverde et al., 2013). The use of artificial intelligence (AI) algorithms in SDMs to estimate relationships between population locations and environmental conditions can provide additional flexibility over traditional statistical models (i.e, regression-based techniques, such as GLMs, GAMs, or MARS, which can have limited application due to sample requirements or unrealistic assumptions of linearity) and an increased ability to extrapolate predictions over broad extents of unmonitored habitat (e.g., Leathwick et al., 2005; Dorman et al., 2007; Miller et al., 2007; Miller, 2010). The further use of ensemble modeling (i.e., weighted consensus prediction) improves prediction accuracy and can largely reduce potential biases associated with individual algorithms (Araújo and New, 2007; Marmion et al., 2009b; Thuiller et al., 2009).

Here we identify a generalized framework for the use of AI-based SDM ensemble models to predict the extent of species habitats in areas allocated for wind energy development, using the federally endangered Atlantic Sturgeon (*Acipenser oxyrinchus oxyrinchus*) as a case study. Atlantic Sturgeon are long-lived, anadromous fishes that migrate extensively in waters of the Atlantic OCS region during periods of marine residency and may occupy multiple wind energy development sites on a seasonal basis (e.g., Stein et al., 2004a; Stein et al., 2004b; Dunton et al., 2010; Erickson et al., 2011; Breece et al, 2017; Melnychuk et al., 2017; Ingram et al., 2019; Rothermel et al., 2020). Quantification of the extent of spatial overlap between Atlantic Sturgeon and wind energy developments in coastal and offshore habitats is necessary to identify and effectively address site- or population-specific threats that may require additional management intervention. However, spatiotemporal patterns of habitat use and occupancy throughout their marine distribution are largely unknown and limited information is available at scales relevant to management needs. Despite the recent proliferation of long-term acoustic telemetry tagging programs along the Atlantic coast of the US (see Bangley et al., 2020), current monitoring and programs for Atlantic Sturgeon are generally limited to riverine or nearshore waters, which can lead to underestimation of offshore habitat use and potential delays in management responses to emergent threats. As such, the goal of this case study was to develop SDMs for Atlantic Sturgeon that leverage existing, high-resolution datasets and advanced AI algorithms to predict the spatiotemporal occurrence in marine waters of the Atlantic OCS. The specific objectives were to 1) quantify the distribution of Atlantic Sturgeon across the entire region, 2) identify spatiotemporal trends in habitat use at sites allocated for offshore wind energy development, and 3) generate detailed ensemble map products capable of feeding directly into management and decision making at multiple scales.

## Methodology

### Study region and spatially referenced observations

Potential offshore habitats for Atlantic Sturgeon are located throughout the Atlantic OCS region of the United States. The boundaries of the Atlantic OCS, from the coast to the shelf break, were used to provide a clipping extent for subsequent map layers (total area = 626,204 km^2^; Figure 1). This extent also included several large estuaries that are seasonally occupied by Atlantic Sturgeon, including Long Island Sound, Delaware Bay, and Chesapeake Bay. Telemetry detection records were compiled for acoustically tagged Atlantic Sturgeon originally captured in coastal and marine waters of the New York Bight (Table S2). Acoustic detections of these fish were available from collaborative agencies operating acoustic receiver arrays throughout the Atlantic OCS during 2011 through 2021 (Table S3). Unique detection locations across the full study period were pooled by month (i.e., monthly occurrence data) and input into R (R Core Team, 2018) as spatial data frame objects (n =12; ‘SF’ package; Pebesma. 2019). The points of these records were aligned with centroids of the relevant 1-km raster cells in the OCS study site map.

**Figure 1.**
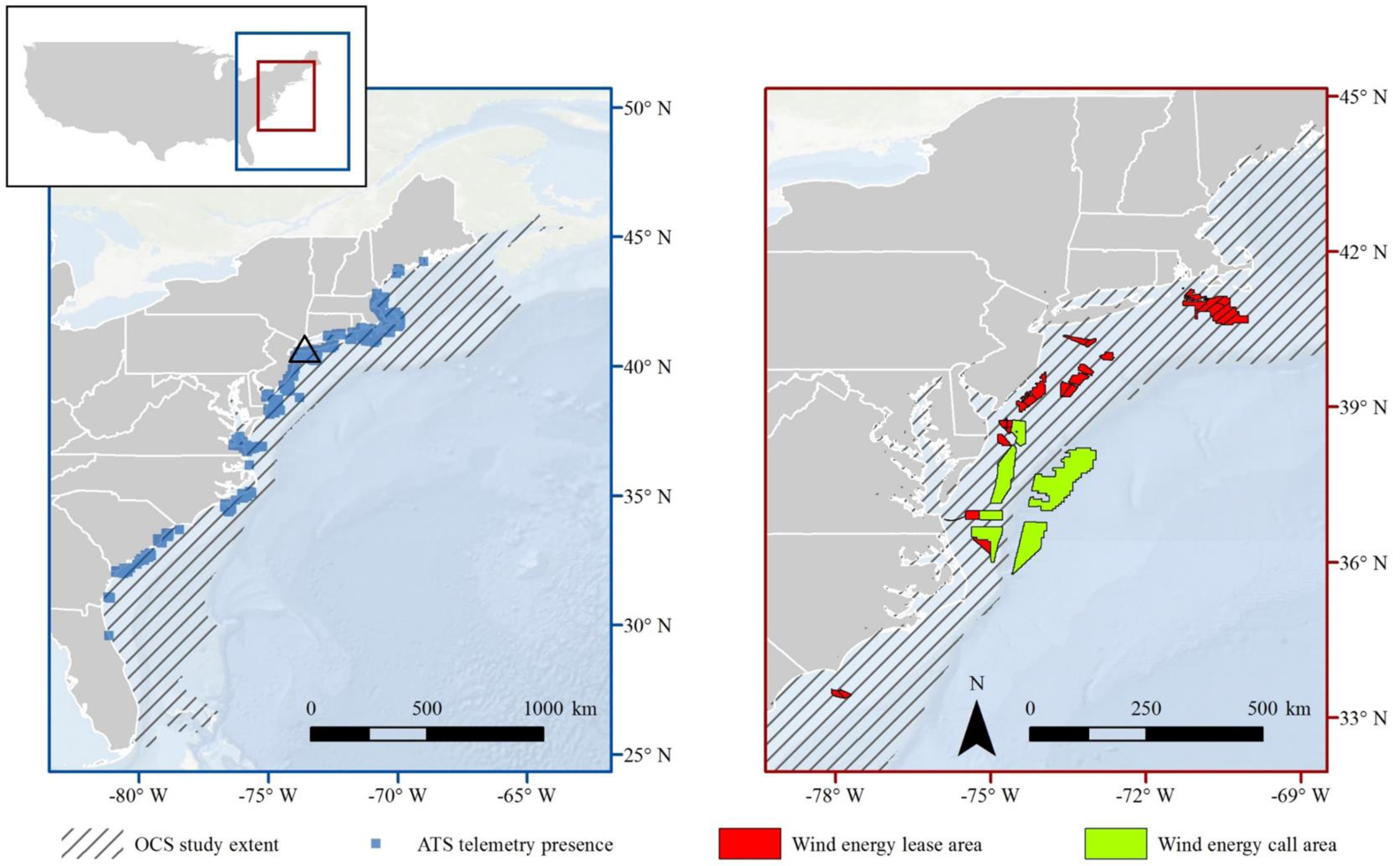
Extent of the United States Atlantic Coast Outer Continental Shelf study site showing unique Atlantic Sturgeon locations from acoustic telemetry monitoring (n = 845; left panel) and planned offshore wind energy development sites (right panel). Atlantic Sturgeon typically sampled in marine waters off New York (open triangle). Note that maps are shown at different scales.

### Environmental predictors

Explanatory variables for modelling efforts were selected based on putative biological significance and data availability across the entirety of the study (Table S4). To ensure generalizability, only publicly accessible datasets were considered. If an environmental gradient included a temporal aspect, pooled monthly averages were calculated for the entire period of interest. Variance inflation factors (VIFs; Marquardt, 1970) were calculated and predictors with VIF values indicating high correlation (i.e., VIF ≥ 10) were removed from further analysis (Chatterjee and Hadi, 2015). The final set of six explanatory variables was used across all models, and included: depth (m), percentage of gravel substrate (%), percentage of mud substrate (%), photoperiod (hours/day), precipitation (kg/m^2^), and sea surface temperature (SST; °C). These environmental predictors were stored as a set of raster layers (i.e., a raster stack) with the same spatial extent (i.e., World Geodetic System 1984; EPSG:4326) and resolution (i.e., 1-km resolution) to allow for interoperability and consistency among the different data sources.

### Species distribution models

Species distribution models were developed and evaluated using the ‘SDM R’ package (Naimi and Araújo, 2016) in R (R Core Team, 2018). Species distribution models were fit using seven methods: classification and regression trees (CART), gradient boosting (GBM), maximum entropy (MAXENT), multilayer perceptron (MLP), radial basis function (RBF), random forest (RF) models, and support vector machine models (SVMs). These methods consisted entirely of artificial intelligence algorithms, including machine learning (i.e, CART, GBM, MAXENT, RF, and SVM) and artificial neural networks (i.e., MLP and RBF). The default settings for tuning and regularization of parameters were used for all seven models (see Table S5). Because telemetry records consisted of presence-only data (i.e., occurrence), random pseudoabsence datapoints (i.e., background data) were generated to provide absence points for each month (n = 1000 datapoints per bootstrap; Phillips and Dudik, 2008; Barbet-Massin et al., 2012). While certain algorithms in species distribution modeling can use presence-only records to establish the conditions under which a species is more likely to be present (e.g., MAXENT; see Elith et al., 2011), others require background data to establish the environmental domain of the study. However, the performance of presence–absence models is generally better than presence-only models (Elith et al., 2006). Additionally, standard evaluation parameters are based on measured of both sensitivity (correctly predicted presences) and specificity (correctly predicted absences; Fielding and Bell, 1997). Because of this, presence-absence models are recommended when only presence data is available by creating artificial background data (Barbet Massin et al., 2012).

### Model quality assessment

While the comparison of model predictions against a completely independent dataset is the most robust approach to assessing model accuracy (Araujo et al., 2005), the equivalent of such data were not available for this study. Instead, data-dependent testing was performed on 20% of the original dataset, while the remainder was used for model training. Model evaluation relied on resampling techniques that used bootstrapping validation to generate multiple datasets (Hastie et al., 2009). Repeated samples were drawn from the original observation data with replacement. Each bootstrap replication was repeated 100 times per algorithm (Barbet-Massin et al., 2012); i.e., each of the seven algorithms was averaged across 100 bootstraps, resulting in a total of 700 models for each month. Model quality metrics were assessed using area under the receiver operating curve (AUC), true skill statistic (TSS), and Kappa index. Both AUC and TSS measure the discriminatory ability of an SDM. AUC is a threshold-independent measure of model performance that compares the true positive rate of observations (i.e., sensitivity) against the false positive rate (1 – specificity; where specificity is the proportion of negatives that can be correctly identified; Fielding and Bell, 1997). TSS is another relatively simple measure of model performance that is calculated as sensitivity + specificity – 1 (Allouche et al., 2006). The Kappa index is a threshold dependent correlation metric (i.e., threshold = 0.4) that is widely used measure to compare observations and predictions, and varies from −1 to 1, with values of 0 or less = no better than random, and values of 1 = perfect agreement (Cohen, 1960).

### Predictor importance

The importance of individual predictor variables was determined following the methods of Thuiller et al. (2009). Pearson correlations were calculated between values predicted for the original environmental data and values predicted for data in which a variable of interest had been randomly permuted. High correlation between the two predictions indicates that the environmental variable was not important. For comparative purposes, metrics for the Pearson correlation are reported here as ‘1 – correlation’ so that more important variables have higher scores.

### Ensemble map products

Ensemble maps of predicted Atlantic Sturgeon occurrence across the study site were generated for each month using the ‘ESDM’ package (Woodman et al., 2019) in R (R Core Team, 2018). Only SDMs that met a high AUC threshold (i.e., AUC > 0.8; Thuiller et al., 2019) were retained for the final ensemble projections. Final ensemble predictions were informed by weighted mean values of the AUC for each retained algorithm for each month (Table S6). The final map products were overlaid with shape files of wind energy development sites in the Atlantic OCS to identify areas of potential spatial or temporal resource conflicts. As such, ensemble maps could be further manipulated in ArcGIS (ESRI, 2020) to allow for focus on specific sites or regions of interest.

## Results

### Spatially referenced observations

All Atlantic Sturgeon described in this study (n = 599) were originally sampled in riverine or coastal marine waters of New York Bight (Table S2). The size range at the time of capture was 401 to 2050 mm fork length [FL], with a mean FL of 994.3 mm (SD = 230.3; Table 1). Monitoring efforts from 2011 through 2021 resulted in 845 unique observations of these fish on acoustic receiver arrays maintained throughout the OCS study site (Figures 1 and S3). The mean number of unique observations pooled by month was 202.8 (SD = 73.3), with the most unique observations in December (n = 348) and the least number of unique observations occurring in August (n = 83; Figure 1). In general, Atlantic Sturgeon were not observed as often during the late winter (i.e., January and February) or late summer (i.e., August and September) months, which could be associated with periods of reduced movement or limited monitoring efforts.

**Table 1.**
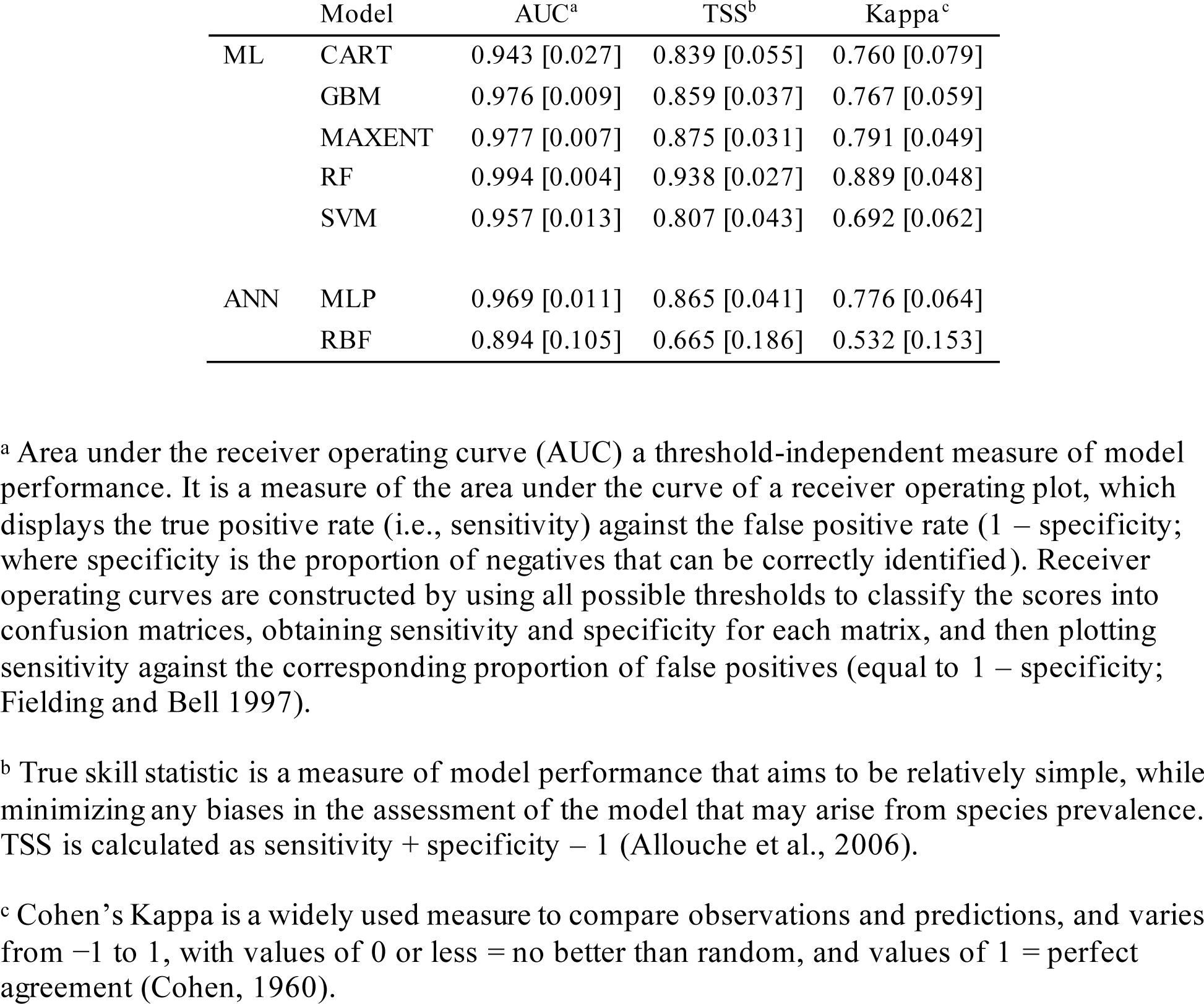
Overall mean [SD] model performance for all model types across all months. AUC = area under the receiver operating curve; TSS = true skill statistic; Kappa = Cohen’s kappa; ML = machine learning; ANN = artificial neural network; CART = classification and regression tree; GBM = gradient boosting machine; MAXENT = maximum entropy; RF = random forest; SVM = support vector machine; MLP = multilayer perceptron; RBF = radial basis function.

|  | Model | AUC<sup>a</sup> | TSS<sup>b</sup> | Kappa<sup>c</sup> |
| --- | --- | --- | --- | --- |
| ML | CART | 0.943 [0.027] | 0.839 [0.055] | 0.760 [0.079] |
|  | GBM | 0.976 [0.009] | 0.859 [0.037] | 0.767 [0.059] |
|  | MAXENT | 0.977 [0.007] | 0.875 [0.031] | 0.791 [0.049] |
|  | RF | 0.994 [0.004] | 0.938 [0.027] | 0.889 [0.048] |
|  | SVM | 0.957 [0.013] | 0.807 [0.043] | 0.692 [0.062] |
| ANN | MLP | 0.969 [0.011] | 0.865 [0.041] | 0.776 [0.064] |
|  | RBF | 0.894 [0.105] | 0.665 [0.186] | 0.532 [0.153] |

<sup>a</sup> Area under the receiver operating curve (AUC) a threshold-independent measure of model performance. It is a measure of the area under the curve of a receiver operating plot, which displays the true positive rate (i.e., sensitivity) against the false positive rate (1 – specificity; where specificity is the proportion of negatives that can be correctly identified). Receiver operating curves are constructed by using all possible thresholds to classify the scores into confusion matrices, obtaining sensitivity and specificity for each matrix, and then plotting sensitivity against the corresponding proportion of false positives (equal to 1 – specificity; Fielding and Bell 1997).
<sup>b</sup> True skill statistic is a measure of model performance that aims to be relatively simple, while minimizing any biases in the assessment of the model that may arise from species prevalence. TSS is calculated as sensitivity + specificity – 1 (Allouche et al., 2006).
<sup>c</sup> Cohen’s Kappa is a widely used measure to compare observations and predictions, and varies from –1 to 1, with values of 0 or less = no better than random, and values of 1 = perfect agreement (Cohen, 1960).

### SDM accuracy and importance of predictors

Model assessment metrics indicated similar trends in accuracy (i.e., ability to correctly classify presences or absences) between AI-model types (Table S6). Across all SDMs, RFs had the highest average performance (AUC [SD] = 0.99 [0.00]) and RBF had the lowest (AUC [SD] = 0.89 [0.11]). While RFs consistently performed the best, all algorithms had high accuracy and discriminating capacity across all metrics. The AUC of overall mean model performance across all months ranged from 0.89 to 0.99, suggesting high accuracy (i.e., RBF; 0.80 ≥ AUC < 0.90) or a near-perfect fit (i.e., CART, GBM, MAXENT, MLP, RF, and SVM; AUC ≥ 0.90; Table 1). TSS values of overall mean performance across all months ranged from 0.67 to 0.94, suggesting a high degree of non-randomness among predictions that were useful (i.e., RBF; 0.40 ≥ TSS < 0.80) or good to excellent (i.e., CART, GBM, MAXENT, MLP, RF, and SVM; TSS ≥ 0.80). Similarly, the Kappa index values of overall mean performance across all months ranged from 0.53 to 0.89, which indicated good (i.e., CART GBM, MAXENT, MLP, SVM, and RBF; 0.40 ≥ TSS < 0.80) or great model accuracy (i.e., RF; TSS ≥ 0.80).

Overall mean predictor importance from calculation of the Pearson correlation indicated that depth was the most important spatial predictor used by SDMs to predicts Atlantic Sturgeon occurrence across all months, followed in order by: SST, photoperiod, and sediment composition (Fig 3). However, minor variation in the order of predictor importance was identified between individual months (Table S7). While depth was typically identified as the predictor with the highest importance, SST was more important during August and photoperiod was more important during the months of October and November.

**Figure 2.**
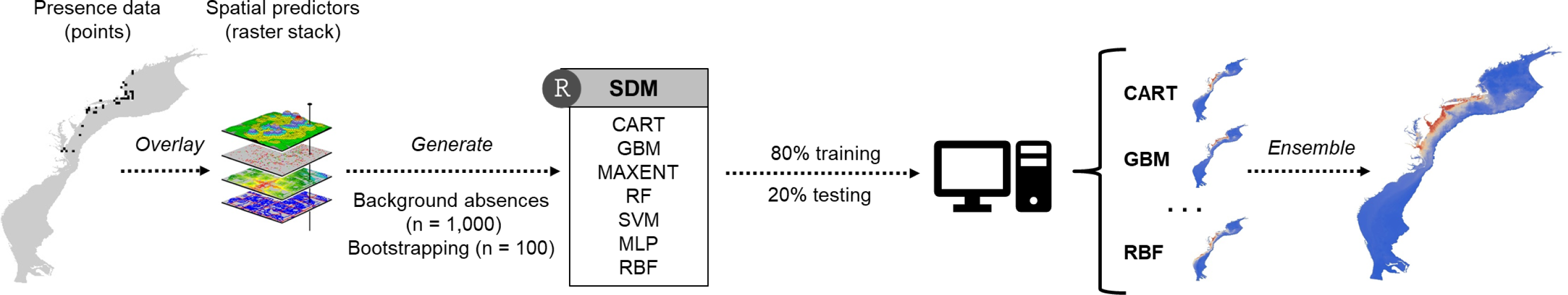
Species distribution model workflow for Atlantic Sturgeon occurring in the OCS study site. Acoustic telemetry detection records and uncorrelated spatial predictors were used to fit seven artificial intelligence SDMs for each month and a final monthly ensemble was generated.

**Figure 3.**
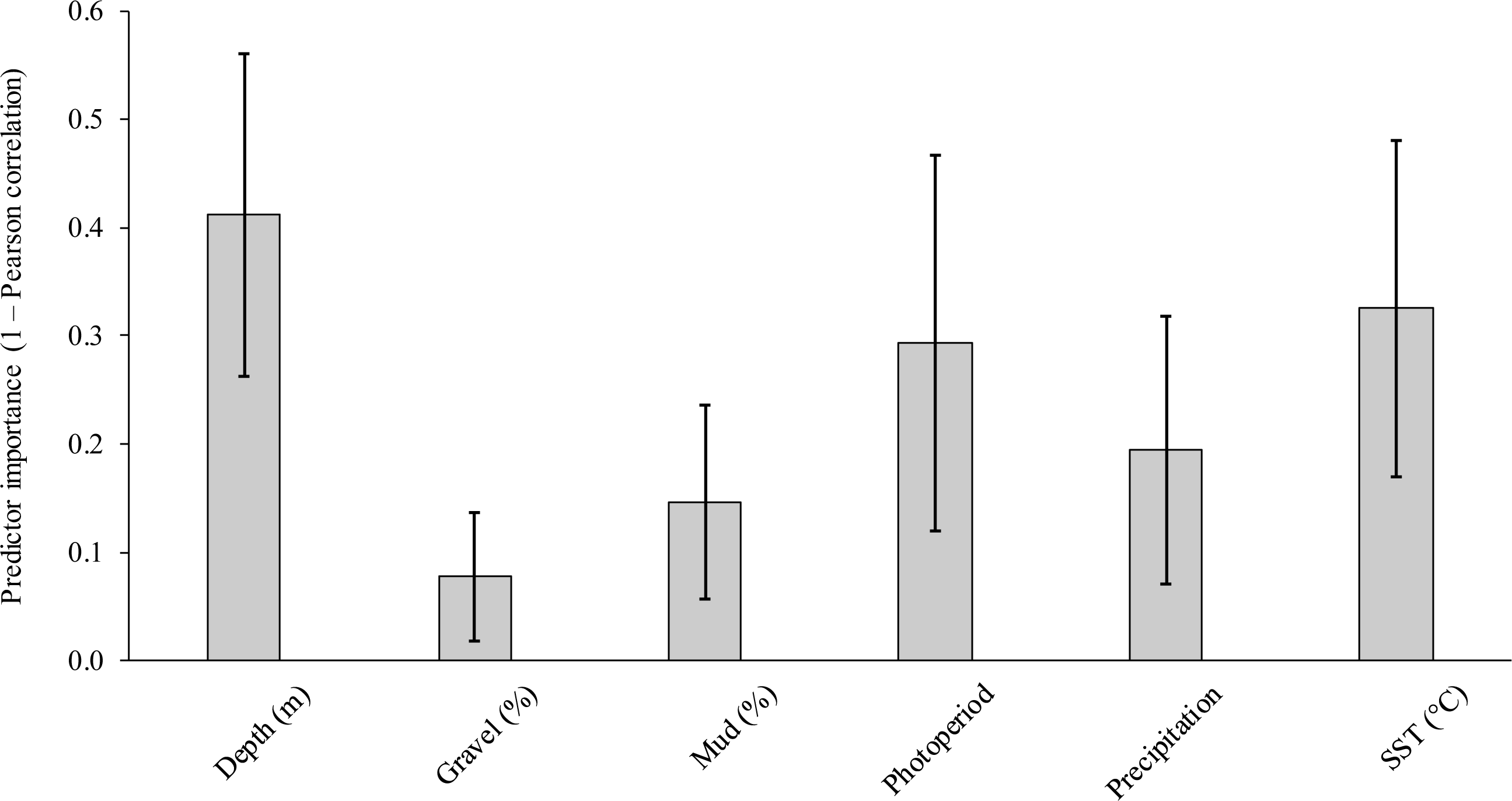
Overall mean predictor importance (1 – Pearson correlation ± 95% confidence interval) for Atlantic Sturgeon in the OCS study site across all months.

### Ensemble map products

All AI-algorithms met the high AUC threshold (i.e., AUC > 0.8; Table 6) and were retained for the final ensemble projections. Seasonal trends in Atlantic Sturgeon distribution were evident from monthly ensemble maps developed for the OCS study site (Figure 4). Atlantic Sturgeon were predicted to range across a large portion of the OCS study site, with expansion into more southernly habitats alongside increased occurrence in offshore waters during the months of November through March (Figures S1, S2, S3, S11, and S12). In contrast, the latitudinal distribution of Atlantic Sturgeon was predicted to contract to relatively nearshore habitats centered around putative aggregation sites and spawning rivers in the New York Bight during June through August (Figures S6, S7, S8). The spring (i.e., April–May; Figures S4 S5) and fall months (i.e., September–October; Figures S9 and S10) appeared to act as transitional periods as Atlantic Sturgeon move between these predicted extremes.

**Figure 4.**
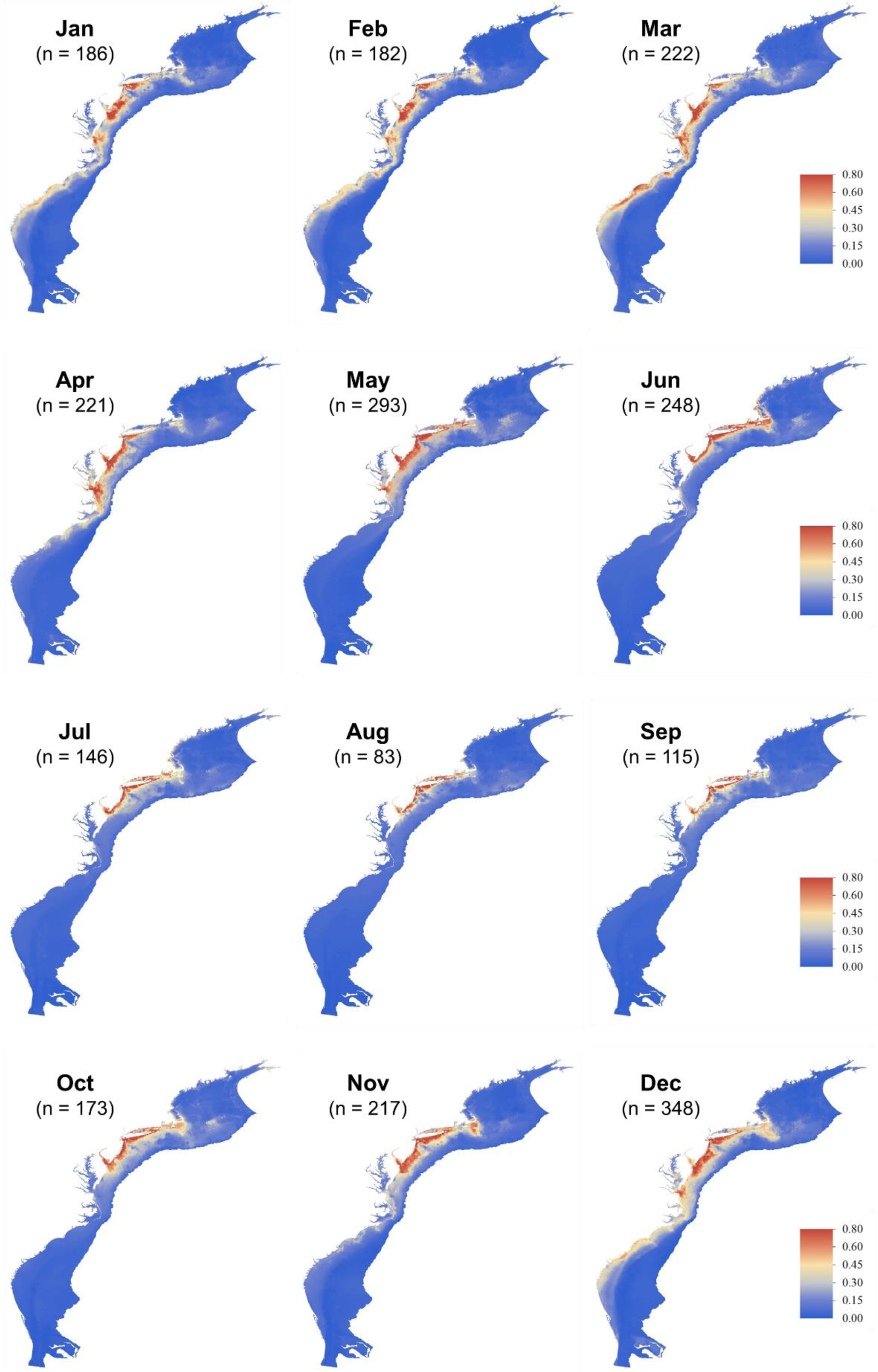
Monthly species distribution model ensemble maps of predicted Atlantic Sturgeon occurrence probability in the OCS study site at 1-km raster resolution. Counts of unique detection locations are indicated for each month.

Detailed map products indicated an increase in habitat overlap at wind energy sites during certain months. As distribution expanded along the coast and further offshore during the winter months, spatial overlap with wind energy was predicted to increase both within- and among development sites (Figures 5 and S13). While thresholds for the lowest predicted suitability value of occurrence were not identified, ensemble projections provided suitable detail at 1-km raster resolution to make monthly comparisons of occurrence at regional (e.g., New York Bight; Figure 5) and even site-specific scales (e.g., South Fork Wind Farm; Figure S13).

**Figure 5.**
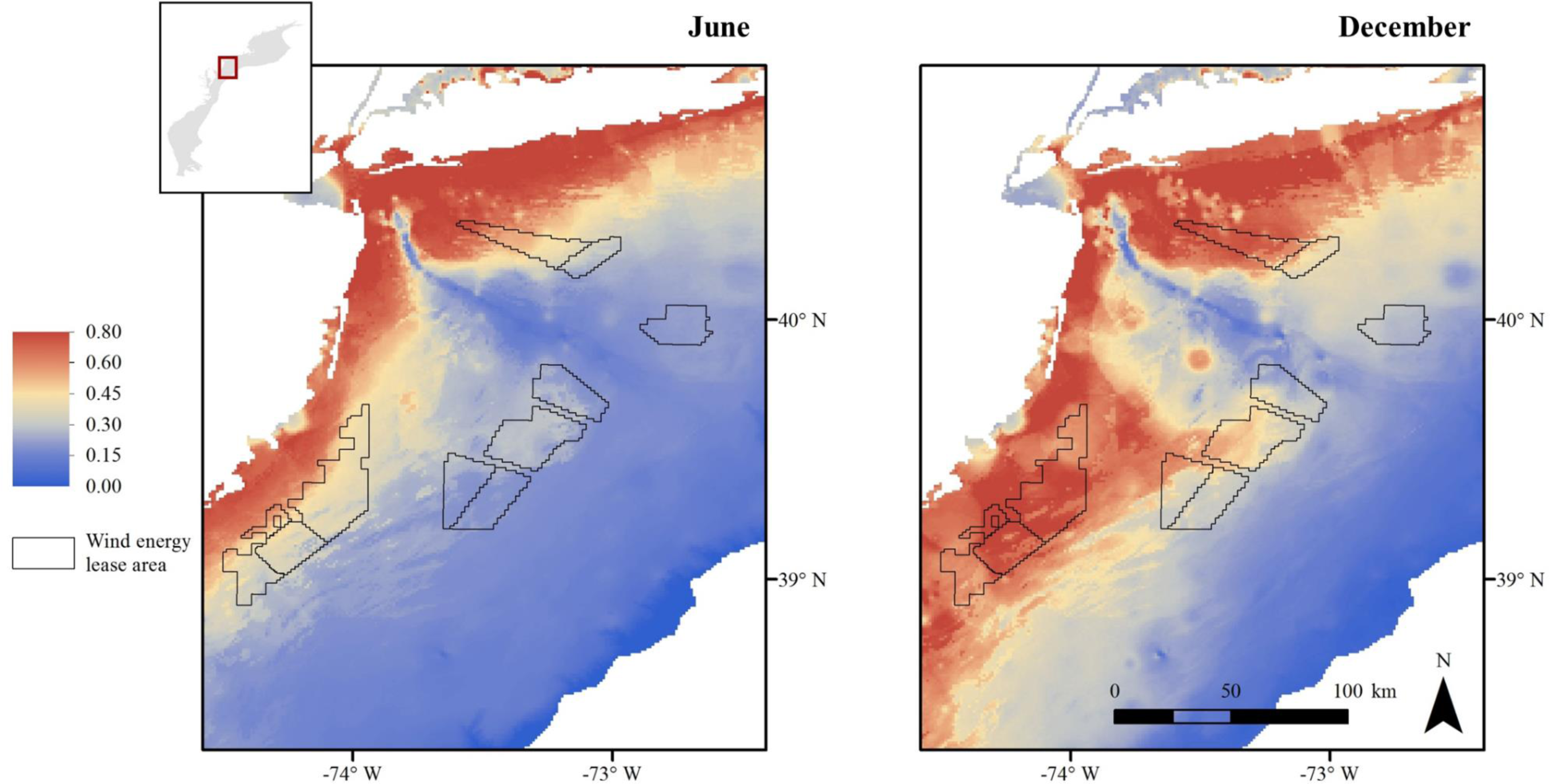
Predicted occurrence of Atlantic Sturgeon in wind energy lease sites located in the New York Bight off coastal New York and New Jersey. Comparative details are shown for species distribution model ensemble maps from June (left panel) and December (right panel) at 1-km raster resolution.

## Discussion

This study provides a novel approach to predicting species occurrence across a broad, largely unmonitored marine regions. The SDM framework presented combines artificial intelligence algorithms and ensemble methods to improve the predictive performance of SDMs while also providing the flexibility to leverage available data sources. With the development of offshore wind energy in the United States comes the recognized need to prioritize offshore research and monitoring effort in the context of ecological risk assessment and mitigation. The results of our case study suggest that predictive maps of rare or data-poor species can be readily constructed via SDM ensemble approaches and used to establish effective baseline criteria regarding spatial and temporal trends of Atlantic Sturgeon within the putative area of effect—an obligatory prerequisite for evaluating the impact of activities during all stages of offshore wind energy development. Ensemble maps products, when overlaid with the locations of offshore wind energy developments, can directly feed into the management of Atlantic Sturgeon in the Atlantic OCS at relevant coastwide, regional, and local scales.

The SDM framework demonstrated here offers the flexibility to leverage existing datasets or incorporate new data into existing models, allowing map products to be rapidly updated as new data become available. This framework is highly generalizable and can be readily applied to other species of interest or to investigate the extent of spatial overlap with ongoing or emergent threats. Because relevant environmental and spatial predictors may be shared among species or across communities (Noss, 1990; Belanger et al., 2012) there is also broad potential to streamline the generation of ensemble map products and allow for standardization across regions of interest with only slight modifications in input data used to inform the underlying model. A diversity of high quality, spatially referenced observations for input into SDMs are currently available from long-term monitoring datasets, including tagging programs, research surveys, museum records, and citizen science efforts. While potential issues with sample bias can occur because of the spatial variation of collection efforts (Boakes et al., 2010), sample size has been demonstrated to be more important than its spatial bias for predictive performance and modelling protocols exist to ensure data suitability (Boria et al., 2014; Fourcade et al., 2014; Shang et al., 2017; Gaul et al., 2020). As such, the identification and compilation of suitable data sources could provide an invaluable resource to expand this SDM framework to other regions or species but would require the support of potential stakeholders (including researchers, developers, and state federal governmental agencies) to maximize data sharing and prevent the duplication of modelling efforts.

Our case study applied telemetry data and AI algorithms within an SDM ensemble framework to reliably evaluate ecological range impacts at scales needed to inform management strategies for Atlantic Sturgeon across the entirety of the Atlantic OCS. Predictive monthly SDM ensemble maps were constructed using telemetry detections of Atlantic Sturgeon that were primarily captured at marine aggregation sites off New York State and subsequently detected throughout habitat in the OCS study site (i.e., Melnychuk et al., 2017; Ingram et al., 2019). Because of this, results from these models are likely skewed towards populations assigned to coastal rivers from the New York Bight distinct population segment (DPS, including the Hudson River in New York and the Delaware River in Delaware and New Jersey; USOFR, 2012). Although mixed aggregations of wide-ranging, coastwide genetic stocks of Atlantic Sturgeon can occur in marine waters, previous work suggest Hudson River fish contribute disproportionately to the composition of these aggregations in areas where sampling occurred (Dunton et al., 2012; Wirgin et al., 2015). As such, any application of these predictions to inform management for the overall coastwide population of Atlantic Sturgeon must proceed under the caveat that habitat use may be severely underestimated in some areas, particularly at the southern extent of the species’ range. Modelling efforts using spatially referenced observations from Atlantic Sturgeon of known-origin are suggested to develop population-specific SDMs across the species range. The incorporation of individuals from additional DPS locations would allow for development of stacked or joint SDMs (Zurell et al., 2020) and provide information on areas of overlap between populations—thus improving the identification of localized or regional threats to multiple stocks of Atlantic Sturgeon.

The excellent accuracy and high discriminatory capacity of AI algorithms in our case study provides strong impetus for their incorporation into future SDM applications. Overall results demonstrated that RF models consistently performed better and had higher accuracy compared to the other methods tested. The RF algorithm is an ensemble of classification or regression trees that combines the output of multiple decision trees to reach a single result (Evans et al., 2011). RFs are widely used in SDM applications because of their demonstrated flexibility and ability to provide better model fit for testing data relative to other methods—particularly when fitting models against varying data size or for under sampled study areas (e.g., Mi et al., 2017; Butler and Sanderson, 2022). Therefore, RFs are likely to outperform other methods when modelling the spatial distribution of data poor species such as Atlantic Sturgeon that would otherwise be limited by sample size. While model accuracy across methods would likely be improved by fine-tuning the settings used for parameter regularization, the use of default settings for these models is generally considered acceptable and allows for the standardization of methods across studies and ease of use (Phillips and Dudík, 2008). Regardless, the overall difference in model performance between the methods was relatively small and a weighted consensus of all seven AI algorithms was used to inform the final monthly ensemble models.

High-resolution occurrence maps of Atlantic Sturgeon generated from monthly SDM ensembles demonstrated an increase in spatial overlap at wind energy sites during certain months and seasons, with distribution biased toward shallow, nearshore regions during the summer months and expanding latitudinally into deeper offshore habitat during the winter. These trends were largely consistent with observations of Atlantic Sturgeon in marine waters from previous studies (e.g., Erickson et al., Ingram et al., 2019; Rothermel et al., 2020) and provide further indication that the potential negative impacts of wind energy development can be largely mitigated b y enacting spatial and temporal considerations during periods of increased incidence. Importantly, the application of SDM ensembles to inform spatial planning and monitoring efforts would provide managers with the discretion to prioritize among management needs through map comparisons at various scales. While shifts in seasonal distribution are easily discerned from coastwide maps, alternative scale options are needed to effectively identify management goals and develop action plans at the regional or state level (e.g., the New York Bight; see Figure 5). Similarly, the use of site-specific maps can inform management plans and policy settings for across all phases of a project’s development. A timely example is the South Fork Wind Farm (OCS-A 0517), located east of Long Island in New York, which has been operational at commercial scale (i.e., 12 turbines; ∼132 MW) since December 2023. Detailed estimates of Atlantic Sturgeon occurrence were not available to inform required statutory consultations during the planning and construction phase of development at the site. However, retrospective SDM ensemble maps show moderate probability of seasonal resource overlap in the turbine array and high predicted probability of spatial overlap across all months along the path of subsea power (export) cables installed in state waters (< 3 nm from shore; see Figure S13). Despite limited data on impacts of electromagnetic fields associated with electricity production (Hutchison et al., 2020; Backstrom and Warden, 2023), the need for spatially explicit monitoring efforts during export cable installation would have been evident from the application of SDMs during the early phases of marine spatial planning.

Modelling efforts used a relatively small number of environmental predictors (n =6) to fit AI SDMs with high accuracy. The influence of these environmental factors on Atlantic Sturgeon occurrence suggests that broadscale, seasonal shifts in distribution and incidence in offshore development sites are associated with predictable spatial and temporal variation in abiotic conditions. Variation in the order of predictor importance may indicate the role of clinal variations in abiotic conditions on temporal distribution and habitat selection, particularly during periods of increased migration (Ingram et al., 2019). Atlantic Sturgeon undertake extensive coastal migrations, ostensibly to exploit favorable clinal gradients and temporally predictable foraging habitat (Gross et al., 1988; Dingle and Drake, 2007). While preferential distribution in shallow, near shore marine waters above sand or mud substrates may indicate high prey density (Stein et al., 2004b; Johnson et al., 1997), a clear linkage between foraging behavior and resource utilization in marine waters has yet to be determined. Photoperiod and water temperature are recognized as factors that modulate migratory strategies in other sturgeon species (Cech and Doroshov, 2004; Papoulious et al., 2011). These predictors may act as reliable, long-term triggers of migration or cues regarding localized conditions, and have been associated with the predictable seasonal movements of Atlantic Sturgeon between shallow, coastal waters and offshore habitat (Ingram et al., 2019).

The predicted occurrence of Atlantic Sturgeon in offshore waters of Atlantic OCS underscores the importance of reliable distribution maps to inform monitoring and management priorities for marine spatial planning, particularly regarding data poor species that are of conservation or commercial interest. While considerable research and management attention has focused on the potential environmental impacts of offshore wind development, empirical evaluations for this and many other species of concern are presently inadequate and limited by a lack of basic knowledge regarding spatial overlap across marine regions. The characterization of broad spatiotemporal trends in distribution and habitat use in offshore waters are necessary to better define monitoring parameters and guide threat assessments in marine wind energy areas. Our results indicate that predictive maps of species distribution can be readily generated using AI-derived SDM ensemble. These methods are highly generalizable and readily applied and generate map products that are scalable to management needs. This approach will help inform development of conservation strategies and the quantitative evaluation in marine waters at scales relevant to ongoing development activities

## Supporting Information

**Table S1.**
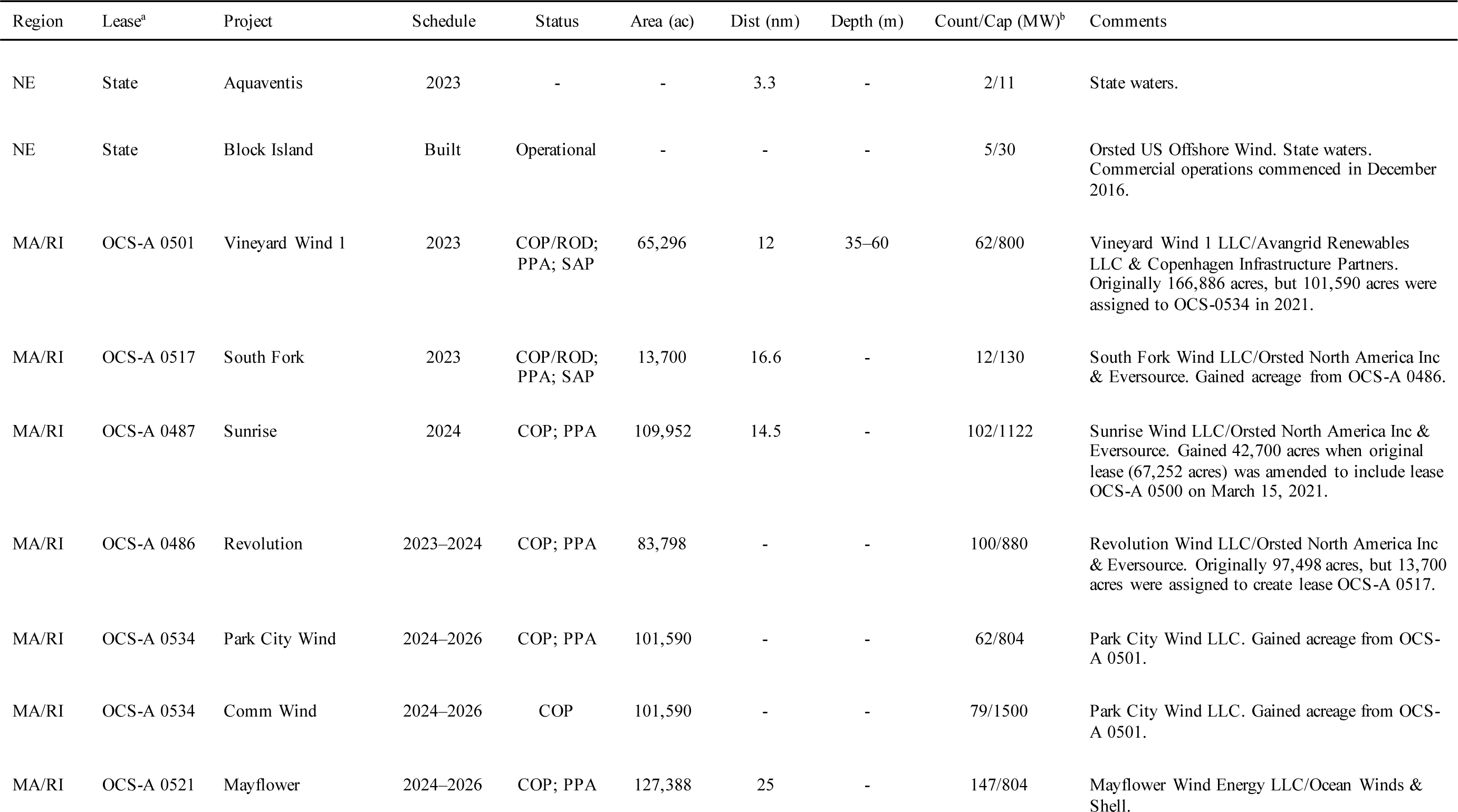
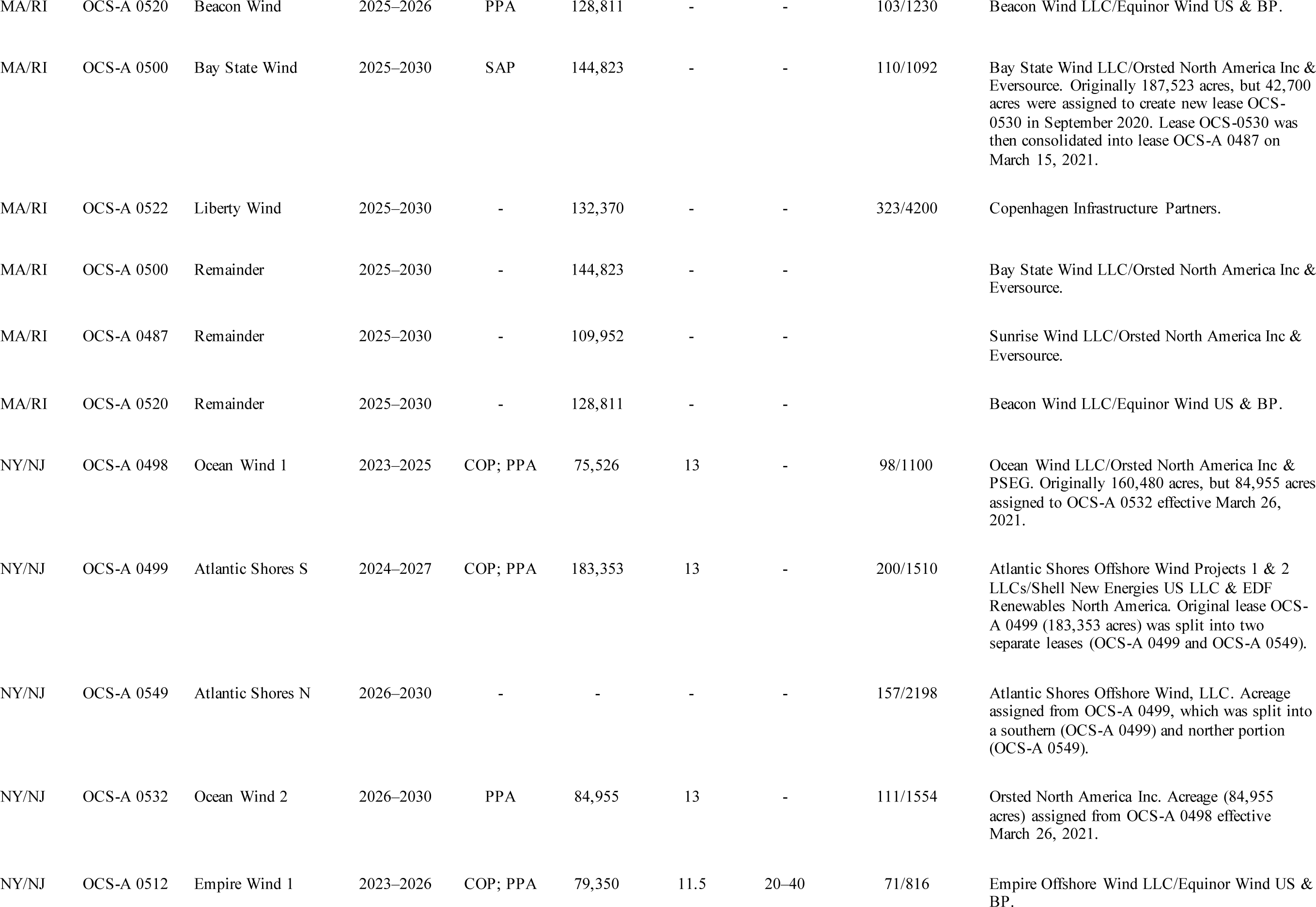
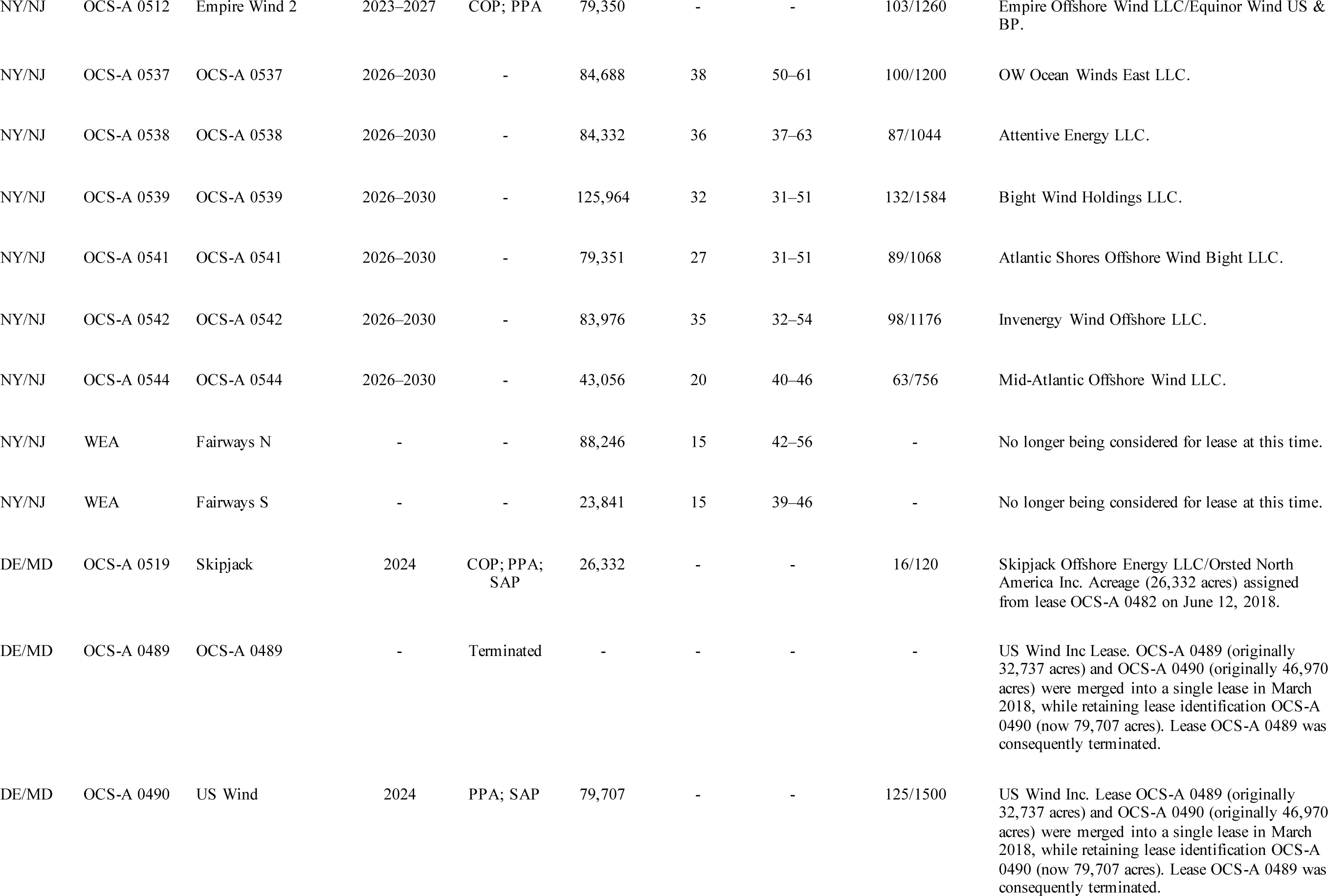
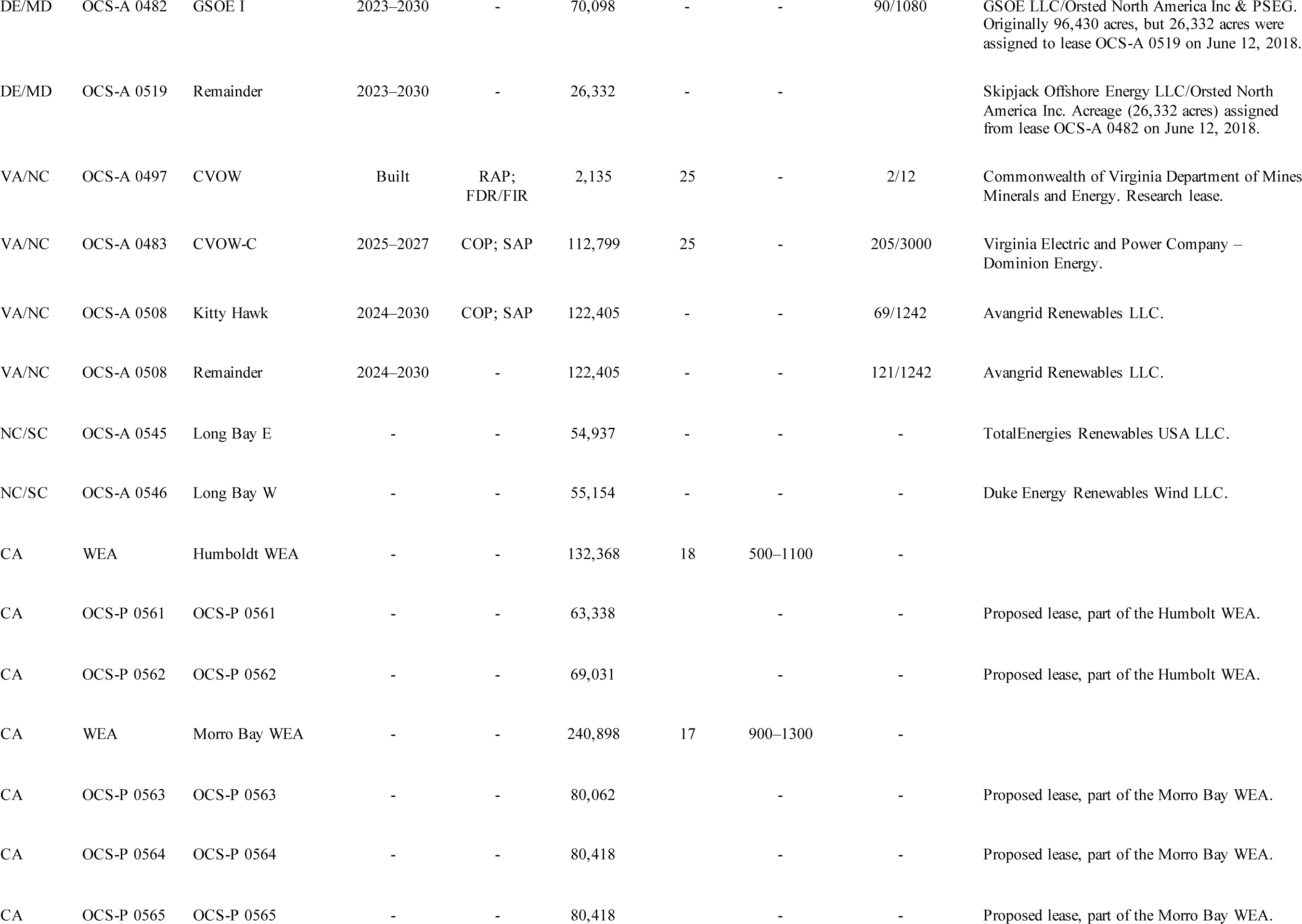
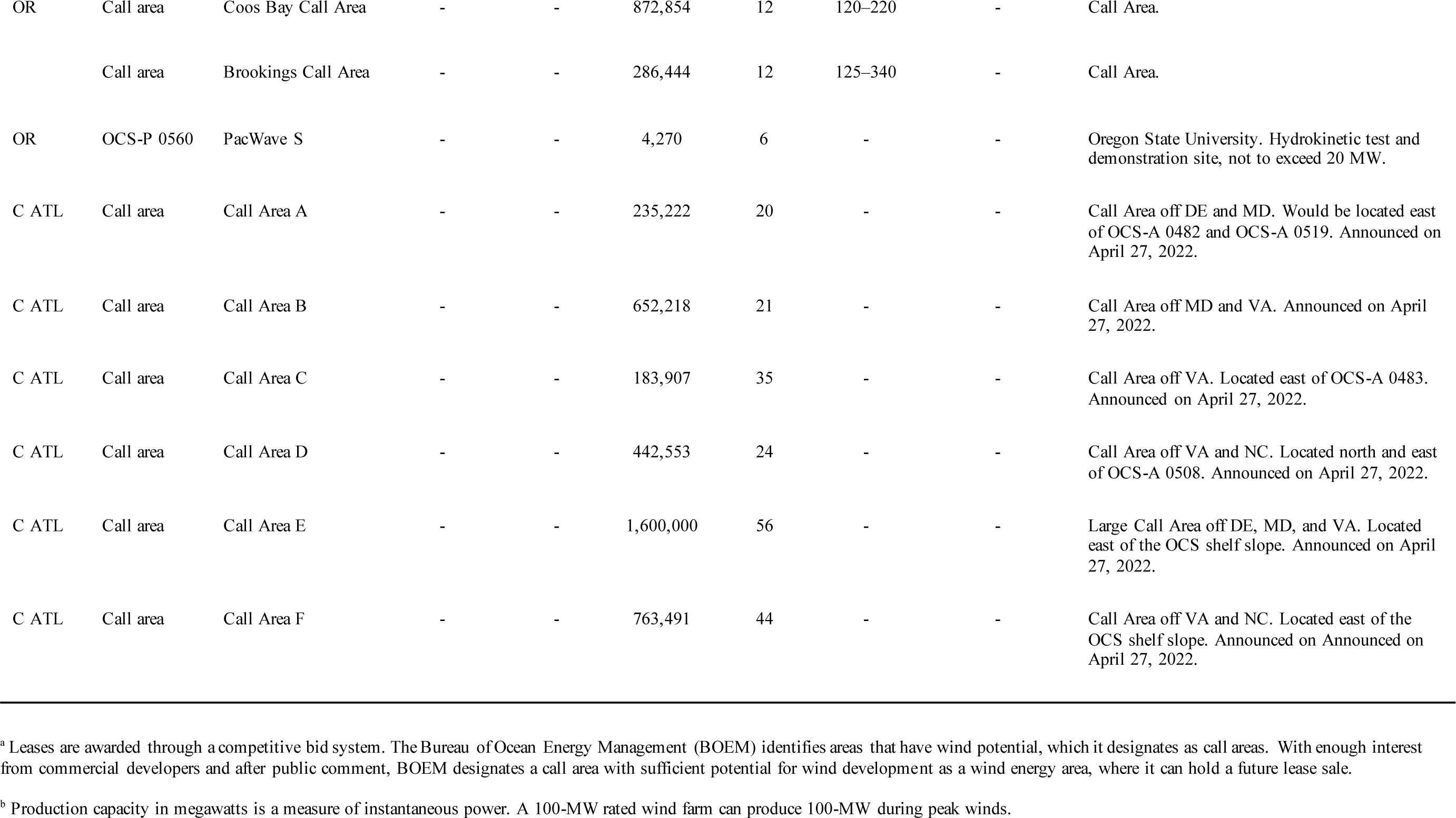
Offshore wind energy in the United States. Information is current as of July 21, 2022. COP = construction and operations plan; COP/ROD = construction and operations plan approved, and record of decision issued; PPA = power purchase agreement; SAP = site assessment plan; RAP = research activities plan; FRD/FIR = f acility design and fabrication and installation reports.

| Region | Lease<sup>a</sup> | Project | Schedule | Status | Area (ac) | Dist (nm) | Depth (m) | Count/Cap (MW)<sup>b</sup> | Comments |
| --- | --- | --- | --- | --- | --- | --- | --- | --- | --- |
| NE | State | Aquaventis | 2023 | - | - | 3.3 | - | 2/11 | State waters. |
| NE | State | Block Island | Built | Operational | - | - | - | 5/30 | Orsted US Offshore Wind. State waters. Commercial operations commenced in December 2016. |
| MA/RI | OCS-A 0501 | Vineyard Wind 1 | 2023 | COP/ROD; PPA; SAP | 65,296 | 12 | 35–60 | 62/800 | Vineyard Wind 1 LLC/Avangrid Renewables LLC & Copenhagen Infrastructure Partners. Originally 166,886 acres, but 101,590 acres were assigned to OCS-0534 in 2021. |
| MA/RI | OCS-A 0517 | South Fork | 2023 | COP/ROD; PPA; SAP | 13,700 | 16.6 | - | 12/130 | South Fork Wind LLC/Orsted North America Inc & Eversource. Gained acreage from OCS-A 0486. |
| MA/RI | OCS-A 0487 | Sunrise | 2024 | COP; PPA | 109,952 | 14.5 | - | 102/1122 | Sunrise Wind LLC/Orsted North America Inc & Eversource. Gained 42,700 acres when original lease (67,252 acres) was amended to include lease OCS-A 0500 on March 15, 2021. |
| MA/RI | OCS-A 0486 | Revolution | 2023–2024 | COP; PPA | 83,798 | - | - | 100/880 | Revolution Wind LLC/Orsted North America Inc & Eversource. Originally 97,498 acres, but 13,700 acres were assigned to create lease OCS-A 0517. |
| MA/RI | OCS-A 0534 | Park City Wind | 2024–2026 | COP; PPA | 101,590 | - | - | 62/804 | Park City Wind LLC. Gained acreage from OCS-A 0501. |
| MA/RI | OCS-A 0534 | Comm Wind | 2024–2026 | COP | 101,590 | - | - | 79/1500 | Park City Wind LLC. Gained acreage from OCS-A 0501. |
| MA/RI | OCS-A 0521 | Mayflower | 2024–2026 | COP; PPA | 127,388 | 25 | - | 147/804 | Mayflower Wind Energy LLC/Ocean Winds & Shell. |
| MA/RI | OCS-A 0520 | Beacon Wind | 2025–2026 | PPA | 128,811 | - | - | 103/1230 | Beacon Wind LLC/Equinor Wind US & BP. |
| MA/RI | OCS-A 0500 | Bay State Wind | 2025–2030 | SAP | 144,823 | - | - | 110/1092 | Bay State Wind LLC/Orsted North America Inc & Eversource. Originally 187,523 acres, but 42,700 acres were assigned to create new lease OCS-0530 in September 2020. Lease OCS-0530 was then consolidated into lease OCS-A 0487 on March 15, 2021. |
| MA/RI | OCS-A 0522 | Liberty Wind | 2025–2030 | - | 132,370 | - | - | 323/4200 | Copenhagen Infrastructure Partners. |
| MA/RI | OCS-A 0500 | Remainder | 2025–2030 | - | 144,823 | - | - |  | Bay State Wind LLC/Orsted North America Inc & Eversource. |
| MA/RI | OCS-A 0487 | Remainder | 2025–2030 | - | 109,952 | - | - |  | Sunrise Wind LLC/Orsted North America Inc & Eversource. |
| MA/RI | OCS-A 0520 | Remainder | 2025–2030 | - | 128,811 | - | - |  | Beacon Wind LLC/Equinor Wind US & BP. |
| NY/NJ | OCS-A 0498 | Ocean Wind 1 | 2023–2025 | COP; PPA | 75,526 | 13 | - | 98/1100 | Ocean Wind LLC/Orsted North America Inc & PSEG. Originally 160,480 acres, but 84,955 acres assigned to OCS-A 0532 effective March 26, 2021. |
| NY/NJ | OCS-A 0499 | Atlantic Shores S | 2024–2027 | COP; PPA | 183,353 | 13 | - | 200/1510 | Atlantic Shores Offshore Wind Projects 1 & 2 LLCs/Shell New Energies US LLC & EDF Renewables North America. Original lease OCS-A 0499 (183,353 acres) was split into two separate leases (OCS-A 0499 and OCS-A 0549). |
| NY/NJ | OCS-A 0549 | Atlantic Shores N | 2026–2030 | - | - | - | - | 157/2198 | Atlantic Shores Offshore Wind, LLC. Acreage assigned from OCS-A 0499, which was split into a southern (OCS-A 0499) and norther portion (OCS-A 0549). |
| NY/NJ | OCS-A 0532 | Ocean Wind 2 | 2026–2030 | PPA | 84,955 | 13 | - | 111/1554 | Orsted North America Inc. Acreage (84,955 acres) assigned from OCS-A 0498 effective March 26, 2021. |
| NY/NJ | OCS-A 0512 | Empire Wind 1 | 2023–2026 | COP; PPA | 79,350 | 11.5 | 20–40 | 71/816 | Empire Offshore Wind LLC/Equinor Wind US & BP. |
| NY/NJ | OCS-A 0512 | Empire Wind 2 | 2023–2027 | COP; PPA | 79,350 | - | - | 103/1260 | Empire Offshore Wind LLC/Equinor Wind US & BP. |
| NY/NJ | OCS-A 0537 | OCS-A 0537 | 2026–2030 | - | 84,688 | 38 | 50–61 | 100/1200 | OW Ocean Winds East LLC. |
| NY/NJ | OCS-A 0538 | OCS-A 0538 | 2026–2030 | - | 84,332 | 36 | 37–63 | 87/1044 | Attentive Energy LLC. |
| NY/NJ | OCS-A 0539 | OCS-A 0539 | 2026–2030 | - | 125,964 | 32 | 31–51 | 132/1584 | Bight Wind Holdings LLC. |
| NY/NJ | OCS-A 0541 | OCS-A 0541 | 2026–2030 | - | 79,351 | 27 | 31–51 | 89/1068 | Atlantic Shores Offshore Wind Bight LLC. |
| NY/NJ | OCS-A 0542 | OCS-A 0542 | 2026–2030 | - | 83,976 | 35 | 32–54 | 98/1176 | Invenergy Wind Offshore LLC. |
| NY/NJ | OCS-A 0544 | OCS-A 0544 | 2026–2030 | - | 43,056 | 20 | 40–46 | 63/756 | Mid-Atlantic Offshore Wind LLC. |
| NY/NJ | WEA | Fairways N | - | - | 88,246 | 15 | 42–56 | - | No longer being considered for lease at this time. |
| NY/NJ | WEA | Fairways S | - | - | 23,841 | 15 | 39–46 | - | No longer being considered for lease at this time. |
| DE/MD | OCS-A 0519 | Skipjack | 2024 | COP; PPA; SAP | 26,332 | - | - | 16/120 | Skipjack Offshore Energy LLC/Orsted North America Inc. Acreage (26,332 acres) assigned from lease OCS-A 0482 on June 12, 2018. |
| DE/MD | OCS-A 0489 | OCS-A 0489 | - | Terminated | - | - | - | - | US Wind Inc Lease. OCS-A 0489 (originally 32,737 acres) and OCS-A 0490 (originally 46,970 acres) were merged into a single lease in March 2018, while retaining lease identification OCS-A 0490 (now 79,707 acres). Lease OCS-A 0489 was consequently terminated. |
| DE/MD | OCS-A 0490 | US Wind | 2024 | PPA; SAP | 79,707 | - | - | 125/1500 | US Wind Inc. Lease OCS-A 0489 (originally 32,737 acres) and OCS-A 0490 (originally 46,970 acres) were merged into a single lease in March 2018, while retaining lease identification OCS-A 0490 (now 79,707 acres). Lease OCS-A 0489 was consequently terminated. |
| DE/MD | OCS-A 0482 | GSOE I | 2023–2030 | - | 70,098 | - | - | 90/1080 | GSOE LLC/Orsted North America Inc & PSEG. Originally 96,430 acres, but 26,332 acres were assigned to lease OCS-A 0519 on June 12, 2018. |
| DE/MD | OCS-A 0519 | Remainder | 2023–2030 | - | 26,332 | - | - |  | Skipjack Offshore Energy LLC/Orsted North America Inc. Acreage (26,332 acres) assigned from lease OCS-A 0482 on June 12, 2018. |
| VA/NC | OCS-A 0497 | CVOW | Built | RAP;<br>FDR/FIR | 2,135 | 25 | - | 2/12 | Commonwealth of Virginia Department of Mines Minerals and Energy. Research lease. |
| VA/NC | OCS-A 0483 | CVOW-C | 2025–2027 | COP; SAP | 112,799 | 25 | - | 205/3000 | Virginia Electric and Power Company – Dominion Energy. |
| VA/NC | OCS-A 0508 | Kitty Hawk | 2024–2030 | COP; SAP | 122,405 | - | - | 69/1242 | Avangrid Renewables LLC. |
| VA/NC | OCS-A 0508 | Remainder | 2024–2030 | - | 122,405 | - | - | 121/1242 | Avangrid Renewables LLC. |
| NC/SC | OCS-A 0545 | Long Bay E | - | - | 54,937 | - | - | - | TotalEnergies Renewables USA LLC. |
| NC/SC | OCS-A 0546 | Long Bay W | - | - | 55,154 | - | - | - | Duke Energy Renewables Wind LLC. |
| CA | WEA | Humboldt WEA | - | - | 132,368 | 18 | 500–1100 | - |  |
| CA | OCS-P 0561 | OCS-P 0561 | - | - | 63,338 |  | - | - | Proposed lease, part of the Humboldt WEA. |
| CA | OCS-P 0562 | OCS-P 0562 | - | - | 69,031 |  | - | - | Proposed lease, part of the Humboldt WEA. |
| CA | WEA | Morro Bay WEA | - | - | 240,898 | 17 | 900–1300 | - |  |
| CA | OCS-P 0563 | OCS-P 0563 | - | - | 80,062 |  | - | - | Proposed lease, part of the Morro Bay WEA. |
| CA | OCS-P 0564 | OCS-P 0564 | - | - | 80,418 |  | - | - | Proposed lease, part of the Morro Bay WEA. |
| CA | OCS-P 0565 | OCS-P 0565 | - | - | 80,418 |  | - | - | Proposed lease, part of the Morro Bay WEA. |
| OR | Call area | Coos Bay Call Area | - | - | 872,854 | 12 | 120–220 | - | Call Area. |
|  | Call area | Brookings Call Area | - | - | 286,444 | 12 | 125–340 | - | Call Area. |
| OR | OCS-P 0560 | PacWave S | - | - | 4,270 | 6 | - | - | Oregon State University. Hydrokinetic test and demonstration site, not to exceed 20 MW. |
| C ATL | Call area | Call Area A | - | - | 235,222 | 20 | - | - | Call Area off DE and MD. Would be located east of OCS-A 0482 and OCS-A 0519. Announced on April 27, 2022. |
| C ATL | Call area | Call Area B | - | - | 652,218 | 21 | - | - | Call Area off MD and VA. Announced on April 27, 2022. |
| C ATL | Call area | Call Area C | - | - | 183,907 | 35 | - | - | Call Area off VA. Located east of OCS-A 0483. Announced on April 27, 2022. |
| C ATL | Call area | Call Area D | - | - | 442,553 | 24 | - | - | Call Area off VA and NC. Located north and east of OCS-A 0508. Announced on April 27, 2022. |
| C ATL | Call area | Call Area E | - | - | 1,600,000 | 56 | - | - | Large Call Area off DE, MD, and VA. Located east of the OCS shelf slope. Announced on April 27, 2022. |
| C ATL | Call area | Call Area F | - | - | 763,491 | 44 | - | - | Call Area off VA and NC. Located east of the OCS shelf slope. Announced on Announced on April 27, 2022. |

<sup>a</sup> Leases are awarded through a competitive bid system. The Bureau of Ocean Energy Management (BOEM) identifies areas that have wind potential, which it designates as call areas. With enough interest from commercial developers and after public comment, BOEM designates a call area with sufficient potential for wind development as a wind energy area, where it can hold a future lease sale.
<sup>b</sup> Production capacity in megawatts is a measure of instantaneous power. A 100-MW rated wind farm can produce 100-MW during peak winds.

**Table S2.** Summary of biological and tagging data from Atlantic Sturgeon (*Acipenser oxyrinchus*; n = 599) used in this study. FL = fork length; TL = total length; SBU = Stony Brook University; DSU = Delaware State University; MU = Monmouth University.

| Date | Fish | Acoustic tag | USFWS ID | FL (mm) | TL (mm) | Weight (kg) | Organization |
| --- | --- | --- | --- | --- | --- | --- | --- |
| 2010-10-23 | ATS-001 | A69-9001-1140 | 404100010 | 1040 | 1200 | 9.42 | SBU |
| 2010-10-25 | ATS-002 | A69-9001-1071 | 404100008 | 1290 | 1450 | 17.84 | SBU |
| 2011-04-13 | ATS-003 | A69-9001-1075 | - | 745 | 894 | 2.80 | SBU |
| 2011-04-14 | ATS-004 | A69-9001-1079 | - | 642 | 753 | 1.80 | MU |
| 2011-04-26 | ATS-005 | A69-9001-1068 | - | 678 | 802 | 2.30 | SBU |
| 2011-04-26 | ATS-006 | A69-9001-1070 | - | 777 | 910 | 4.80 | SBU |
| 2011-04-27 | ATS-007 | A69-9001-1074 | - | 742 | 838 | 3.50 | SBU |
| 2011-05-24 | ATS-008 | A69-9001-32461 | 355110045 | - | - | - | SBU |
| 2011-05-24 | ATS-009 | A69-9001-32460 | 355110056 | 767 | 903 | 3.62 | SBU |
| 2011-05-24 | ATS-010 | A69-9001-32473 | 355110037 | 824 | 965 | 4.40 | SBU |
| 2011-05-24 | ATS-011 | A69-9001-32445 | 355110071 | 784 | 891 | 4.56 | SBU |
| 2011-05-24 | ATS-012 | A69-9001-32452 | 355110064 | 917 | 1070 | 7.28 | SBU |
| 2011-05-24 | ATS-013 | A69-9001-1094 | 355110051 | 965 | 1089 | 7.76 | SBU |
| 2011-05-24 | ATS-014 | A69-9001-1088 | 355110005 | 983 | 1149 | 8.87 | SBU |
| 2011-05-24 | ATS-015 | A69-9001-32459 | 355110057 | 985 | 1129 | 9.01 | SBU |
| 2011-05-24 | ATS-016 | A69-9001-32453 | 355110063 | 971 | 1108 | 9.09 | SBU |
| 2011-05-24 | ATS-017 | A69-9001-1097 | 355110029 | 1027 | 1195 | 9.84 | SBU |
| 2011-05-24 | ATS-018 | A69-9001-32446 | 355110070 | 1054 | 1225 | 9.84 | SBU |
| 2011-05-24 | ATS-019 | A69-9001-1086 | 355110007 | 1030 | 1196 | 9.87 | SBU |
| 2011-05-24 | ATS-020 | A69-9001-32457 | 355110059 | 1040 | 1191 | 10.11 | SBU |
| 2011-05-24 | ATS-021 | A69-9001-1091 | 355110006 | 1042 | 1206 | 10.50 | SBU |
| 2011-05-24 | ATS-022 | A69-9001-32454 | 355110062 | 1068 | 1253 | 12.23 | SBU |
| 2011-05-24 | ATS-023 | A69-9001-1087 | 355110032 | 1053 | 1209 | 12.54 | SBU |
| 2011-05-24 | ATS-024 | A69-9001-32469 | 355110053 | 1084 | 1250 | 12.62 | SBU |
| 2011-05-24 | ATS-025 | A69-9001-1105 | 355110020 | 1089 | 1276 | 12.81 | SBU |
| 2011-05-24 | ATS-026 | A69-9001-32471 | 355110039 | 1167 | 1254 | 12.94 | SBU |
| 2011-05-24 | ATS-027 | A69-9001-32447 | 355110068 | 1067 | 1234 | 13.04 | SBU |
| 2011-05-24 | ATS-028 | A69-9001-1102 | 355110027 | 1110 | 1283 | 13.22 | SBU |
| 2011-05-24 | ATS-029 | A69-9001-1110 | 355110019 | 1147 | 1290 | 13.27 | SBU |
| 2011-05-24 | ATS-030 | A69-9001-32444 | 355110073 | 1140 | 1299 | 13.62 | SBU |
| 2011-05-24 | ATS-031 | A69-9001-1092 | 355110050 | 1122 | 1321 | 13.88 | SBU |
| 2011-05-24 | ATS-032 | A69-9001-32472 | 355110038 | 1149 | 1342 | 14.12 | SBU |
| 2011-05-24 | ATS-033 | A69-9001-32451 | 355110066 | 1103 | 1266 | 14.41 | SBU |
| 2011-05-24 | ATS-034 | A69-9001-1106 | 355110024 | 1129 | 1334 | 14.64 | SBU |
| 2011-05-24 | ATS-035 | A69-9001-1093 | 355110046 | 1182 | 1353 | 15.22 | SBU |
| 2011-05-24 | ATS-036 | A69-9001-32465 | 355110042 | 1114 | 1281 | 15.75 | SBU |
| 2011-05-24 | ATS-037 | A69-9001-1108 | 355110022 | 1172 | 1357 | 16.06 | SBU |
| 2011-05-24 | ATS-038 | A69-9001-32474 | 355110009 | 1224 | 1433 | 16.98 | SBU |
| 2011-05-24 | ATS-039 | A69-9001-32463 | 355110044 | 1220 | 1430 | 17.09 | SBU |
| 2011-05-24 | ATS-040 | A69-9001-32477 | 355110012 | 1213 | 1392 | 17.14 | SBU |
| 2011-05-24 | ATS-041 | A69-9001-32470 | 355110054 | 1232 | 1397 | 17.64 | SBU |
| 2011-05-24 | ATS-042 | A69-9001-32476 | 355110011 | 1207 | 1433 | 17.71 | SBU |
| 2011-05-24 | ATS-043 | A69-9001-32467 | 355110041 | 1203 | 1395 | 17.81 | SBU |
| 2011-05-24 | ATS-044 | A69-9001-32456 | 355110060 | 1223 | 1403 | 18.61 | SBU |
| 2011-05-24 | ATS-045 | A69-9001-1096 | 355110048 | 1239 | 1436 | 18.69 | SBU |
| 2011-05-24 | ATS-046 | A69-9001-32458 | 355110058 | 1208 | 1401 | 19.33 | SBU |
| 2011-05-24 | ATS-047 | A69-9001-1103 | 355110023 | 1246 | 1433 | 19.48 | SBU |
| 2011-05-24 | ATS-048 | A69-9001-1115 | 355110013 | 1248 | 1452 | 19.95 | SBU |
| 2011-05-24 | ATS-049 | A69-9001-32448 | 355110069 | 1259 | 1445 | 20.76 | SBU |
| 2011-05-24 | ATS-050 | A69-9001-1111 | 355110017 | 1260 | 1463 | 21.00 | SBU |
| 2011-05-24 | ATS-051 | A69-9001-1113 | 355110014 | 1280 | 1482 | 21.11 | SBU |
| 2011-05-24 | ATS-052 | A69-9001-32455 | 355110061 | 1343 | 1560 | 21.85 | SBU |
| 2011-05-24 | ATS-053 | A69-9001-1089 | 355110031 | 1353 | 1670 | 26.39 | SBU |
| 2011-05-24 | ATS-054 | A69-9001-1100 | 355110028 | 1454 | 1723 | 27.96 | SBU |
| 2011-05-24 | ATS-055 | A69-9001-1095 | 355110047 | 1510 | 1739 | 29.32 | SBU |
| 2011-05-24 | ATS-056 | A69-9001-1107 | 355110018 | 1472 | 1698 | 29.71 | SBU |
| 2011-05-24 | ATS-057 | A69-9001-32449 | 355110067 | 1550 | 1768 | 32.64 | SBU |
| 2011-10-12 | ATS-058 | A69-9001-30333 | 355110080 | 1193 | 1390 | 13.01 | SBU |
| 2011-10-12 | ATS-059 | A69-9001-30331 | 654080038 | 1238 | 1395 | 16.02 | SBU |
| 2011-10-13 | ATS-060 | A69-9001-30330 | 355110082 | 1210 | 1380 | 15.99 | SBU |
| 2011-10-24 | ATS-061 | A69-9001-30182 | 404110002 | 1335 | 1554 | 21.50 | SBU |
| 2011-10-25 | ATS-062 | A69-9001-30183 | 404110003 | 1035 | 1223 | 8.70 | SBU |
| 2011-10-31 | ATS-063 | A69-9001-30329 | 355110100 | 737 | 872 | 3.50 | SBU |
| 2011-10-31 | ATS-064 | A69-9001-30332 | 355110090 | 805 | 947 | 3.90 | SBU |
| 2011-10-31 | ATS-065 | A69-9001-30200 | 355110105 | 785 | 921 | 4.00 | SBU |
| 2011-10-31 | ATS-066 | A69-9001-30211 | 355110116 | 798 | 941 | 4.50 | SBU |
| 2011-10-31 | ATS-067 | A69-9001-30210 | 355110108 | 852 | 992 | 4.90 | SBU |
| 2011-10-31 | ATS-068 | A69-9001-30206 | 355110110 | 811 | 969 | 5.10 | SBU |
| 2011-10-31 | ATS-069 | A69-9001-30222 | 355110128 | 960 | 1130 | 5.80 | SBU |
| 2011-10-31 | ATS-070 | A69-9001-30321 | 355110084 | 935 | 1095 | 6.30 | SBU |
| 2011-10-31 | ATS-071 | A69-9001-30324 | 355110099 | 915 | 1078 | 6.40 | SBU |
| 2011-10-31 | ATS-072 | A69-9001-30205 | 355110113 | 940 | 1072 | 6.70 | SBU |
| 2011-10-31 | ATS-073 | A69-9001-30195 | 355110141 | 920 | 1070 | 6.80 | SBU |
| 2011-10-31 | ATS-074 | A69-9001-30203 | 355110138 | 965 | 1130 | 6.90 | SBU |
| 2011-10-31 | ATS-075 | A69-9001-30197 | 355110140 | 950 | 1085 | 7.50 | SBU |
| 2011-10-31 | ATS-076 | A69-9001-30207 | 355110114 | 973 | 1126 | 8.50 | SBU |
| 2011-10-31 | ATS-077 | A69-9001-30221 | 355110119 | 1008 | 1175 | 9.20 | SBU |
| 2011-10-31 | ATS-078 | A69-9001-30198 | 350040134 | 1033 | 1187 | 9.60 | SBU |
| 2011-10-31 | ATS-079 | A69-9001-30306 | 355110083 | 1060 | 1247 | 10.30 | SBU |
| 2011-10-31 | ATS-080 | A69-9001-30311 | 355110088 | 1070 | 1210 | 10.40 | SBU |
| 2011-10-31 | ATS-081 | A69-9001-30328 | 355110091 | 1070 | 1265 | 10.50 | SBU |
| 2011-10-31 | ATS-082 | A69-9001-30220 | 355110122 | 1060 | 1225 | 10.50 | SBU |
| 2011-10-31 | ATS-083 | A69-9001-30326 | 350060121 | 1120 | 1295 | 11.00 | SBU |
| 2011-10-31 | ATS-084 | A69-9001-30224 | 355110125 | 1080 | 1240 | 11.20 | SBU |
| 2011-10-31 | ATS-085 | A69-9001-30232 | 355110133 | 1115 | 1300 | 11.50 | SBU |
| 2011-10-31 | ATS-086 | A69-9001-30196 | 355110135 | 1125 | 1315 | 12.00 | SBU |
| 2011-10-31 | ATS-087 | A69-9001-30322 | 355110098 | 1080 | 1200 | 12.10 | SBU |
| 2011-10-31 | ATS-088 | A69-9001-30223 | 355110126 | 1140 | 1340 | 12.20 | SBU |
| 2011-10-31 | ATS-089 | A69-9001-30227 | 355110127 | 1130 | 1285 | 12.20 | SBU |
| 2011-10-31 | ATS-090 | A69-9001-30307 | 355110092 | 1090 | 1295 | 12.50 | SBU |
| 2011-10-31 | ATS-091 | A69-9001-30309 | 355110086 | 1110 | 1300 | 12.70 | SBU |
| 2011-10-31 | ATS-092 | A69-9001-30228 | 355110124 | 1140 | 1335 | 12.70 | SBU |
| 2011-10-31 | ATS-093 | A69-9001-30308 | 355110087 | 1100 | 1225 | 12.80 | SBU |
| 2011-10-31 | ATS-094 | A69-9001-30325 | 355110085 | 1130 | 1300 | 13.00 | SBU |
| 2011-10-31 | ATS-095 | A69-9001-30313 | 355110102 | 1080 | 1285 | 13.00 | SBU |
| 2011-10-31 | ATS-096 | A69-9001-30225 | 355110129 | 1180 | 1345 | 13.50 | SBU |
| 2011-10-31 | ATS-097 | A69-9001-30312 | 355110103 | 1097 | 1290 | 13.60 | SBU |
| 2011-10-31 | ATS-098 | A69-9001-30215 | 355110111 | 1160 | 1385 | 13.70 | SBU |
| 2011-10-31 | ATS-099 | A69-9001-30314 | 355110104 | 1212 | 1403 | 15.00 | SBU |
| 2011-10-31 | ATS-100 | A69-9001-30229 | 355110130 | 1230 | 1422 | 15.30 | SBU |
| 2011-10-31 | ATS-101 | A69-9001-30320 | 355110096 | 1235 | 1409 | 15.40 | SBU |
| 2011-10-31 | ATS-102 | A69-9001-30323 | 355110097 | 1210 | 1409 | 15.40 | SBU |
| 2011-10-31 | ATS-103 | A69-9001-30218 | 355110121 | 1135 | 1320 | 15.50 | SBU |
| 2011-10-31 | ATS-104 | A69-9001-30327 | 355110095 | 1275 | 1460 | 16.70 | SBU |
| 2011-10-31 | ATS-105 | A69-9001-30310 | 355110089 | 1287 | 1504 | 17.30 | SBU |
| 2011-10-31 | ATS-106 | A69-9001-30315 | 355110175 | 1225 | 1424 | 17.30 | SBU |
| 2011-10-31 | ATS-107 | A69-9001-30217 | 355110112 | 1209 | 1390 | 17.70 | SBU |
| 2011-10-31 | ATS-108 | A69-9001-30226 | 355110123 | 1240 | 1435 | 19.30 | SBU |
| 2011-10-31 | ATS-109 | A69-9001-30231 | 355110132 | 1280 | 1479 | 19.50 | SBU |
| 2011-10-31 | ATS-110 | A69-9001-30213 | 355110117 | 1320 | 1536 | 20.00 | SBU |
| 2011-10-31 | ATS-111 | A69-9001-30199 | 355110136 | 1380 | 1570 | 21.50 | SBU |
| 2011-10-31 | ATS-112 | A69-9001-30208 | 355110109 | 1454 | 1676 | 26.40 | SBU |
| 2011-10-31 | ATS-113 | A69-9001-30219 | 355110120 | 1447 | 1682 | 30.50 | SBU |
| 2011-10-31 | ATS-114 | A69-9001-30230 | 355110137 | 1810 | 2070 | 85.00 | SBU |
| 2011-11-08 | ATS-115 | A69-9001-30262 | 355110185 | 756 | 892 | 3.20 | SBU |
| 2011-11-08 | ATS-116 | A69-9001-30240 | 354080034 | 814 | 925 | 4.29 | SBU |
| 2011-11-08 | ATS-117 | A69-9001-30253 | 355110169 | 829 | 966 | 4.69 | SBU |
| 2011-11-08 | ATS-118 | A69-9001-30202 | 350080032 | 855 | 980 | 4.99 | SBU |
| 2011-11-08 | ATS-119 | A69-9001-30250 | 355110167 | 875 | 1035 | 5.24 | SBU |
| 2011-11-08 | ATS-120 | A69-9001-30238 | 355110157 | 965 | 1080 | 6.68 | SBU |
| 2011-11-08 | ATS-121 | A69-9001-30187 | 355110146 | 950 | 1110 | 6.99 | SBU |
| 2011-11-08 | ATS-122 | A69-9001-30237 | 355110162 | 966 | 1120 | 7.32 | SBU |
| 2011-11-08 | ATS-123 | A69-9001-30263 | 355110178 | 947 | 1055 | 7.42 | SBU |
| 2011-11-08 | ATS-124 | A69-9001-30234 | 355110172 | 979 | 1105 | 7.81 | SBU |
| 2011-11-08 | ATS-125 | A69-9001-30261 | 355110180 | 957 | 1090 | 8.19 | SBU |
| 2011-11-08 | ATS-126 | A69-9001-30247 | 355110168 | 1027 | 1165 | 8.56 | SBU |
| 2011-11-08 | ATS-127 | A69-9001-30186 | 355110147 | 1005 | 1120 | 8.69 | SBU |
| 2011-11-08 | ATS-128 | A69-9001-30243 | 355110156 | 1020 | 1150 | 9.62 | SBU |
| 2011-11-08 | ATS-129 | A69-9001-30184 | 355110148 | 1185 | 1380 | 9.71 | SBU |
| 2011-11-08 | ATS-130 | A69-9001-30260 | 355110182 | 1069 | 1262 | 10.18 | SBU |
| 2011-11-08 | ATS-131 | A69-9001-30256 | 355110179 | 1081 | 1282 | 10.52 | SBU |
| 2011-11-08 | ATS-132 | A69-9001-30241 | 355110161 | 1120 | 1325 | 11.56 | SBU |
| 2011-11-08 | ATS-133 | A69-9001-30319 | 355110150 | 1140 | 1320 | 12.14 | SBU |
| 2011-11-08 | ATS-134 | A69-9001-30242 | 355110163 | 1138 | 1305 | 13.35 | SBU |
| 2011-11-08 | ATS-135 | A69-9001-30318 | 355110151 | 1155 | 1335 | 13.36 | SBU |
| 2011-11-08 | ATS-136 | A69-9001-30248 | 355110166 | 1165 | 1350 | 14.72 | SBU |
| 2011-11-08 | ATS-137 | A69-9001-30191 | 355110152 | 1220 | 1395 | 14.74 | SBU |
| 2011-11-08 | ATS-138 | A69-9001-30244 | 355110158 | 1148 | 1285 | 14.80 | SBU |
| 2011-11-08 | ATS-139 | A69-9001-30257 | 355110171 | 1184 | 1376 | 14.90 | SBU |
| 2011-11-08 | ATS-140 | A69-9001-30255 | 355110164 | 1250 | 1448 | 14.95 | SBU |
| 2011-11-08 | ATS-141 | A69-9001-30236 | 355110173 | 1190 | 1330 | 15.13 | SBU |
| 2011-11-08 | ATS-142 | A69-9001-30194 | 350060036 | 1192 | 1345 | 15.34 | SBU |
| 2011-11-08 | ATS-143 | A69-9001-30192 | 355110145 | 1215 | 1440 | 15.56 | SBU |
| 2011-11-08 | ATS-144 | A69-9001-30245 | 355110154 | 1156 | 1350 | 15.90 | SBU |
| 2011-11-08 | ATS-145 | A69-9001-30264 | 355110183 | 1258 | 1430 | 16.14 | SBU |
| 2011-11-08 | ATS-146 | A69-9001-30188 | 355110142 | 1280 | 1440 | 16.36 | SBU |
| 2011-11-08 | ATS-147 | A69-9001-30258 | 355110184 | 1258 | 1446 | 16.44 | SBU |
| 2011-11-08 | ATS-148 | A69-9001-30189 | 355110143 | 1260 | 1470 | 17.25 | SBU |
| 2011-11-08 | ATS-149 | A69-9001-30246 | 355110165 | 1255 | 1430 | 17.28 | SBU |
| 2011-11-08 | ATS-150 | A69-9001-30254 | 355110176 | 1240 | 1470 | 17.50 | SBU |
| 2011-11-08 | ATS-151 | A69-9001-30249 | 355110174 | 1254 | 1490 | 17.67 | SBU |
| 2011-11-08 | ATS-152 | A69-9001-30193 | 355110153 | 1275 | 1490 | 17.86 | SBU |
| 2011-11-08 | ATS-153 | A69-9001-30259 | 355110181 | 1305 | 1500 | 18.11 | SBU |
| 2011-11-08 | ATS-154 | A69-9001-30252 | 355110170 | 1310 | 1492 | 18.26 | SBU |
| 2011-11-08 | ATS-155 | A69-9001-30251 | 355110175 | 1340 | 1560 | 20.82 | SBU |
| 2011-11-09 | ATS-156 | A69-9001-30280 | 350040044 | 401 | 466 | 0.45 | SBU |
| 2011-11-09 | ATS-157 | A69-9001-30292 | 355110216 | 719 | 830 | 3.03 | SBU |
| 2011-11-09 | ATS-158 | A69-9001-30275 | 355110199 | 746 | 854 | 3.21 | SBU |
| 2011-11-09 | ATS-159 | A69-9001-30269 | 355110187 | 786 | 917 | 3.56 | SBU |
| 2011-11-09 | ATS-160 | A69-9001-30291 | 355110211 | 782 | 918 | 3.79 | SBU |
| 2011-11-09 | ATS-161 | A69-9001-30271 | 355110188 | 849 | 980 | 5.08 | SBU |
| 2011-11-09 | ATS-162 | A69-9001-30282 | 355110210 | 886 | 1015 | 5.15 | SBU |
| 2011-11-09 | ATS-163 | A69-9001-30016 | 355110222 | 849 | 985 | 5.21 | SBU |
| 2011-11-09 | ATS-164 | A69-9001-30276 | 350080042 | 856 | 990 | 5.70 | SBU |
| 2011-11-09 | ATS-165 | A69-9001-30304 | 355110205 | 906 | 1025 | 6.08 | SBU |
| 2011-11-09 | ATS-166 | A69-9001-30301 | 355110215 | 926 | 1065 | 6.35 | SBU |
| 2011-11-09 | ATS-167 | A69-9001-30293 | 355110212 | 934 | 1050 | 6.88 | SBU |
| 2011-11-09 | ATS-168 | A69-9001-30303 | 355110201 | 938 | 1065 | 6.95 | SBU |
| 2011-11-09 | ATS-169 | A69-9001-30278 | 355110194 | 940 | 1075 | 7.08 | SBU |
| 2011-11-09 | ATS-170 | A69-9001-30274 | 355110193 | 935 | 1085 | 7.12 | SBU |
| 2011-11-09 | ATS-171 | A69-9001-30284 | 355110219 | 921 | 1060 | 7.13 | SBU |
| 2011-11-09 | ATS-172 | A69-9001-30268 | 355110196 | 953 | 1078 | 7.92 | SBU |
| 2011-11-09 | ATS-173 | A69-9001-30267 | 355110186 | 982 | 1125 | 9.00 | SBU |
| 2011-11-09 | ATS-174 | A69-9001-30305 | 355110202 | 1024 | 1188 | 9.17 | SBU |
| 2011-11-09 | ATS-175 | A69-9001-30281 | 355110190 | 1044 | 1200 | 9.72 | SBU |
| 2011-11-09 | ATS-176 | A69-9001-30296 | 355110207 | 997 | 1145 | 9.90 | SBU |
| 2011-11-09 | ATS-177 | A69-9001-30285 | 355110213 | 1046 | 1195 | 10.30 | SBU |
| 2011-11-09 | ATS-178 | A69-9001-30298 | 355110206 | 1132 | 1265 | 12.75 | SBU |
| 2011-11-09 | ATS-179 | A69-9001-30290 | 355110218 | 1225 | 1370 | 15.75 | SBU |
| 2011-11-09 | ATS-180 | A69-9001-30286 | 355110221 | 1224 | 1400 | 16.72 | SBU |
| 2011-11-09 | ATS-181 | A69-9001-30273 | 355110195 | 1144 | 1285 | 17.43 | SBU |
| 2011-11-09 | ATS-182 | A69-9001-30279 | 355110189 | 1331 | 1500 | 18.97 | SBU |
| 2011-11-09 | ATS-183 | A69-9001-30289 | 355110208 | 1343 | 1580 | 26.32 | SBU |
| 2011-11-09 | ATS-184 | A69-9001-30270 | 355110249 | 1640 | 1780 | 32.76 | SBU |
| 2011-11-10 | ATS-185 | A69-9001-30015 | 355060010 | 1236 | 1434 | 13.32 | SBU |
| 2011-11-10 | ATS-186 | A69-9001-30017 | 350090012 | 935 | 1070 | 6.41 | SBU |
| 2011-11-10 | ATS-187 | A69-9001-30036 | 355110235 | 773 | 918 | 3.70 | SBU |
| 2011-11-10 | ATS-188 | A69-9001-30026 | 355110237 | 796 | 931 | 3.85 | SBU |
| 2011-11-10 | ATS-189 | A69-9001-30035 | 355110244 | 813 | 942 | 3.96 | SBU |
| 2011-11-10 | ATS-190 | A69-9001-30034 | 355110234 | 793 | 916 | 4.11 | SBU |
| 2011-11-10 | ATS-191 | A69-9001-30029 | 355110241 | 819 | 932 | 4.42 | SBU |
| 2011-11-10 | ATS-192 | A69-9001-30020 | 355110226 | 839 | 945 | 4.82 | SBU |
| 2011-11-10 | ATS-193 | A69-9001-30046 | 355110247 | 864 | 985 | 4.94 | SBU |
| 2011-11-10 | ATS-194 | A69-9001-30300 | 355110224 | 873 | 980 | 5.50 | SBU |
| 2011-11-10 | ATS-195 | A69-9001-30023 | 355110236 | 914 | 1047 | 6.24 | SBU |
| 2011-11-10 | ATS-196 | A69-9001-30018 | 355110223 | 940 | 1065 | 6.32 | SBU |
| 2011-11-10 | ATS-197 | A69-9001-30019 | 355110229 | 904 | 1050 | 6.77 | SBU |
| 2011-11-10 | ATS-198 | A69-9001-30021 | 355110231 | 943 | 1093 | 6.84 | SBU |
| 2011-11-10 | ATS-199 | A69-9001-30032 | 355110233 | 956 | 1099 | 7.44 | SBU |
| 2011-11-10 | ATS-200 | A69-9001-30025 | 355110232 | 1027 | 1205 | 8.90 | SBU |
| 2011-11-10 | ATS-201 | A69-9001-30033 | 355110243 | 1019 | 1180 | 9.03 | SBU |
| 2011-11-10 | ATS-202 | A69-9001-30030 | 355110239 | 1124 | 1287 | 12.72 | SBU |
| 2011-11-10 | ATS-203 | A69-9001-30028 | 355110238 | 1213 | 1363 | 14.62 | SBU |
| 2011-11-10 | ATS-204 | A69-9001-30037 | 355110248 | 1177 | 1330 | 14.98 | SBU |
| 2011-11-10 | ATS-205 | A69-9001-30024 | 355110230 | 1188 | 1325 | 16.48 | SBU |
| 2011-11-10 | ATS-206 | A69-9001-30022 | 355110228 | 1270 | 1466 | 17.92 | SBU |
| 2011-11-10 | ATS-207 | A69-9001-30027 | 355110240 | 1264 | 1444 | 18.13 | SBU |
| 2011-11-10 | ATS-208 | A69-9001-30031 | 355110245 | 1265 | 1460 | 19.26 | SBU |
| 2012-05-02 | ATS-209 | A69-9001-30079 | 355120120 | 798 | 914 | 1.60 | SBU |
| 2012-05-02 | ATS-210 | A69-9001-30082 | 355120121 | 795 | 926 | 1.90 | SBU |
| 2012-05-02 | ATS-211 | A69-9001-30066 | 355120135 | 778 | 884 | 3.38 | SBU |
| 2012-05-02 | ATS-212 | A69-9001-30071 | 355120138 | 725 | 836 | 3.63 | SBU |
| 2012-05-02 | ATS-213 | A69-9001-30074 | 355120131 | 794 | 908 | 4.24 | SBU |
| 2012-05-02 | ATS-214 | A69-9001-30061 | 355120134 | 769 | 885 | 4.30 | SBU |
| 2012-05-02 | ATS-215 | A69-9001-30051 | 355120152 | 825 | 885 | 4.44 | SBU |
| 2012-05-02 | ATS-216 | A69-9001-30090 | 355120117 | 781 | 891 | 4.58 | SBU |
| 2012-05-02 | ATS-217 | A69-9001-30058 | 355120144 | 811 | 942 | 4.70 | SBU |
| 2012-05-02 | ATS-218 | A69-9001-30089 | 355120124 | 836 | 970 | 5.09 | SBU |
| 2012-05-02 | ATS-219 | A69-9001-30070 | 355120142 | 852 | 988 | 5.16 | SBU |
| 2012-05-02 | ATS-220 | A69-9001-30052 | 355120149 | 815 | 940 | 5.28 | SBU |
| 2012-05-02 | ATS-221 | A69-9001-30063 | 355120113 | 815 | 956 | 5.30 | SBU |
| 2012-05-02 | ATS-222 | A69-9001-30075 | 355120127 | 823 | 928 | 5.36 | SBU |
| 2012-05-02 | ATS-223 | A69-9001-30056 | 355120132 | 881 | 984 | 5.36 | SBU |
| 2012-05-02 | ATS-224 | A69-9001-30050 | 355120151 | 880 | 970 | 5.43 | SBU |
| 2012-05-02 | ATS-225 | A69-9001-30093 | 355120106 | 846 | 980 | 5.66 | SBU |
| 2012-05-02 | ATS-226 | A69-9001-30065 | 355120110 | 860 | 995 | 5.85 | SBU |
| 2012-05-02 | ATS-227 | A69-9001-30073 | 355120139 | 860 | 998 | 5.96 | SBU |
| 2012-05-02 | ATS-228 | A69-9001-30092 | 355120109 | 897 | 1038 | 6.01 | SBU |
| 2012-05-02 | ATS-229 | A69-9001-30095 | 355120108 | 888 | 1032 | 6.15 | SBU |
| 2012-05-02 | ATS-230 | A69-9001-30087 | 355120123 | 887 | 1022 | 6.19 | SBU |
| 2012-05-02 | ATS-231 | A69-9001-30088 | 355120118 | 917 | 1074 | 6.24 | SBU |
| 2012-05-02 | ATS-232 | A69-9001-30080 | 355120126 | 896 | 1029 | 6.24 | SBU |
| 2012-05-02 | ATS-233 | A69-9001-30072 | 355120143 | 906 | 1060 | 6.33 | SBU |
| 2012-05-02 | ATS-234 | A69-9001-30086 | 355120122 | 911 | 1039 | 6.44 | SBU |
| 2012-05-02 | ATS-235 | A69-9001-30081 | 355120114 | 925 | 1065 | 6.67 | SBU |
| 2012-05-02 | ATS-236 | A69-9001-30091 | 355120125 | 907 | 1053 | 7.01 | SBU |
| 2012-05-02 | ATS-237 | A69-9001-30078 | 355120128 | 961 | 1098 | 7.49 | SBU |
| 2012-05-02 | ATS-238 | A69-9001-30068 | 355120137 | 940 | 1065 | 7.64 | SBU |
| 2012-05-02 | ATS-239 | A69-9001-30053 | 355120150 | 891 | 1020 | 7.73 | SBU |
| 2012-05-02 | ATS-240 | A69-9001-30064 | 355120111 | 951 | 1093 | 7.84 | SBU |
| 2012-05-02 | ATS-241 | A69-9001-30076 | 355120119 | 947 | 1074 | 8.40 | SBU |
| 2012-05-02 | ATS-242 | A69-9001-30069 | 355120141 | 972 | 1110 | 9.42 | SBU |
| 2012-05-02 | ATS-243 | A69-9001-30067 | 355120140 | 1004 | 1149 | 9.65 | SBU |
| 2012-05-02 | ATS-244 | A69-9001-30083 | 355120115 | 1034 | 1195 | 10.16 | SBU |
| 2012-05-02 | ATS-245 | A69-9001-30085 | 355120116 | 1018 | 1159 | 10.34 | SBU |
| 2012-05-02 | ATS-246 | A69-9001-30084 | 355120175 | 1021 | 1202 | 10.42 | SBU |
| 2012-05-02 | ATS-247 | A69-9001-30060 | 355120136 | 1030 | 1178 | 10.44 | SBU |
| 2012-05-02 | ATS-248 | A69-9001-30077 | 355120129 | 1059 | 1205 | 10.98 | SBU |
| 2012-05-02 | ATS-249 | A69-9001-30062 | 355120112 | 1081 | 1250 | 12.28 | SBU |
| 2012-05-02 | ATS-250 | A69-9001-30055 | 355120148 | 1140 | 1280 | 14.88 | SBU |
| 2012-05-03 | ATS-251 | A69-9001-30148 | 355120080 | 769 | 886 | 3.66 | SBU |
| 2012-05-03 | ATS-252 | A69-9001-30100 | 355120100 | 760 | 836 | 3.72 | SBU |
| 2012-05-03 | ATS-253 | A69-9001-30152 | 355120077 | 792 | 928 | 4.30 | SBU |
| 2012-05-03 | ATS-254 | A69-9001-30038 | 355120059 | 805 | 936 | 4.50 | SBU |
| 2012-05-03 | ATS-255 | A69-9001-30156 | 355120074 | 826 | 967 | 4.84 | SBU |
| 2012-05-03 | ATS-256 | A69-9001-30128 | 355120051 | 857 | 984 | 5.18 | SBU |
| 2012-05-03 | ATS-257 | A69-9001-30116 | 355120062 | 825 | 952 | 5.21 | SBU |
| 2012-05-03 | ATS-258 | A69-9001-30094 | 355120105 | 823 | 956 | 5.22 | SBU |
| 2012-05-03 | ATS-259 | A69-9001-30117 | 355120063 | 814 | 944 | 5.24 | SBU |
| 2012-05-03 | ATS-260 | A69-9001-30042 | 355120060 | 819 | 948 | 5.30 | SBU |
| 2012-05-03 | ATS-261 | A69-9001-30136 | 355120071 | 860 | 986 | 5.42 | SBU |
| 2012-05-03 | ATS-262 | A69-9001-30131 | 355120057 | 823 | 945 | 5.46 | SBU |
| 2012-05-03 | ATS-263 | A69-9001-30150 | 355120076 | 893 | 995 | 5.58 | SBU |
| 2012-05-03 | ATS-264 | A69-9001-30123 | 355120053 | 864 | 1007 | 5.64 | SBU |
| 2012-05-03 | ATS-265 | A69-9001-30040 | 355120043 | 828 | 959 | 5.70 | SBU |
| 2012-05-03 | ATS-266 | A69-9001-30105 | 355120089 | 861 | 968 | 5.70 | SBU |
| 2012-05-03 | ATS-267 | A69-9001-30096 | 355120011 | 848 | 984 | 5.74 | SBU |
| 2012-05-03 | ATS-268 | A69-9001-30129 | 355120056 | 874 | 1001 | 5.74 | SBU |
| 2012-05-03 | ATS-269 | A69-9001-30107 | 355120087 | 861 | 1014 | 5.83 | SBU |
| 2012-05-03 | ATS-270 | A69-9001-30155 | 355120079 | 871 | 1019 | 5.92 | SBU |
| 2012-05-03 | ATS-271 | A69-9001-30118 | 355120094 | 843 | 982 | 5.94 | SBU |
| 2012-05-03 | ATS-272 | A69-9001-30121 | 355120096 | 864 | 993 | 5.98 | SBU |
| 2012-05-03 | ATS-273 | A69-9001-30153 | 355120078 | 859 | 985 | 6.46 | SBU |
| 2012-05-03 | ATS-274 | A69-9001-30149 | 355120046 | 888 | 1037 | 6.54 | SBU |
| 2012-05-03 | ATS-275 | A69-9001-30102 | 355120103 | 909 | 1052 | 6.64 | SBU |
| 2012-05-03 | ATS-276 | A69-9001-30151 | 355120045 | 860 | 1000 | 6.70 | SBU |
| 2012-05-03 | ATS-277 | A69-9001-30043 | 355120034 | 947 | 1089 | 6.94 | SBU |
| 2012-05-03 | ATS-278 | A69-9001-30154 | 355120084 | 900 | 1066 | 7.22 | SBU |
| 2012-05-03 | ATS-279 | A69-9001-30177 | 355120037 | 936 | 1076 | 7.36 | SBU |
| 2012-05-03 | ATS-280 | A69-9001-30104 | 355120088 | 926 | 1062 | 7.74 | SBU |
| 2012-05-03 | ATS-281 | A69-9001-30099 | 355120101 | 967 | 1132 | 8.01 | SBU |
| 2012-05-03 | ATS-282 | A69-9001-30120 | 355120095 | 938 | 1086 | 8.03 | SBU |
| 2012-05-03 | ATS-283 | A69-9001-30145 | 355120086 | 923 | 1062 | 8.12 | SBU |
| 2012-05-03 | ATS-284 | A69-9001-30146 | 355120075 | 963 | 1114 | 8.26 | SBU |
| 2012-05-03 | ATS-285 | A69-9001-30047 | 355120036 | 981 | 1091 | 8.36 | SBU |
| 2012-05-03 | ATS-286 | A69-9001-30147 | 355120044 | 983 | 1154 | 8.80 | SBU |
| 2012-05-03 | ATS-287 | A69-9001-30130 | 355120050 | 980 | 1134 | 9.36 | SBU |
| 2012-05-03 | ATS-288 | A69-9001-30108 | 355120098 | 1021 | 1166 | 9.40 | SBU |
| 2012-05-03 | ATS-289 | A69-9001-30119 | 355120061 | 975 | 1136 | 9.78 | SBU |
| 2012-05-03 | ATS-290 | A69-9001-30115 | 355120090 | 1020 | 1218 | 10.46 | SBU |
| 2012-05-03 | ATS-291 | A69-9001-30101 | 355120099 | 1023 | 1182 | 10.56 | SBU |
| 2012-05-03 | ATS-292 | A69-9001-30179 | 355120038 | 1039 | 1194 | 10.94 | SBU |
| 2012-05-03 | ATS-293 | A69-9001-30106 | 355120092 | 1022 | 1198 | 11.26 | SBU |
| 2012-05-03 | ATS-294 | A69-9001-30142 | 355120069 | 1069 | 1234 | 11.62 | SBU |
| 2012-05-03 | ATS-295 | A69-9001-30181 | 355120041 | 1077 | 1234 | 11.86 | SBU |
| 2012-05-03 | ATS-296 | A69-9001-30122 | 355120067 | 1050 | 1163 | 12.24 | SBU |
| 2012-05-03 | ATS-297 | A69-9001-30126 | 355120049 | 1093 | 1267 | 12.62 | SBU |
| 2012-05-03 | ATS-298 | A69-9001-30144 | 355120068 | 1089 | 1256 | 12.86 | SBU |
| 2012-05-03 | ATS-299 | A69-9001-30143 | 355120073 | 1097 | 1263 | 13.20 | SBU |
| 2012-05-03 | ATS-300 | A69-9001-30133 | 355120058 | 1150 | 1364 | 14.56 | SBU |
| 2012-05-03 | ATS-301 | A69-9001-30111 | 355120065 | 1175 | 1341 | 14.98 | SBU |
| 2012-05-03 | ATS-302 | A69-9001-30127 | 355120055 | 1152 | 1328 | 15.14 | SBU |
| 2012-05-03 | ATS-303 | A69-9001-30138 | 355120070 | 1181 | 1348 | 15.38 | SBU |
| 2012-05-03 | ATS-304 | A69-9001-30132 | 355120052 | 1180 | 1377 | 15.40 | SBU |
| 2012-05-03 | ATS-305 | A69-9001-30178 | 355120039 | 1161 | 1299 | 16.44 | SBU |
| 2012-05-03 | ATS-306 | A69-9001-30137 | 355120082 | 1189 | 1407 | 16.82 | SBU |
| 2012-05-03 | ATS-307 | A69-9001-30134 | 355120072 | 1251 | 1423 | 19.42 | SBU |
| 2012-05-03 | ATS-308 | A69-9001-30044 | 355120042 | 1225 | 1421 | 20.16 | SBU |
| 2012-05-03 | ATS-309 | A69-9001-30125 | 355120054 | 1281 | 1462 | 20.26 | SBU |
| 2012-05-03 | ATS-310 | A69-9001-30114 | 355120091 | 1318 | 1379 | 21.80 | SBU |
| 2012-05-03 | ATS-311 | A69-9001-30112 | 355120008 | 1334 | 1530 | 22.94 | SBU |
| 2012-05-03 | ATS-312 | A69-9001-30135 | 355120081 | 1275 | 1484 | 23.12 | SBU |
| 2012-05-03 | ATS-313 | A69-9001-30109 | 355120097 | 1279 | 1495 | 23.28 | SBU |
| 2012-05-03 | ATS-314 | A69-9001-30140 | 355120048 | 1450 | 1705 | 27.02 | SBU |
| 2012-05-03 | ATS-315 | A69-9001-30098 | 355120102 | 1434 | 1665 | 28.60 | SBU |
| 2012-05-04 | ATS-316 | A69-9001-30172 | 355120023 | 854 | 981 | 4.85 | SBU |
| 2012-05-04 | ATS-317 | A69-9001-30162 | 355120016 | 858 | 1007 | 5.53 | SBU |
| 2012-05-04 | ATS-318 | A69-9001-30163 | 355120018 | 875 | 1011 | 5.84 | SBU |
| 2012-05-04 | ATS-319 | A69-9001-30175 | 355120027 | 863 | 1009 | 6.22 | SBU |
| 2012-05-04 | ATS-320 | A69-9001-30171 | 355120022 | 884 | 1000 | 6.36 | SBU |
| 2012-05-04 | ATS-321 | A69-9001-30157 | 355120021 | 889 | 1017 | 6.62 | SBU |
| 2012-05-04 | ATS-322 | A69-9001-30039 | 355120025 | 929 | 1040 | 7.70 | SBU |
| 2012-05-04 | ATS-323 | A69-9001-30158 | 355120019 | 947 | 1109 | 8.35 | SBU |
| 2012-05-04 | ATS-324 | A69-9001-30041 | 355120005 | 958 | 1117 | 9.78 | SBU |
| 2012-05-04 | ATS-325 | A69-9001-30168 | 355120032 | 1015 | 1193 | 9.84 | SBU |
| 2012-05-04 | ATS-326 | A69-9001-30166 | 355120031 | 1014 | 1189 | 9.86 | SBU |
| 2012-05-04 | ATS-327 | A69-9001-30164 | 355120020 | 1022 | 1165 | 10.59 | SBU |
| 2012-05-04 | ATS-328 | A69-9001-30173 | 355120029 | 1054 | 1218 | 12.20 | SBU |
| 2012-05-04 | ATS-329 | A69-9001-30167 | 355120030 | 1298 | 1553 | 21.22 | SBU |
| 2012-05-04 | ATS-330 | A69-9001-30159 | 355120012 | 1344 | 1513 | 22.00 | SBU |
| 2012-05-04 | ATS-331 | A69-9001-30169 | 355120026 | 1316 | 1419 | 28.50 | SBU |
| 2012-05-04 | ATS-332 | A69-9001-30165 | 355120024 | 1785 | 2033 | 54.50 | SBU |
| 2014-05-28 | ATS-333 | A69-9001-28857 | 355140004 | 718 | 815 | 4.03 | SBU |
| 2014-05-28 | ATS-334 | A69-9001-28841 | 355140013 | 849 | 972 | 4.28 | SBU |
| 2014-05-28 | ATS-335 | A69-9001-28843 | 355140015 | 815 | 922 | 4.56 | SBU |
| 2014-05-28 | ATS-336 | A69-9001-28853 | 355140006 | 820 | 941 | 4.64 | SBU |
| 2014-05-28 | ATS-337 | A69-9001-28856 | 355140007 | 815 | 915 | 4.72 | SBU |
| 2014-05-28 | ATS-338 | A69-9001-28854 | 355140005 | 798 | 910 | 4.80 | SBU |
| 2014-05-28 | ATS-339 | A69-9001-28861 | 355140001 | 825 | 905 | 5.00 | SBU |
| 2014-05-28 | ATS-340 | A69-9001-28860 | 355140010 | 868 | 998 | 6.28 | SBU |
| 2014-05-28 | ATS-341 | A69-9001-28859 | 355140002 | 890 | 985 | 6.55 | SBU |
| 2014-05-28 | ATS-342 | A69-9001-28846 | 355140019 | 918 | 1037 | 6.64 | SBU |
| 2014-05-28 | ATS-343 | A69-9001-28849 | 355140016 | 944 | 1066 | 7.68 | SBU |
| 2014-05-28 | ATS-344 | A69-9001-28845 | 355140014 | 982 | 1077 | 9.40 | SBU |
| 2014-05-28 | ATS-345 | A69-9001-28847 | 355140017 | 1053 | 1208 | 9.80 | SBU |
| 2014-05-28 | ATS-346 | A69-9001-28858 | 355140008 | 1024 | 1165 | 9.85 | SBU |
| 2014-05-28 | ATS-347 | A69-9001-28844 | 355140018 | 1043 | 1150 | 10.32 | SBU |
| 2014-05-28 | ATS-348 | A69-9001-28851 | 355140021 | 1054 | 1201 | 10.34 | SBU |
| 2014-05-28 | ATS-349 | A69-9001-28855 | 355140003 | 1001 | 1160 | 10.68 | SBU |
| 2014-05-28 | ATS-350 | A69-9001-28862 | 355140011 | 1063 | 1232 | 10.86 | SBU |
| 2014-05-28 | ATS-351 | A69-9001-28840 | 355140012 | 1078 | 1225 | 11.74 | SBU |
| 2014-05-28 | ATS-352 | A69-9001-28842 | 355140020 | 1142 | 1313 | 14.04 | SBU |
| 2014-05-28 | ATS-353 | A69-9001-28863 | 355140009 | 1144 | 1305 | 15.58 | SBU |
| 2014-05-29 | ATS-354 | A69-9001-28848 | 355140024 | 827 | 943 | 5.23 | SBU |
| 2014-05-29 | ATS-355 | A69-9001-28850 | 355140023 | 954 | 1088 | 7.60 | SBU |
| 2014-05-29 | ATS-356 | A69-9001-28852 | 355140022 | 948 | 1103 | 7.95 | SBU |
| 2015-05-19 | ATS-357 | A69-9001-21930 | 355150018 | 738 | 856 | 2.98 | SBU |
| 2015-05-19 | ATS-358 | A69-9001-21921 | 355150041 | 724 | 849 | 3.16 | SBU |
| 2015-05-19 | ATS-359 | A69-9001-21955 | 355150013 | 792 | 906 | 3.85 | SBU |
| 2015-05-19 | ATS-360 | A69-9001-21931 | 355150020 | 770 | 915 | 3.98 | SBU |
| 2015-05-19 | ATS-361 | A69-9001-21956 | 355150011 | 781 | 915 | 4.19 | SBU |
| 2015-05-19 | ATS-362 | A69-9001-21920 | 355150035 | 819 | 941 | 4.35 | SBU |
| 2015-05-19 | ATS-363 | A69-9001-21939 | 355150027 | 820 | 957 | 4.48 | SBU |
| 2015-05-19 | ATS-364 | A69-9001-21946 | 355150001 | 819 | 940 | 4.67 | SBU |
| 2015-05-19 | ATS-365 | A69-9001-21919 | 355150039 | 842 | 986 | 4.70 | SBU |
| 2015-05-19 | ATS-366 | A69-9001-21945 | 355150007 | 827 | 969 | 4.72 | SBU |
| 2015-05-19 | ATS-367 | A69-9001-21957 | 355150012 | 813 | 954 | 4.80 | SBU |
| 2015-05-19 | ATS-368 | A69-9001-21940 | 355150026 | 829 | 956 | 4.84 | SBU |
| 2015-05-19 | ATS-369 | A69-9001-21954 | 355150014 | 819 | 949 | 4.85 | SBU |
| 2015-05-19 | ATS-370 | A69-9001-21936 | 355150024 | 842 | 977 | 4.86 | SBU |
| 2015-05-19 | ATS-371 | A69-9001-21929 | 355150031 | 852 | 992 | 4.90 | SBU |
| 2015-05-19 | ATS-372 | A69-9001-21924 | 355150038 | 838 | 996 | 4.95 | SBU |
| 2015-05-19 | ATS-373 | A69-9001-21942 | 355150016 | 863 | 993 | 5.29 | SBU |
| 2015-05-19 | ATS-374 | A69-9001-21950 | 355150004 | 843 | 980 | 5.50 | SBU |
| 2015-05-19 | ATS-375 | A69-9001-21925 | 355150034 | 860 | 991 | 5.63 | SBU |
| 2015-05-19 | ATS-376 | A69-9001-21933 | 355150021 | 881 | 1010 | 5.72 | SBU |
| 2015-05-19 | ATS-377 | A69-9001-21932 | 355150019 | 888 | 1002 | 6.10 | SBU |
| 2015-05-19 | ATS-378 | A69-9001-21948 | 355150002 | 881 | 1008 | 6.13 | SBU |
| 2015-05-19 | ATS-379 | A69-9001-21958 | 355150015 | 868 | 1003 | 6.27 | SBU |
| 2015-05-19 | ATS-380 | A69-9001-21951 | 355150006 | 901 | 1050 | 6.43 | SBU |
| 2015-05-19 | ATS-381 | A69-9001-21928 | 355150037 | 924 | 1080 | 6.63 | SBU |
| 2015-05-19 | ATS-382 | A69-9001-21941 | 355150029 | 894 | 1051 | 6.65 | SBU |
| 2015-05-19 | ATS-383 | A69-9001-21935 | 355150023 | 939 | 1096 | 6.70 | SBU |
| 2015-05-19 | ATS-384 | A69-9001-21934 | 355150022 | 906 | 1051 | 6.78 | SBU |
| 2015-05-19 | ATS-385 | A69-9001-21952 | 355150003 | 929 | 1070 | 7.02 | SBU |
| 2015-05-19 | ATS-386 | A69-9001-21944 | 355150008 | 918 | 1069 | 7.18 | SBU |
| 2015-05-19 | ATS-387 | A69-9001-21953 | 355150005 | 953 | 1115 | 7.19 | SBU |
| 2015-05-19 | ATS-388 | A69-9001-21949 | 355150009 | 954 | 1099 | 7.28 | SBU |
| 2015-05-19 | ATS-389 | A69-9001-21918 | 355150033 | 944 | 1001 | 7.30 | SBU |
| 2015-05-19 | ATS-390 | A69-9001-21927 | 355150036 | 950 | 1119 | 7.40 | SBU |
| 2015-05-19 | ATS-391 | A69-9001-21926 | 355150032 | 934 | 1094 | 7.44 | SBU |
| 2015-05-19 | ATS-392 | A69-9001-21943 | 355150017 | 988 | 1145 | 7.65 | SBU |
| 2015-05-19 | ATS-393 | A69-9001-21923 | 355150040 | 970 | 1141 | 7.65 | SBU |
| 2015-05-19 | ATS-394 | A69-9001-21947 | 355150010 | 960 | 1118 | 7.95 | SBU |
| 2015-05-19 | ATS-395 | A69-9001-21937 | 355150025 | 1004 | 1192 | 9.88 | SBU |
| 2015-05-19 | ATS-396 | A69-9001-21922 | 355150030 | 1080 | 1255 | 11.41 | SBU |
| 2015-05-19 | ATS-397 | A69-9001-21938 | 355150028 | 1153 | 1280 | 12.73 | SBU |
| 2016-05-04 | ATS-398 | A69-9001-18628 | 355160033 | 703 | 841 | 2.72 | SBU |
| 2016-05-04 | ATS-399 | A69-9001-18605 | 355160013 | 730 | 800 | 3.10 | SBU |
| 2016-05-04 | ATS-400 | A69-9001-18600 | 355160015 | 720 | 820 | 3.18 | SBU |
| 2016-05-04 | ATS-401 | A69-9001-18627 | 355160039 | 755 | 878 | 3.29 | SBU |
| 2016-05-04 | ATS-402 | A69-9001-18618 | 355160023 | 720 | 844 | 3.37 | SBU |
| 2016-05-04 | ATS-403 | A69-9001-18595 | 355160004 | 765 | 840 | 3.42 | SBU |
| 2016-05-04 | ATS-404 | A69-9001-18612 | 355160021 | 732 | 839 | 3.89 | SBU |
| 2016-05-04 | ATS-405 | A69-9001-18625 | 355160037 | 746 | 866 | 3.97 | SBU |
| 2016-05-04 | ATS-406 | A69-9001-18615 | 355160025 | 760 | 870 | 4.10 | SBU |
| 2016-05-04 | ATS-407 | A69-9001-18594 | 355160002 | 780 | 910 | 4.30 | SBU |
| 2016-05-04 | ATS-408 | A69-9001-18602 | 355160012 | 790 | 920 | 4.34 | SBU |
| 2016-05-04 | ATS-409 | A69-9001-18601 | 355160016 | 790 | 890 | 4.39 | SBU |
| 2016-05-04 | ATS-410 | A69-9001-18610 | 355160017 | 783 | 870 | 4.39 | SBU |
| 2016-05-04 | ATS-411 | A69-9001-18598 | 355160005 | 770 | 860 | 4.42 | SBU |
| 2016-05-04 | ATS-412 | A69-9001-18626 | 355160036 | 790 | 919 | 4.50 | SBU |
| 2016-05-04 | ATS-413 | A69-9001-18624 | 355160035 | 820 | 965 | 4.65 | SBU |
| 2016-05-04 | ATS-414 | A69-9001-18607 | 355160010 | 795 | 950 | 4.71 | SBU |
| 2016-05-04 | ATS-415 | A69-9001-18609 | 355160018 | 830 | 915 | 4.86 | SBU |
| 2016-05-04 | ATS-416 | A69-9001-18596 | 355160008 | 820 | 940 | 5.10 | SBU |
| 2016-05-04 | ATS-417 | A69-9001-18622 | 355160034 | 1009 | 1126 | 5.50 | SBU |
| 2016-05-04 | ATS-418 | A69-9001-18604 | 355160011 | 853 | 969 | 5.52 | SBU |
| 2016-05-04 | ATS-419 | A69-9001-18619 | 355160030 | 843 | 962 | 5.62 | SBU |
| 2016-05-04 | ATS-420 | A69-9001-18606 | 355160009 | 907 | 1033 | 5.91 | SBU |
| 2016-05-04 | ATS-421 | A69-9001-18617 | 355160029 | 911 | 985 | 6.11 | SBU |
| 2016-05-04 | ATS-422 | A69-9001-18611 | 355160019 | 913 | 1021 | 6.38 | SBU |
| 2016-05-04 | ATS-423 | A69-9001-18603 | 355160014 | 920 | 1040 | 6.60 | SBU |
| 2016-05-04 | ATS-424 | A69-9001-18623 | 355160028 | 859 | 977 | 6.65 | SBU |
| 2016-05-04 | ATS-425 | A69-9001-18608 | 355160020 | 910 | 1042 | 6.74 | SBU |
| 2016-05-04 | ATS-426 | A69-9001-18631 | 355160038 | 943 | 1070 | 6.75 | SBU |
| 2016-05-04 | ATS-427 | A69-9001-18614 | 355160022 | 919 | 1032 | 6.78 | SBU |
| 2016-05-04 | ATS-428 | A69-9001-18613 | 355160026 | 944 | 1070 | 7.01 | SBU |
| 2016-05-04 | ATS-429 | A69-9001-18616 | 355160027 | 926 | 1041 | 7.28 | SBU |
| 2016-05-04 | ATS-430 | A69-9001-18597 | 355160006 | 940 | 1060 | 7.41 | SBU |
| 2016-05-04 | ATS-431 | A69-9001-18621 | 355160031 | 944 | 1091 | 7.98 | SBU |
| 2016-05-04 | ATS-432 | A69-9001-18593 | 355160001 | 920 | 1060 | 8.26 | SBU |
| 2016-05-04 | ATS-433 | A69-9001-18592 | 355160003 | 985 | 1115 | 8.75 | SBU |
| 2016-05-04 | ATS-434 | A69-9001-18630 | 355160032 | 1020 | 1161 | 9.71 | SBU |
| 2016-05-04 | ATS-435 | A69-9001-18599 | 355160007 | 1030 | 1130 | 10.64 | SBU |
| 2016-05-04 | ATS-436 | A69-9001-18629 | 355160047 | 1082 | 1219 | 11.16 | SBU |
| 2016-05-04 | ATS-437 | A69-9001-18620 | 355160024 | 1153 | 1311 | 13.37 | SBU |
| 2017-05-09 | ATS-438 | A69-9001-14310 | 355170018 | 673 | 760 | 2.34 | SBU |
| 2017-05-09 | ATS-439 | A69-9001-14306 | 355170008 | 629 | 740 | 2.38 | SBU |
| 2017-05-09 | ATS-440 | A69-9001-14332 | 355170029 | 677 | 746 | 2.64 | SBU |
| 2017-05-09 | ATS-441 | A69-9001-14300 | 355170007 | 681 | 801 | 2.76 | SBU |
| 2017-05-09 | ATS-442 | A69-9001-14323 | 355170030 | 673 | 782 | 2.84 | SBU |
| 2017-05-09 | ATS-443 | A69-9001-14326 | 355170026 | 659 | 764 | 2.88 | SBU |
| 2017-05-09 | ATS-444 | A69-9001-14312 | 355170019 | 704 | 757 | 2.91 | SBU |
| 2017-05-09 | ATS-445 | A69-9001-14298 | 355170006 | 702 | 821 | 2.98 | SBU |
| 2017-05-09 | ATS-446 | A69-9001-14301 | 355170013 | 715 | 837 | 3.09 | SBU |
| 2017-05-09 | ATS-447 | A69-9001-14311 | 355170004 | 694 | 776 | 3.22 | SBU |
| 2017-05-09 | ATS-448 | A69-9001-14309 | 355170017 | 721 | 815 | 3.28 | SBU |
| 2017-05-09 | ATS-449 | A69-9001-14305 | 355170016 | 736 | 852 | 3.38 | SBU |
| 2017-05-09 | ATS-450 | A69-9001-14327 | 355170031 | 722 | 837 | 3.38 | SBU |
| 2017-05-09 | ATS-451 | A69-9001-14314 | 355170020 | 718 | 810 | 3.40 | SBU |
| 2017-05-09 | ATS-452 | A69-9001-14328 | 355170028 | 708 | 792 | 3.40 | SBU |
| 2017-05-09 | ATS-453 | A69-9001-14313 | 350150092 | 723 | 826 | 3.55 | SBU |
| 2017-05-09 | ATS-454 | A69-9001-14318 | 355170023 | 777 | 861 | 3.60 | SBU |
| 2017-05-09 | ATS-455 | A69-9001-14315 | 355170003 | 729 | 805 | 3.69 | SBU |
| 2017-05-09 | ATS-456 | A69-9001-14321 | 355170001 | 760 | 876 | 3.95 | SBU |
| 2017-05-09 | ATS-457 | A69-9001-14299 | 355170014 | 810 | 936 | 4.71 | SBU |
| 2017-05-09 | ATS-458 | A69-9001-14319 | 355170005 | 810 | 925 | 5.17 | SBU |
| 2017-05-09 | ATS-459 | A69-9001-14322 | 355170025 | 855 | 980 | 5.73 | SBU |
| 2017-05-09 | ATS-460 | A69-9001-14317 | 355170002 | 876 | 998 | 6.08 | SBU |
| 2017-05-09 | ATS-461 | A69-9001-14320 | 355170022 | 860 | 991 | 6.16 | SBU |
| 2017-05-09 | ATS-462 | A69-9001-14308 | 355170011 | 890 | 1014 | 6.40 | SBU |
| 2017-05-09 | ATS-463 | A69-9001-14307 | 355170015 | 885 | 1022 | 7.08 | SBU |
| 2017-05-09 | ATS-464 | A69-9001-14303 | 355170012 | 949 | 1056 | 7.92 | SBU |
| 2017-05-09 | ATS-465 | A69-9001-14304 | 355170009 | 935 | 1075 | 7.93 | SBU |
| 2017-05-09 | ATS-466 | A69-9001-14324 | 355170024 | 974 | 1108 | 8.84 | SBU |
| 2017-05-09 | ATS-467 | A69-9001-14302 | 355170010 | 955 | 1100 | 9.19 | SBU |
| 2017-05-09 | ATS-468 | A69-9001-14325 | 355170032 | 1017 | 1160 | 10.88 | SBU |
| 2017-05-09 | ATS-469 | A69-9001-14316 | 355170021 | 1102 | 1231 | 13.40 | SBU |
| 2017-05-09 | ATS-470 | A69-9001-14330 | 355170027 | 1620 | 1832 | 38.12 | SBU |
| 2017-05-10 | ATS-471 | A69-9001-14336 | 355170042 | 656 | 753 | 2.32 | SBU |
| 2017-05-10 | ATS-472 | A69-9001-14351 | 355170059 | 654 | 739 | 2.36 | SBU |
| 2017-05-10 | ATS-473 | A69-9001-14344 | 355170045 | 734 | 842 | 2.78 | SBU |
| 2017-05-10 | ATS-474 | A69-9001-14345 | 355170039 | 655 | 814 | 2.80 | SBU |
| 2017-05-10 | ATS-475 | A69-9001-14341 | 355170044 | 690 | 795 | 2.92 | SBU |
| 2017-05-10 | ATS-476 | A69-9001-14349 | 355170058 | 709 | 825 | 3.04 | SBU |
| 2017-05-10 | ATS-477 | A69-9001-14340 | 355170046 | 692 | 782 | 3.16 | SBU |
| 2017-05-10 | ATS-478 | A69-9001-14357 | 355170057 | 726 | 838 | 3.46 | SBU |
| 2017-05-10 | ATS-479 | A69-9001-14343 | 355170038 | 754 | 863 | 3.54 | SBU |
| 2017-05-10 | ATS-480 | A69-9001-14352 | 355170050 | 714 | 810 | 3.56 | SBU |
| 2017-05-10 | ATS-481 | A69-9001-14347 | 355170054 | 770 | 892 | 3.82 | SBU |
| 2017-05-10 | ATS-482 | A69-9001-14353 | 355170055 | 765 | 904 | 3.84 | SBU |
| 2017-05-10 | ATS-483 | A69-9001-14329 | 355170034 | 775 | 901 | 4.08 | SBU |
| 2017-05-10 | ATS-484 | A69-9001-14333 | 355170033 | 790 | 875 | 4.20 | SBU |
| 2017-05-10 | ATS-485 | A69-9001-14342 | 355170047 | 781 | 890 | 4.44 | SBU |
| 2017-05-10 | ATS-486 | A69-9001-14354 | 355170052 | 820 | 937 | 5.06 | SBU |
| 2017-05-10 | ATS-487 | A69-9001-14346 | 355170048 | 851 | 942 | 5.40 | SBU |
| 2017-05-10 | ATS-488 | A69-9001-14358 | 350120209 | 862 | 1002 | 6.50 | SBU |
| 2017-05-10 | ATS-489 | A69-9001-14356 | 355170053 | 923 | 1048 | 6.50 | SBU |
| 2017-05-10 | ATS-490 | A69-9001-14339 | 355170037 | 872 | 934 | 6.52 | SBU |
| 2017-05-10 | ATS-491 | A69-9001-14348 | 355170049 | 949 | 1070 | 7.68 | SBU |
| 2017-05-10 | ATS-492 | A69-9001-14334 | 355170040 | 972 | 1126 | 8.56 | SBU |
| 2017-05-10 | ATS-493 | A69-9001-14335 | 355170041 | 980 | 1130 | 8.66 | SBU |
| 2017-05-10 | ATS-494 | A69-9001-14331 | 355170035 | 1059 | 1234 | 11.04 | SBU |
| 2017-05-10 | ATS-495 | A69-9001-14350 | 355170051 | 1124 | 1279 | 13.92 | SBU |
| 2017-05-10 | ATS-496 | A69-9001-14338 | 355170043 | 1228 | 1405 | 16.86 | SBU |
| 2017-05-10 | ATS-497 | A69-9001-14337 | 355170036 | 2050 | 2320 | 77.00 | SBU |
| 2017-05-11 | ATS-498 | A69-9001-14370 | 355170075 | 620 | 715 | 1.72 | SBU |
| 2017-05-11 | ATS-499 | A69-9001-14371 | 355170071 | 650 | 739 | 2.49 | SBU |
| 2017-05-11 | ATS-500 | A69-9001-14365 | 355170066 | 619 | 751 | 2.73 | SBU |
| 2017-05-11 | ATS-501 | A69-9001-14379 | 355170079 | 696 | 776 | 2.82 | SBU |
| 2017-05-11 | ATS-502 | A69-9001-14360 | 355170061 | 682 | 787 | 2.89 | SBU |
| 2017-05-11 | ATS-503 | A69-9001-14364 | 355170065 | 715 | 834 | 3.67 | SBU |
| 2017-05-11 | ATS-504 | A69-9001-14367 | 355170068 | 734 | 829 | 3.70 | SBU |
| 2017-05-11 | ATS-505 | A69-9001-14359 | 355170060 | 808 | 954 | 4.15 | SBU |
| 2017-05-11 | ATS-506 | A69-9001-14366 | 355170067 | 803 | 920 | 4.26 | SBU |
| 2017-05-11 | ATS-507 | A69-9001-14374 | 355170074 | 789 | 921 | 4.28 | SBU |
| 2017-05-11 | ATS-508 | A69-9001-14362 | 355170063 | 770 | 880 | 4.32 | SBU |
| 2017-05-11 | ATS-509 | A69-9001-14363 | 355170064 | 778 | 909 | 4.36 | SBU |
| 2017-05-11 | ATS-510 | A69-9001-14361 | 355170062 | 802 | 942 | 4.75 | SBU |
| 2017-05-11 | ATS-511 | A69-9001-14373 | 355170073 | 813 | 892 | 4.83 | SBU |
| 2017-05-11 | ATS-512 | A69-9001-14368 | 355170069 | 843 | 952 | 4.98 | SBU |
| 2017-05-11 | ATS-513 | A69-9001-14377 | 355170078 | 888 | 1029 | 5.48 | SBU |
| 2017-05-11 | ATS-514 | A69-9001-14376 | 355170077 | 880 | 1017 | 6.24 | SBU |
| 2017-05-11 | ATS-515 | A69-9001-14375 | 355170076 | 868 | 1011 | 6.29 | SBU |
| 2017-05-11 | ATS-516 | A69-9001-14369 | 355170070 | 915 | 1090 | 6.72 | SBU |
| 2017-05-11 | ATS-517 | A69-9001-14372 | 355170072 | 963 | 1201 | 8.50 | SBU |
| 2017-10-04 | ATS-518 | A69-9001-14393 | 355170080 | 1236 | 1376 | 15.71 | SBU |
| 2017-10-10 | ATS-519 | A69-9001-13192 | 355170081 | 952 | 1031 | 6.42 | SBU |
| 2017-10-10 | ATS-520 | A69-9001-14378 | 355170082 | 1020 | 1188 | 8.20 | SBU |
| 2017-10-10 | ATS-521 | A69-9001-14380 | 355170083 | 1278 | 1474 | 17.14 | SBU |
| 2017-10-11 | ATS-522 | A69-9001-14388 | 355170086 | 917 | 1055 | 5.11 | SBU |
| 2017-10-11 | ATS-523 | A69-9001-14386 | 355170084 | 880 | 1039 | 5.28 | SBU |
| 2017-10-11 | ATS-524 | A69-9001-14387 | 355170088 | 896 | 1031 | 5.47 | SBU |
| 2017-10-11 | ATS-525 | A69-9001-14390 | 350150134 | 904 | 1031 | 5.54 | SBU |
| 2017-10-11 | ATS-526 | A69-9001-14392 | 355170090 | 1026 | 1161 | 8.06 | SBU |
| 2017-10-11 | ATS-527 | A69-9001-14389 | 355170087 | 1025 | 1190 | 8.31 | SBU |
| 2017-10-11 | ATS-528 | A69-9001-14391 | 355170089 | 1071 | 1245 | 10.37 | SBU |
| 2017-10-11 | ATS-529 | A69-9001-14385 | 355170085 | 1044 | 1227 | 10.53 | SBU |
| 2018-05-09 | ATS-530 | A69-9001-10095 | 355180001 | 778 | 872 | 4.12 | SBU |
| 2018-05-09 | ATS-531 | A69-9001-10101 | 355180007 | 791 | 919 | 4.14 | SBU |
| 2018-05-09 | ATS-532 | A69-9001-10097 | 355180003 | 820 | 937 | 4.60 | SBU |
| 2018-05-09 | ATS-533 | A69-9001-10099 | 355180005 | 844 | 975 | 4.86 | SBU |
| 2018-05-09 | ATS-534 | A69-9001-10102 | 355180008 | 934 | 1073 | 7.22 | SBU |
| 2018-05-09 | ATS-535 | A69-9001-10103 | 355180009 | 1020 | 1148 | 9.28 | SBU |
| 2018-05-09 | ATS-536 | A69-9001-10096 | 355180002 | 1068 | 1181 | 10.29 | SBU |
| 2018-05-09 | ATS-537 | A69-9001-10100 | 355180006 | 1170 | 1346 | 13.48 | SBU |
| 2018-05-09 | ATS-538 | A69-9001-10098 | 355180004 | 1356 | 1549 | 22.24 | SBU |
| 2018-05-10 | ATS-539 | A69-9001-10109 | 355180015 | 693 | 763 | 2.69 | SBU |
| 2018-05-10 | ATS-540 | A69-9001-10110 | 355180016 | 750 | 831 | 3.29 | SBU |
| 2018-05-10 | ATS-541 | A69-9001-10113 | 355180019 | 811 | 931 | 4.50 | SBU |
| 2018-05-10 | ATS-542 | A69-9001-10107 | 355180013 | 798 | 881 | 4.51 | SBU |
| 2018-05-10 | ATS-543 | A69-9001-10108 | 355180014 | 823 | 902 | 4.75 | SBU |
| 2018-05-10 | ATS-544 | A69-9001-10112 | 355180018 | 849 | 971 | 5.12 | SBU |
| 2018-05-10 | ATS-545 | A69-9001-10104 | 355180010 | 876 | 992 | 5.81 | SBU |
| 2018-05-10 | ATS-546 | A69-9001-10106 | 355180012 | 876 | 1003 | 5.97 | SBU |
| 2018-05-10 | ATS-547 | A69-9001-10105 | 355180011 | 985 | 1107 | 8.28 | SBU |
| 2018-05-10 | ATS-548 | A69-9001-10114 | 355180020 | 984 | 1118 | 8.90 | SBU |
| 2018-05-10 | ATS-549 | A69-9001-10111 | 355180017 | 1561 | 1784 | 38.62 | SBU |
| 2018-05-11 | ATS-550 | A69-9001-10118 | 355180024 | 667 | 782 | 2.14 | SBU |
| 2018-05-11 | ATS-551 | A69-9001-10119 | 355180025 | 739 | 836 | 3.60 | SBU |
| 2018-05-11 | ATS-552 | A69-9001-10117 | 355180023 | 920 | 1039 | 7.31 | SBU |
| 2018-05-11 | ATS-553 | A69-9001-10116 | 355180022 | 1020 | 1175 | 9.81 | SBU |
| 2019-07-24 | ATS-554 | A69-9001-10124 | - | 1640 | 1840 | 42.00 | SBU/DSU |
| 2019-07-24 | ATS-555 | A69-9001-10132 | - | 1770 | 2000 | 55.00 | SBU/DSU |
| 2019-07-24 | ATS-556 | A69-9001-10125 | - | 1810 | 2030 | 58.00 | SBU/DSU |
| 2019-07-24 | ATS-557 | A69-9001-10131 | - | 1870 | 2110 | 66.00 | SBU/DSU |
| 2019-07-25 | ATS-558 | A69-9001-10126 | - | 1540 | 1730 | 37.00 | SBU/DSU |
| 2019-07-25 | ATS-559 | A69-9001-10120 | - | 1610 | 1810 | 41.00 | SBU/DSU |
| 2019-07-25 | ATS-560 | A69-9001-10121 | - | 1610 | 1860 | 42.00 | SBU/DSU |
| 2019-07-26 | ATS-561 | A69-9001-10128 | - | 1290 | 1420 | 28.00 | SBU/DSU |
| 2019-07-26 | ATS-562 | A69-9001-10130 | - | 1400 | 1620 | 29.00 | SBU/DSU |
| 2019-07-26 | ATS-563 | A69-9001-10123 | - | 1580 | 1750 | 39.00 | SBU/DSU |
| 2019-07-26 | ATS-564 | A69-9001-10122 | - | 1620 | 1840 | 46.00 | SBU/DSU |
| 2019-07-26 | ATS-565 | A69-9001-12724 | - | 1680 | 1910 | 49.00 | SBU/DSU |
| 2019-07-26 | ATS-566 | A69-9001-10133 | - | 1640 | 1890 | 52.00 | SBU/DSU |
| 2019-07-26 | ATS-567 | A69-9001-10129 | - | 1790 | 2020 | 61.00 | SBU/DSU |
| 2019-07-26 | ATS-568 | A69-9001-10127 | - | 1860 | 2120 | 69.00 | SBU/DSU |
| 2019-07-26 | ATS-569 | A69-9001-10134 | - | 1920 | 2080 | 73.00 | SBU/DSU |
| - | ATS-570 | A69-1602-30516 | - | - | - | - | MU |
| - | ATS-571 | A69-1602-30519 | - | - | - | - | MU |
| - | ATS-572 | A69-1602-30521 | - | - | - | - | MU |
| - | ATS-573 | A69-1602-30522 | - | - | - | - | MU |
| - | ATS-574 | A69-1602-30524 | - | - | - | - | MU |
| - | ATS-575 | A69-1602-30526 | - | - | - | - | MU |
| - | ATS-576 | A69-1602-30528 | - | - | - | - | MU |
| - | ATS-577 | A69-1602-30530 | - | - | - | - | MU |
| - | ATS-578 | A69-1602-30532 | - | - | - | - | MU |
| - | ATS-579 | A69-1602-30537 | - | - | - | - | MU |
| - | ATS-580 | A69-1602-30539 | - | - | - | - | MU |
| - | ATS-581 | A69-1602-30546 | - | - | - | - | MU |
| - | ATS-582 | A69-1602-30550 | - | - | - | - | MU |
| - | ATS-583 | A69-1602-30551 | - | - | - | - | MU |
| - | ATS-584 | A69-1602-30552 | - | - | - | - | MU |
| - | ATS-585 | A69-1602-30564 | - | - | - | - | MU |
| - | ATS-586 | A69-9001-28889 | - | - | - | - | MU |
| - | ATS-587 | A69-9001-28890 | - | - | - | - | MU |
| - | ATS-588 | A69-9001-28891 | - | - | - | - | MU |
| - | ATS-589 | A69-9001-28892 | - | - | - | - | MU |
| - | ATS-590 | A69-9001-28893 | - | - | - | - | MU |
| - | ATS-591 | A69-9001-28894 | - | - | - | - | MU |
| - | ATS-592 | A69-9001-28895 | - | - | - | - | MU |
| - | ATS-593 | A69-9001-28896 | - | - | - | - | MU |
| - | ATS-594 | A69-9001-28897 | - | - | - | - | MU |
| - | ATS-595 | A69-9001-28898 | - | - | - | - | MU |
| - | ATS-596 | A69-9001-28899 | - | - | - | - | MU |
| - | ATS-597 | A69-9001-28900 | - | - | - | - | MU |
| - | ATS-598 | A69-9001-28903 | - | - | - | - | MU |
| - | ATS-599 | A69-9001-28904 | - | - | - | - | MU |

**Table S3.** Sources and data contributors for coastwide acoustic telemetry detections of Atlantic Sturgeon.

| General location | Organization | Data provider(s) |
| --- | --- | --- |
| Maine | National Oceanic and Atmospheric Association | J. Hawkes |
| Massachusetts | Massachusetts Division of Marine Fisheries<br>National Oceanic and Atmospheric Association<br>United States Geological Survey | B. Gahagan; B. Hoffman; M. Kieffer<br>T. Rowell<br>M. Winton |
| Rhode Island | Atlantic Shark Institute<br>INSPIRE Environmental<br>Rhode Island Department of Environmental Management | J. Dodd<br>B. Gervelis<br>C. McManus; E. Schneider |
| Connecticut | Connecticut Department of Energy and Environmental Protection | D. Pacileo |
| New York | Stony Brook University | S. Cernadas-Martin; B. Scannell |
| New Jersey | National Oceanic and Atmospheric Association<br>New England Aquarium<br>Rutgers University<br>University of Delaware | G. Goulette<br>J. Kneebone<br>L. Nazzaro<br>D. Haulsee |
| Delaware | Delaware Division of Fish and Wildlife<br>Delaware State University | I. Park<br>D. Fox |
| Maryland | University of Maryland Center for Environmental Science | M. O'Brien; E. Rothermel |
| Virginia | National Oceanic and Atmospheric Association<br>United States Department of the Navy<br>Virginia Institute of Marine Science | M. Bromilow; W. Cruz-Marrero<br>C. Watterson<br>D. Crear |
| North Carolina | East Carolina University | J. Luczkovich |
|  | New Jersey Department of Environmental Protection | J. Cimino |
|  | North Carolina Department of Environment and Natural Resources | R. Gregory; S. Poland |
|  | North Carolina State University | J. Buckel; R. Gallagher; J. Krause; |
|  | South Carolina Department of Natural Resources | T. Darden |
|  | South-East Zoo Alliance for Reproduction and Conservation | N. Pham Ho |
| South Carolina | South Carolina Department of Natural Resources | M. Arendt; B. Post |
| Georgia | Georgia Department of Natural Resources | C. Kalinowsky |
| Florida | Georgia Aquarium | E. Alberts |

**Table S4.**
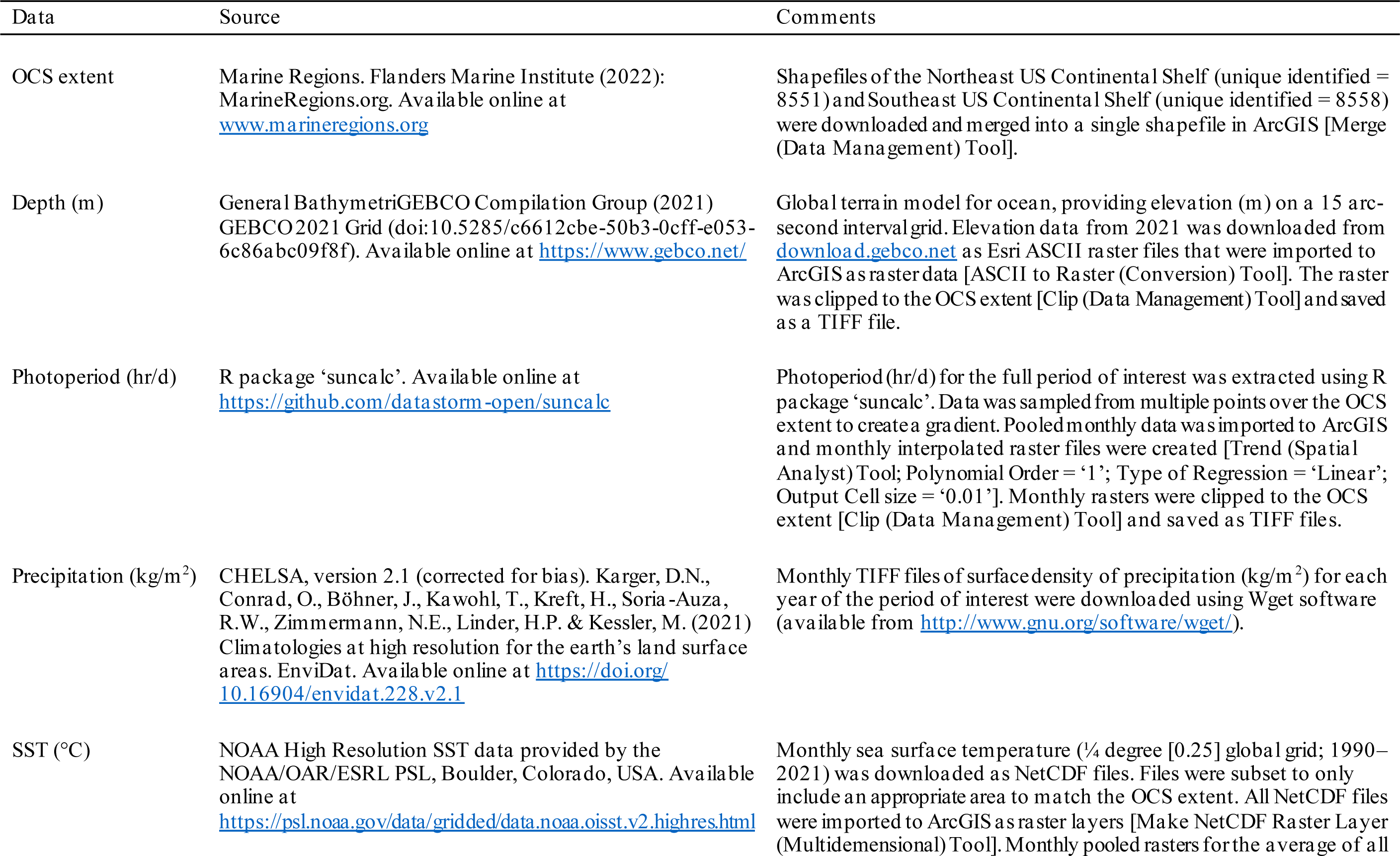
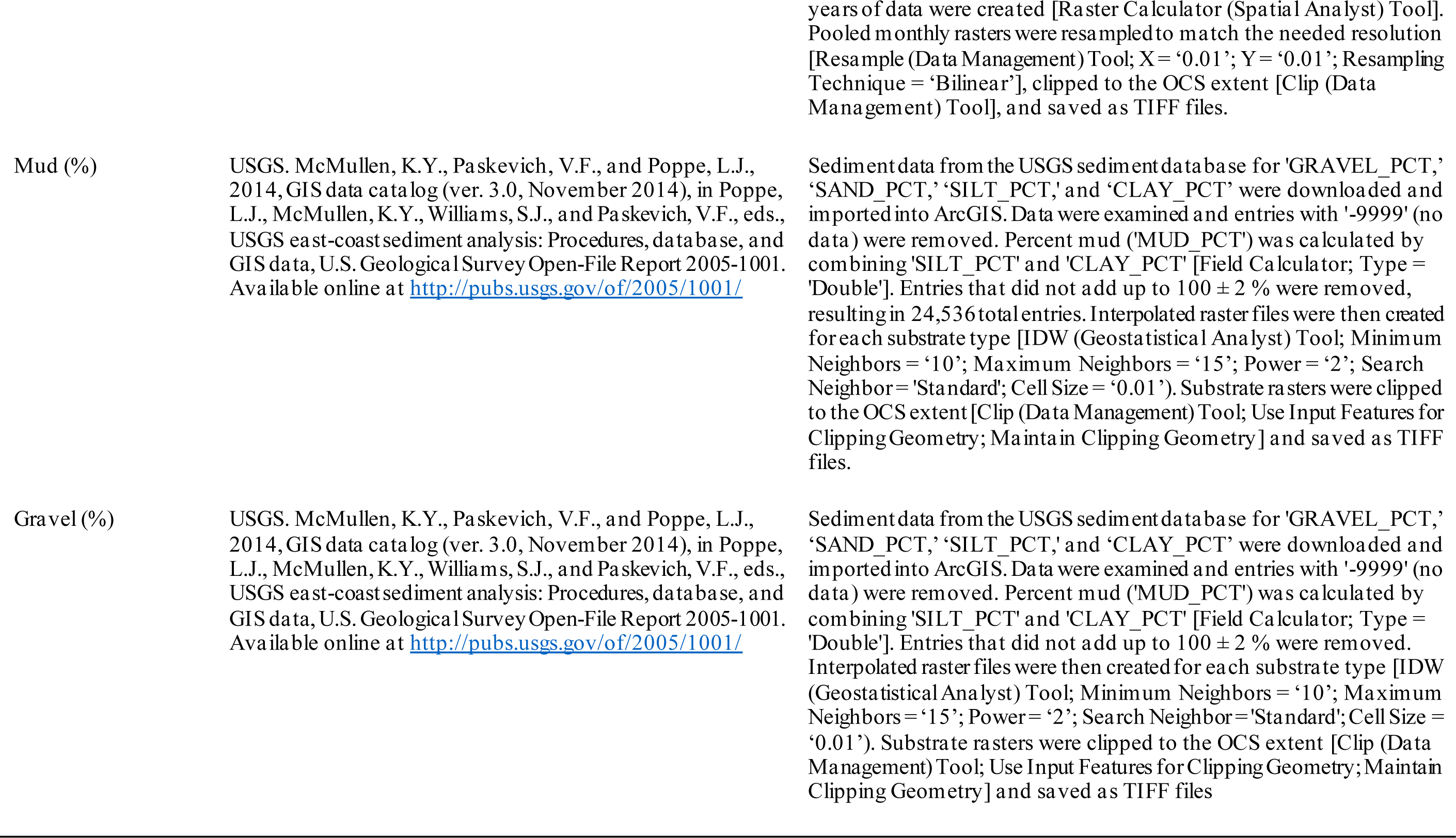
Sources of environmental data used to develop species distribution models.

**Table S5.**
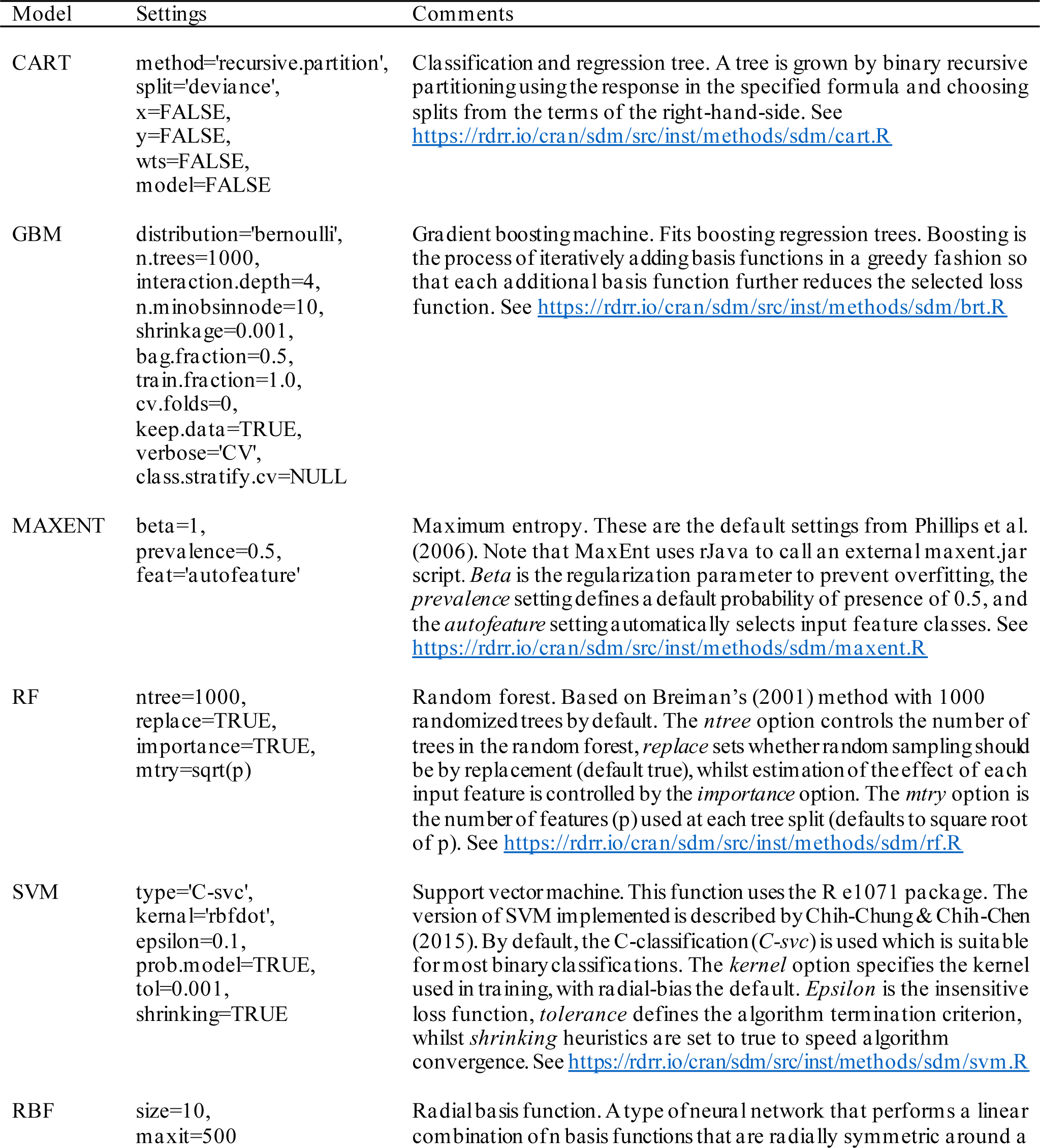
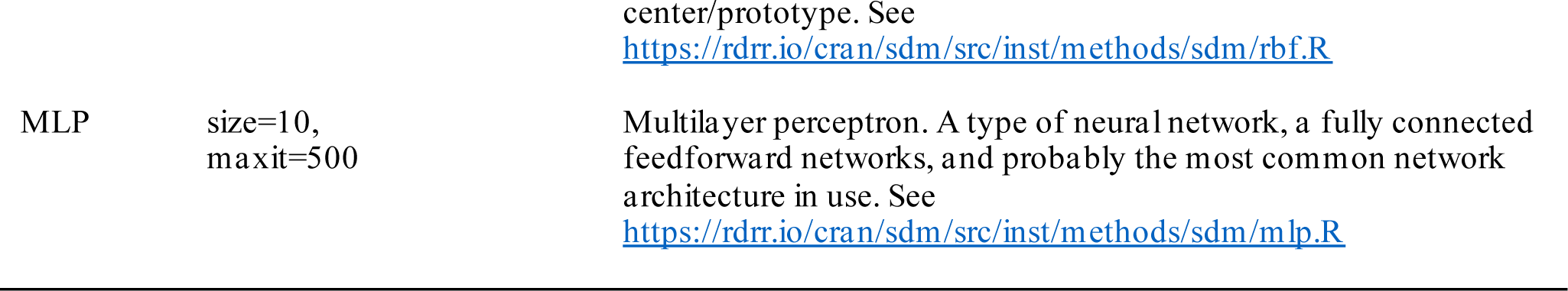
Summary of artificial intelligence algorithms and settings used to fit species distribution models. The default settings for tuning and regularization of parameters were used. Further details of default settings used for all available models are available at https://rdrr.io/cran/sdm/f/.

**Table S6.** Overall mean [SD] model performance for all model types across each month. AUC = area under the receiver operating curve; TSS = true skill statistic; Kappa = Cohen’s kappa; CART = classification and regression tree; GBM = gradient boosting machine; MAXENT = maximum entropy; RF = random forest; SVM = support vector machine; MLP = multilayer perceptron; RBF = radial basis function.

|  | Model | AUC<sup>a</sup> | TSS<sup>b</sup> | Kappa<sup>c</sup> |
| --- | --- | --- | --- | --- |
| January | CART | 0.938 [0.026] | 0.809 [0.062] | 0.695 [0.088] |
|  | GBM | 0.968 [0.010] | 0.831 [0.038] | 0.720 [0.058] |
|  | MAXENT | 0.966 [0.008] | 0.835 [0.035] | 0.727 [0.050] |
|  | RF | 0.992 [0.005] | 0.920 [0.028] | 0.857 [0.046] |
|  | SVM | 0.928 [0.017] | 0.701 [0.051] | 0.552 [0.061] |
|  | MLP | 0.952 [0.014] | 0.804 [0.051] | 0.685 [0.068] |
|  | RBF | 0.853 [0.143] | 0.590 [0.242] | 0.456 [0.174] |
| February | CART | 0.939 [0.031] | 0.811 [0.067] | 0.700 [0.091] |
|  | GBM | 0.971 [0.009] | 0.832 [0.046] | 0.722 [0.069] |
|  | MAXENT | 0.968 [0.008] | 0.849 [0.035] | 0.747 [0.053] |
|  | RF | 0.994 [0.003] | 0.937 [0.027] | 0.884 [0.046] |
|  | SVM | 0.928 [0.018] | 0.716 [0.048] | 0.570 [0.058] |
|  | MLP | 0.954 [0.017] | 0.813 [0.054] | 0.696 [0.070] |
|  | RBF | 0.873 [0.115] | 0.607 [0.206] | 0.466 [0.154] |
| March | CART | 0.934 [0.024] | 0.818 [0.052] | 0.732 [0.068] |
|  | GBM | 0.967 [0.008] | 0.828 [0.033] | 0.742 [0.045] |
|  | MAXENT | 0.969 [0.008] | 0.850 [0.029] | 0.771 [0.043] |
|  | RF | 0.992 [0.003] | 0.919 [0.029] | 0.871 [0.044] |
|  | SVM | 0.946 [0.013] | 0.778 [0.040] | 0.676 [0.052] |
|  | MLP | 0.959 [0.012] | 0.830 [0.040] | 0.745 [0.057] |
|  | RBF | 0.883 [0.067] | 0.619 [0.120] | 0.498 [0.108] |
| April | CART | 0.948 [0.023] | 0.851 [0.048] | 0.781 [0.069] |
|  | GBM | 0.978 [0.008] | 0.871 [0.038] | 0.801 [0.054] |
|  | MAXENT | 0.983 [0.006] | 0.891 [0.028] | 0.829 [0.041] |
|  | RF | 0.994 [0.004] | 0.939 [0.026] | 0.901 [0.039] |
|  | SVM | 0.952 [0.011] | 0.781 [0.048] | 0.679 [0.061] |
|  | MLP | 0.963 [0.010] | 0.833 [0.035] | 0.747 [0.050] |
|  | RBF | 0.872 [0.121] | 0.645 [0.210] | 0.537 [0.172] |
| May | CART | 0.948 [0.021] | 0.856 [0.038] | 0.812 [0.045] |
|  | GBM | 0.975 [0.008] | 0.859 [0.032] | 0.810 [0.041] |
|  | MAXENT | 0.977 [0.006] | 0.863 [0.027] | 0.815 [0.033] |
|  | RF | 0.993 [0.004] | 0.942 [0.017] | 0.919 [0.023] |
|  | SVM | 0.964 [0.008] | 0.836 [0.027] | 0.781 [0.034] |
|  | MLP | 0.969 [0.010] | 0.878 [0.034] | 0.834 [0.043] |
|  | RBF | 0.893 [0.059] | 0.646 [0.112] | 0.565 [0.119] |
| June | CART | 0.938 [0.023] | 0.823 [0.067] | 0.765 [0.094] |
|  | GBM | 0.972 [0.008] | 0.840 [0.035] | 0.770 [0.049] |
|  | MAXENT | 0.978 [0.007] | 0.886 [0.027] | 0.832 [0.036] |
|  | RF | 0.994 [0.003] | 0.941 [0.020] | 0.911 [0.029] |
|  | SVM | 0.975 [0.008] | 0.884 [0.028] | 0.830 [0.038] |
|  | MLP | 0.978 [0.007] | 0.907 [0.023] | 0.861 [0.034] |
|  | RBF | 0.919 [0.121] | 0.725 [0.214] | 0.645 [0.189] |
| July | CART | 0.945 [0.027] | 0.856 [0.042] | 0.765 [0.066] |
|  | GBM | 0.981 [0.007] | 0.876 [0.034] | 0.758 [0.060] |
|  | MAXENT | 0.988 [0.005] | 0.915 [0.026] | 0.828 [0.054] |
|  | RF | 0.996 [0.003] | 0.946 [0.024] | 0.885 [0.048] |
|  | SVM | 0.977 [0.010] | 0.858 [0.039] | 0.730 [0.069] |
|  | MLP | 0.984 [0.007] | 0.917 [0.034] | 0.831 [0.063] |
|  | RBF | 0.892 [0.111] | 0.667 [0.202] | 0.500 [0.174] |
| August | CART | 0.955 [0.040] | 0.892 [0.070] | 0.768 [0.114] |
|  | GBM | 0.989 [0.010] | 0.925 [0.047] | 0.780 [0.109] |
|  | MAXENT | 0.986 [0.014] | 0.920 [0.041] | 0.762 [0.090] |
|  | RF | 0.996 [0.007] | 0.958 [0.046] | 0.864 [0.115] |
|  | SVM | 0.976 [0.020] | 0.870 [0.066] | 0.661 [0.117] |
|  | MLP | 0.983 [0.014] | 0.903 [0.064] | 0.733 [0.132] |
|  | RBF | 0.934 [0.059] | 0.753 [0.120] | 0.482 [0.127] |
| September | CART | 0.947 [0.035] | 0.875 [0.062] | 0.803 [0.088] |
|  | GBM | 0.984 [0.014] | 0.904 [0.044] | 0.776 [0.089] |
|  | MAXENT | 0.991 [0.004] | 0.925 [0.026] | 0.819 [0.056] |
|  | RF | 0.998 [0.002] | 0.965 [0.033] | 0.908 [0.070] |
|  | SVM | 0.977 [0.010] | 0.868 [0.041] | 0.708 [0.079] |
|  | MLP | 0.988 [0.007] | 0.931 [0.042] | 0.833 [0.086] |
|  | RBF | 0.910 [0.137] | 0.696 [0.236] | 0.492 [0.154] |
| October | CART | 0.932 [0.030] | 0.829 [0.068] | 0.754 [0.107] |
|  | GBM | 0.982 [0.007] | 0.882 [0.030] | 0.792 [0.046] |
|  | MAXENT | 0.983 [0.006] | 0.887 [0.030] | 0.798 [0.048] |
|  | RF | 0.995 [0.003] | 0.942 [0.024] | 0.892 [0.041] |
|  | SVM | 0.970 [0.010] | 0.839 [0.037] | 0.724 [0.057] |
|  | MLP | 0.978 [0.010] | 0.887 [0.041] | 0.799 [0.064] |
|  | RBF | 0.902 [0.131] | 0.676 [0.220] | 0.540 [0.181] |
| November | CART | 0.946 [0.024] | 0.825 [0.049] | 0.757 [0.068] |
|  | GBM | 0.973 [0.009] | 0.841 [0.035] | 0.756 [0.048] |
|  | MAXENT | 0.974 [0.008] | 0.857 [0.038] | 0.779 [0.056] |
|  | RF | 0.992 [0.004] | 0.921 [0.030] | 0.872 [0.045] |
|  | SVM | 0.946 [0.020] | 0.797 [0.045] | 0.697 [0.060] |
|  | MLP | 0.964 [0.013] | 0.848 [0.044] | 0.767 [0.062] |
|  | RBF | 0.917 [0.114] | 0.716 [0.198] | 0.614 [0.168] |
| December | CART | 0.946 [0.018] | 0.828 [0.040] | 0.789 [0.048] |
|  | GBM | 0.968 [0.007] | 0.815 [0.036] | 0.772 [0.041] |
|  | MAXENT | 0.967 [0.007] | 0.824 [0.027] | 0.783 [0.033] |
|  | RF | 0.993 [0.003] | 0.927 [0.026] | 0.907 [0.032] |
|  | SVM | 0.941 [0.015] | 0.753 [0.050] | 0.701 [0.055] |
|  | MLP | 0.954 [0.011] | 0.828 [0.034] | 0.787 [0.040] |
|  | RBF | 0.887 [0.086] | 0.641 [0.150] | 0.584 [0.127] |

<sup>a</sup> Area under the receiver operating curve (AUC) a threshold-independent measure of model performance. It is a measure of the area under the curve of a receiver operating plot, which displays the true positive rate (i.e., sensitivity) against the false positive rate ( $1 - \text{specificity}$ ; where specificity is the proportion of negatives that can be correctly identified). Receiver operating curves are constructed by using all possible thresholds to classify the scores into confusion matrices, obtaining sensitivity and specificity for each matrix, and then plotting sensitivity against the corresponding proportion of false positives (equal to $1 - \text{specificity}$ ; Fielding and Bell, 1997).
<sup>b</sup> True skill statistic is a measure of model performance that aims to be relatively simple, while minimizing any biases in the assessment of the model that may arise from species prevalence. TSS is calculated as $\text{sensitivity} + \text{specificity} - 1$ (Allouche et al., 2006).
<sup>c</sup> Cohen's Kappa is a widely used measure to compare observations and predictions, and varies from $-1$ to $1$ , with values of $0$ or less = no better than random, and values of $1$ = perfect agreement (Cohen, 1960).

**Table S7.**
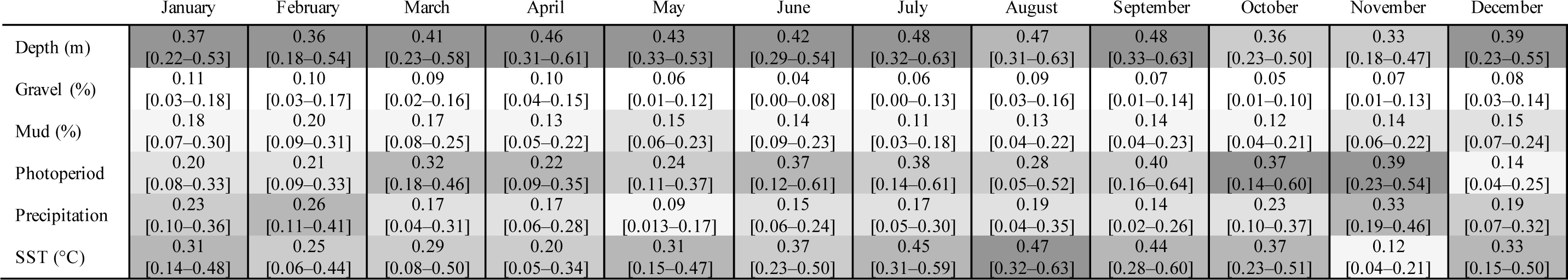
Table of predictor importance for all months. Grey shading scales with variable importance, with darker shades representing further greater importance. Summary graphs indicating variable importance, measured as 1 - Pearson correlation, averaged across all SDM models [95% confidence intervals].

**Figure S1.**
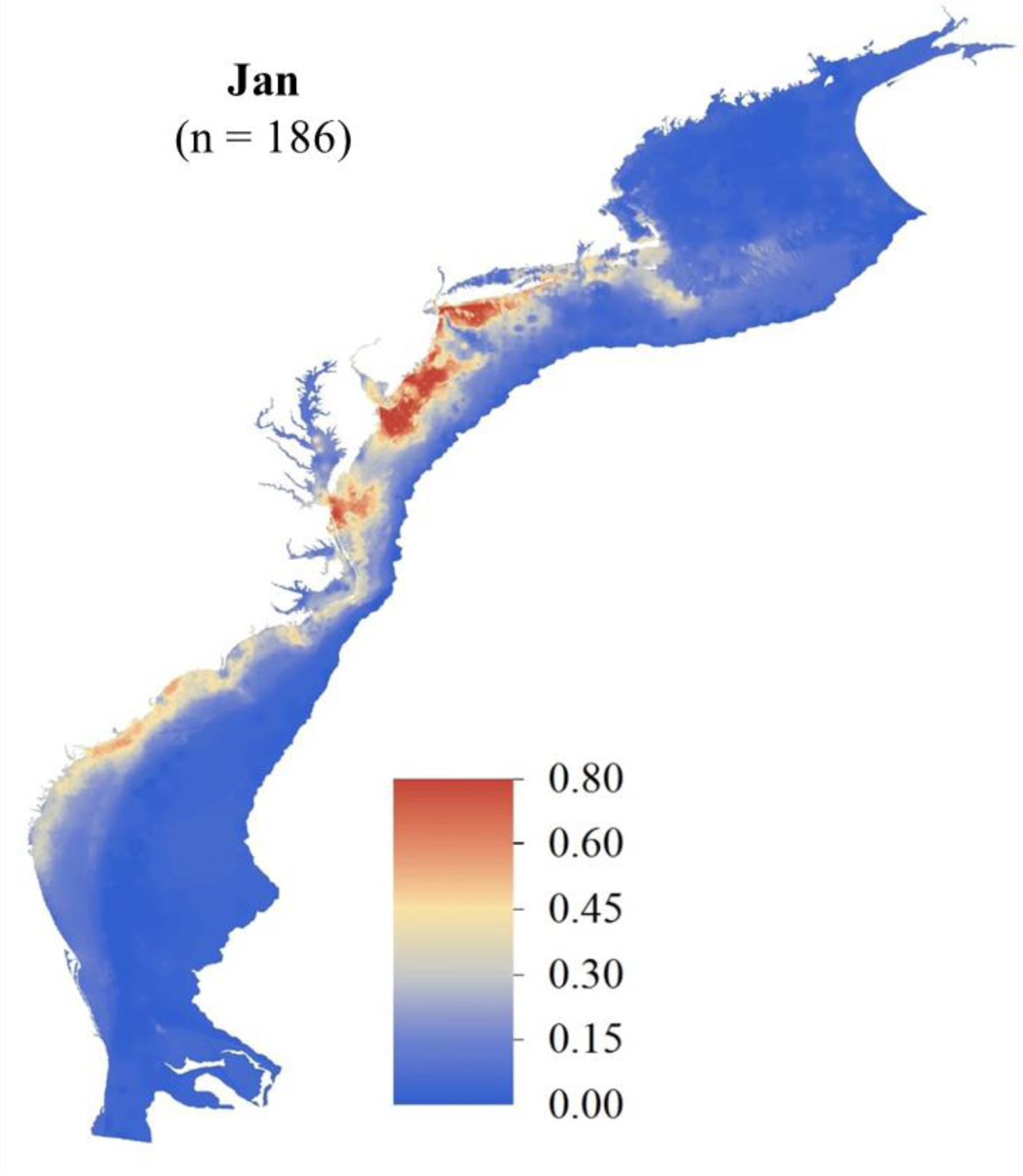
January SDM ensemble map of predicted Atlantic Sturgeon occurrence probabilities in the OCS (raster resolution = 1-km; total area = 626,204 km^2^). Count of total unique detection locations across the OCS is indicated.

**Figure S2.**
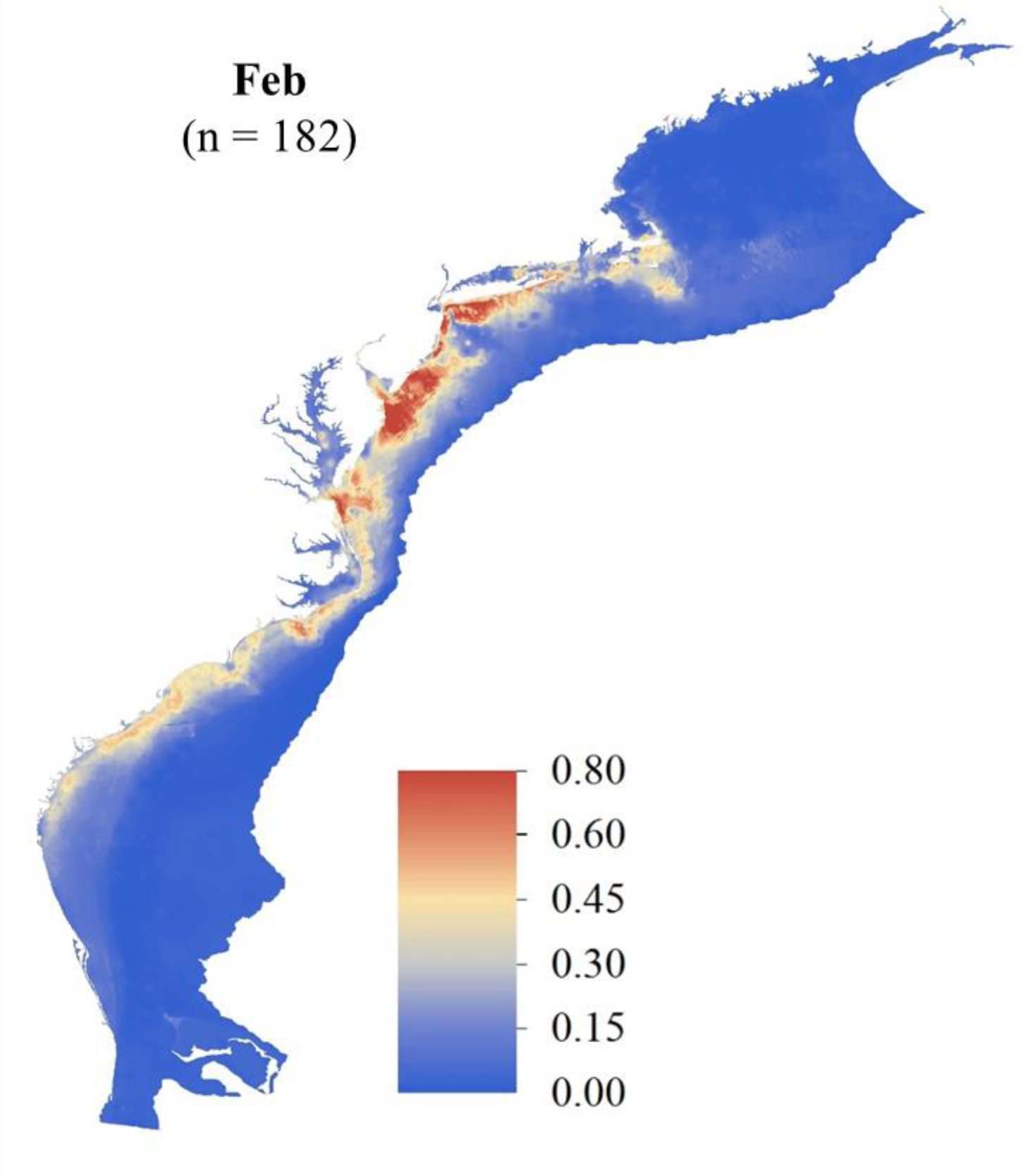
February SDM ensemble map of predicted Atlantic Sturgeon occurrence probabilities in the OCS (raster resolution = 1-km; total area = 626,204 km^2^). Count of total unique detection locations across the OCS is indicated.

**Figure S3.**
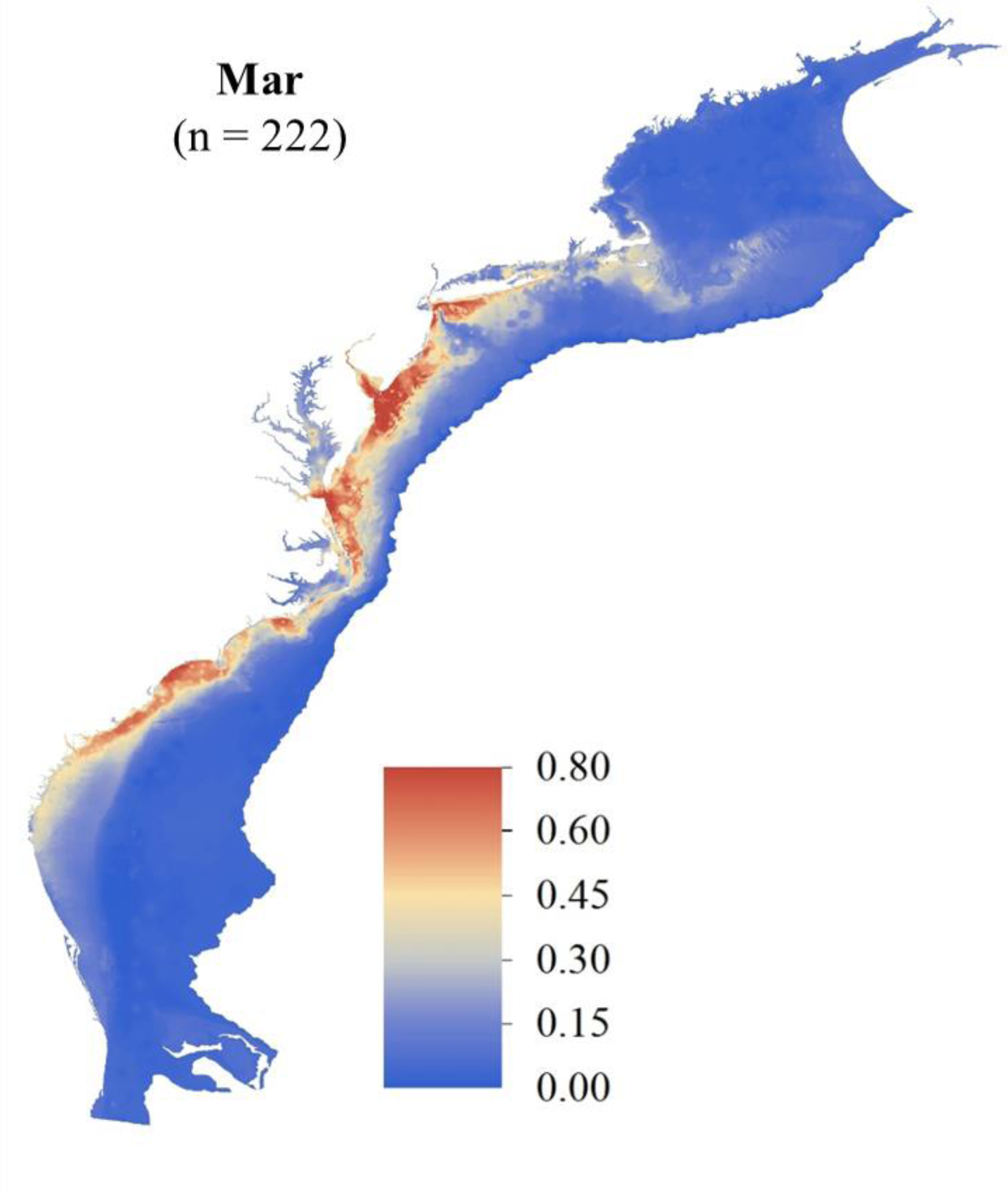
March SDM ensemble map of predicted Atlantic Sturgeon occurrence probabilities in the OCS (raster resolution = 1-km; total area = 626,204 km^2^). Count of total unique detection locations across the OCS is indicated.

**Figure S4.**
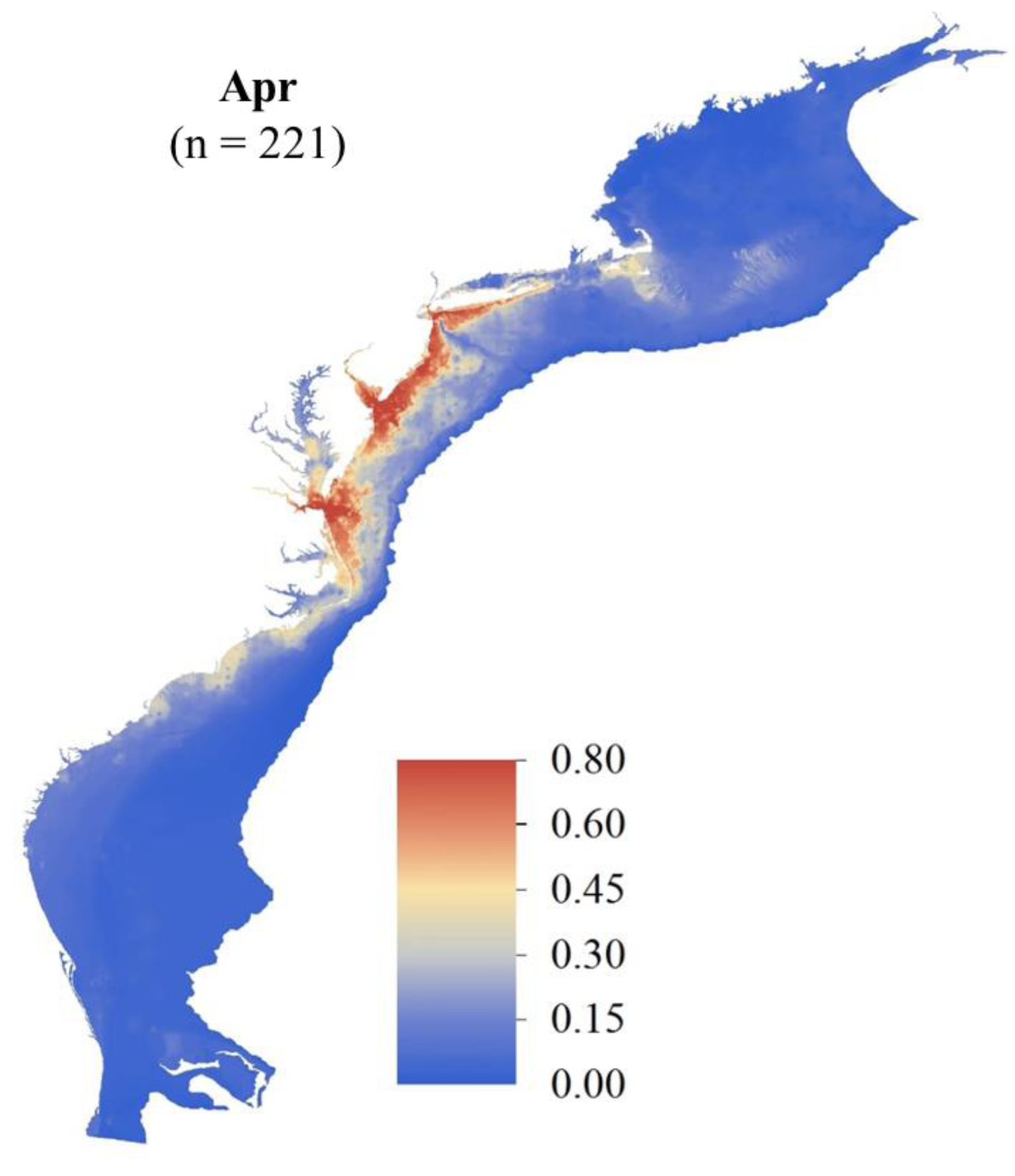
April SDM ensemble map of predicted Atlantic Sturgeon occurrence probabilities in the OCS (raster resolution = 1-km; total area = 626,204 km^2^). Count of total unique detection locations across the OCS is indicated.

**Figure S5.**
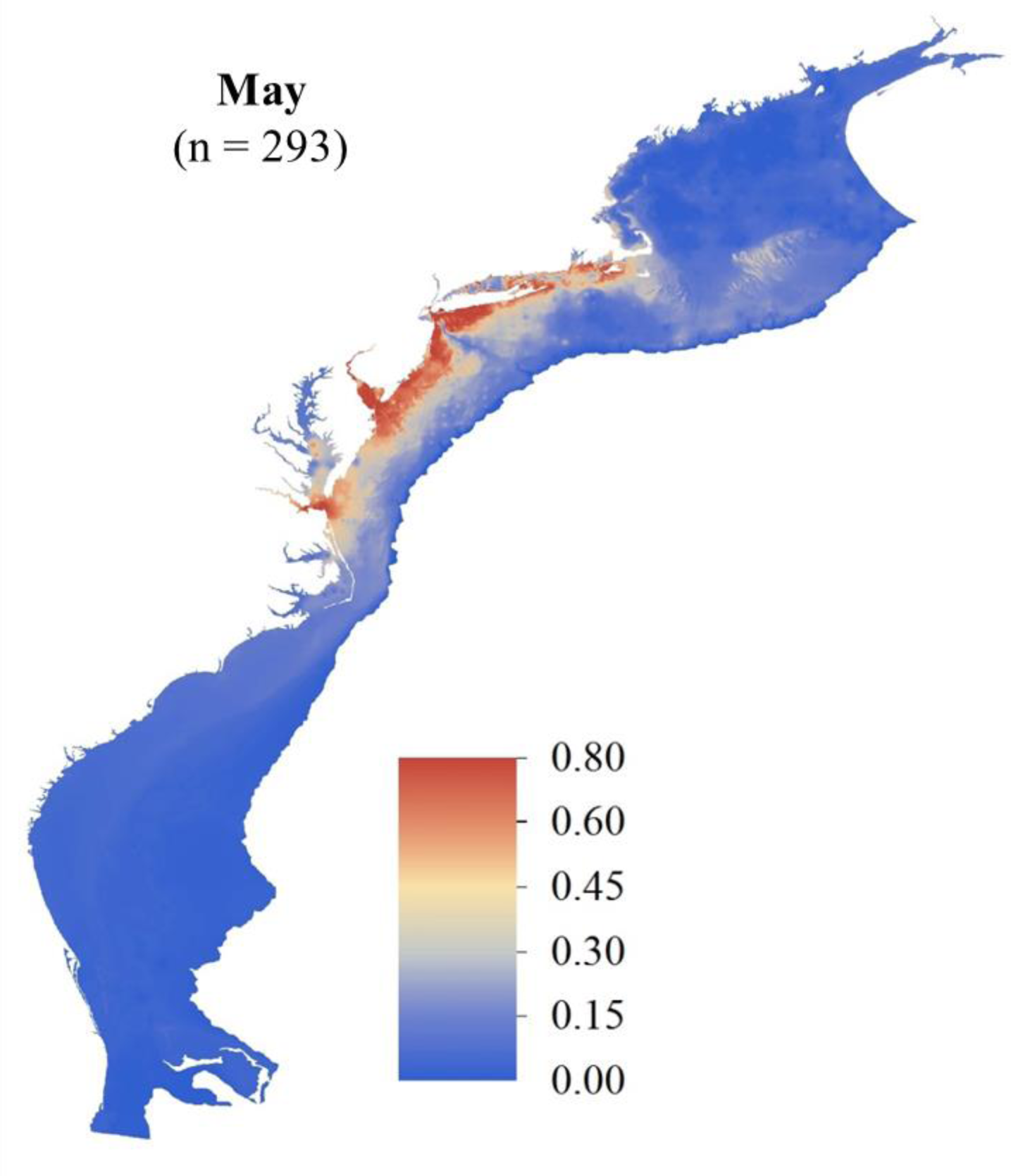
May SDM ensemble map of predicted Atlantic Sturgeon occurrence probabilities in the OCS (raster resolution = 1-km; total area = 626,204 km^2^). Count of total unique detection locations across the OCS is indicated.

**Figure S6.**
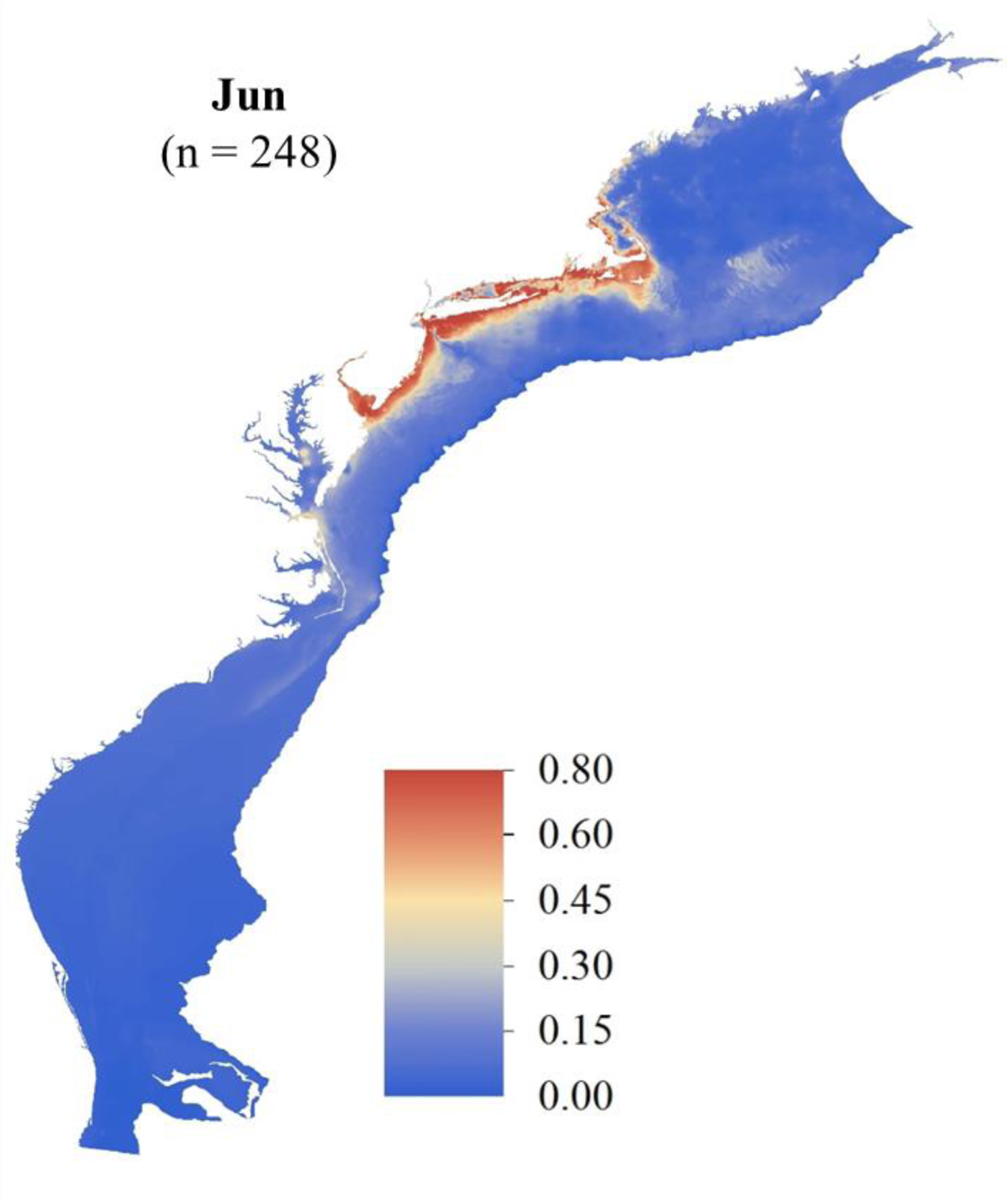
June SDM ensemble map of predicted Atlantic Sturgeon occurrence probabilities in the OCS (raster resolution = 1-km; total area = 626,204 km^2^). Count of total unique detection locations across the OCS is indicated.

**Figure S7.**
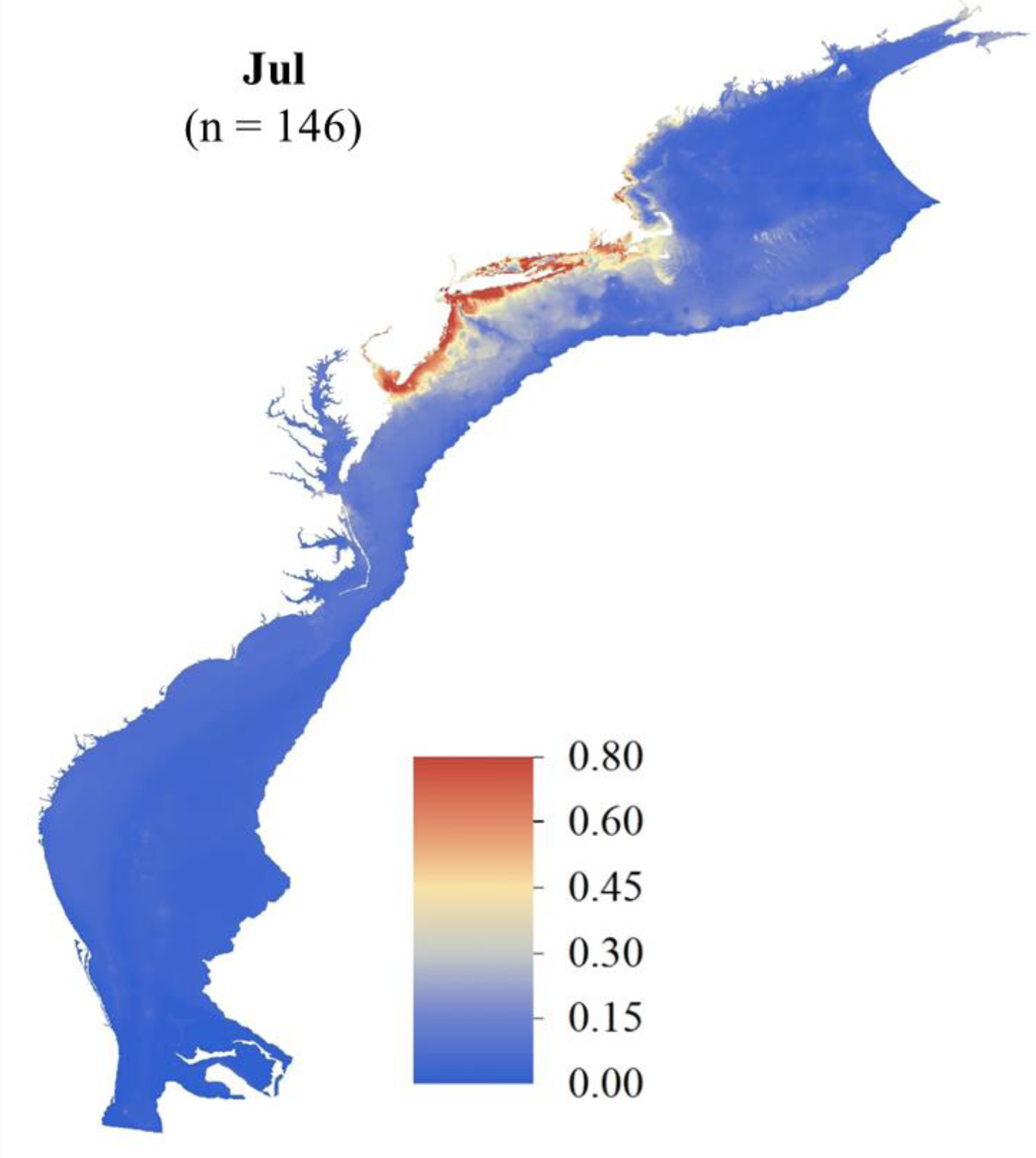
July SDM ensemble map of predicted Atlantic Sturgeon occurrence probabilities in the OCS (raster resolution = 1-km; total area = 626,204 km^2^). Count of total unique detection locations across the OCS is indicated.

**Figure S8.**
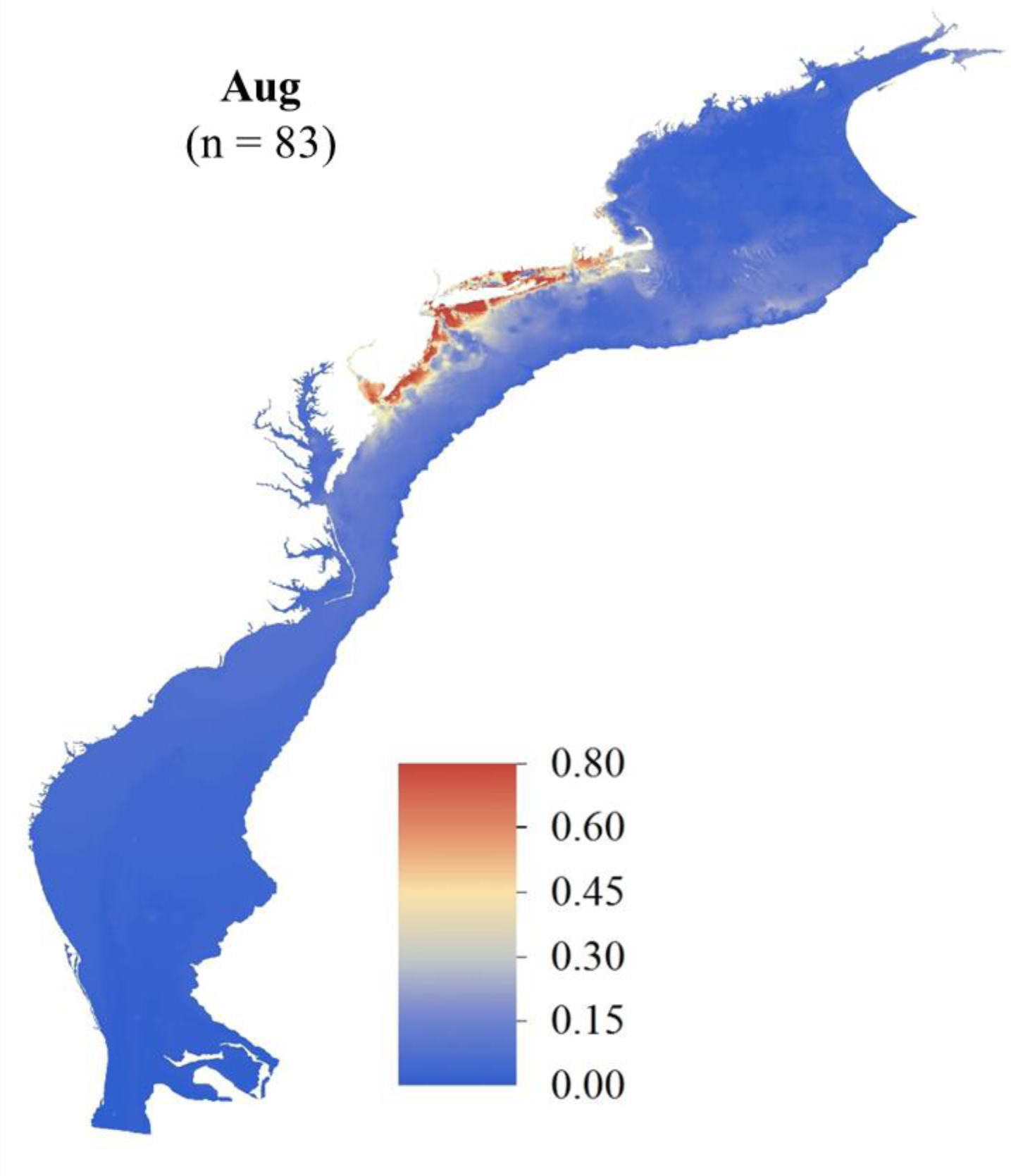
August SDM ensemble map of predicted Atlantic Sturgeon occurrence probabilities in the OCS (raster resolution = 1-km; total area = 626,204 km^2^). Count of total unique detection locations across the OCS is indicated.

**Figure S9.**
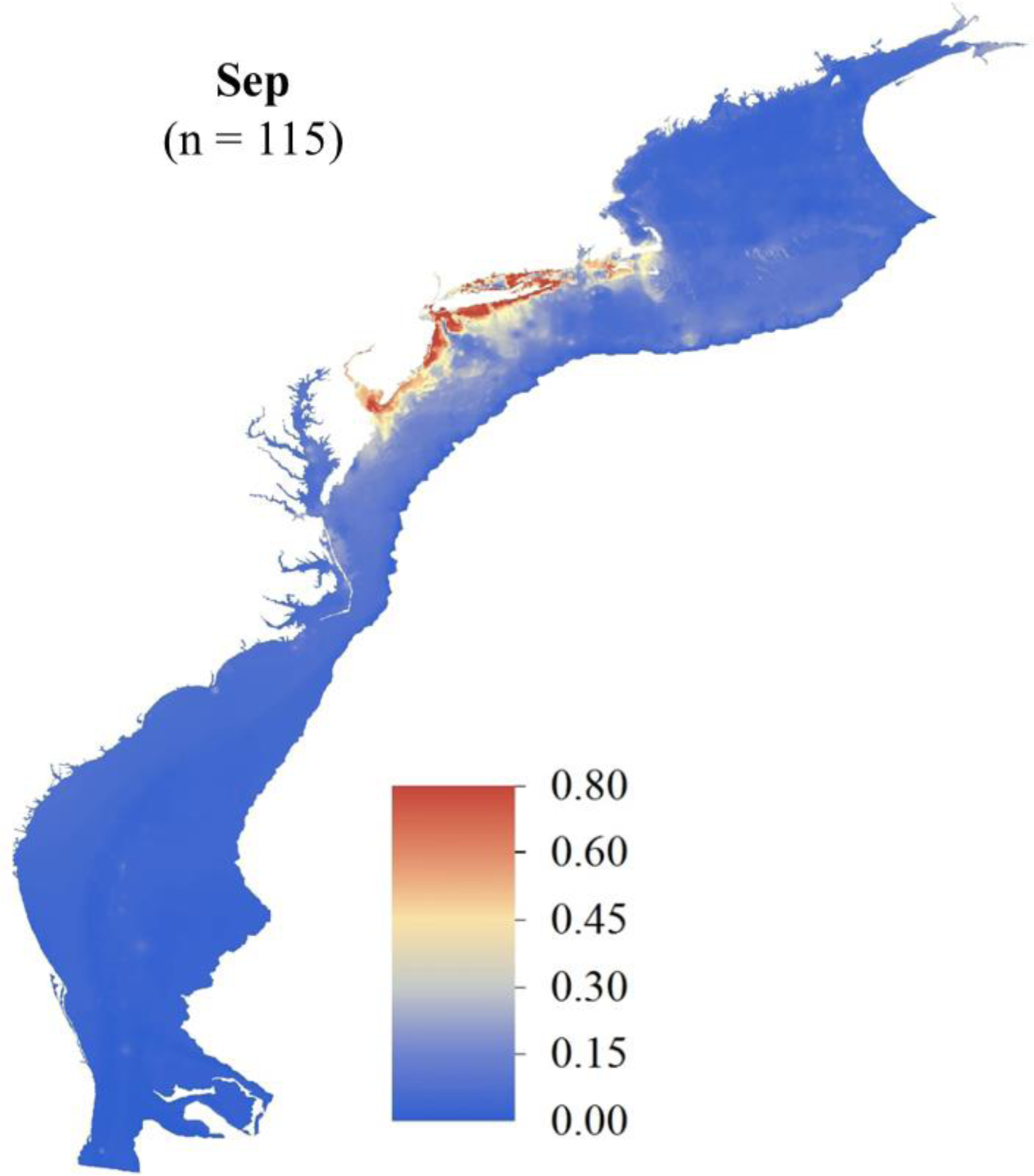
September SDM ensemble map of predicted Atlantic Sturgeon occurrence probabilities in the OCS (raster resolution = 1-km; total area = 626,204 km^2^). Count of total unique detection locations across the OCS is indicated.

**Figure S10.**
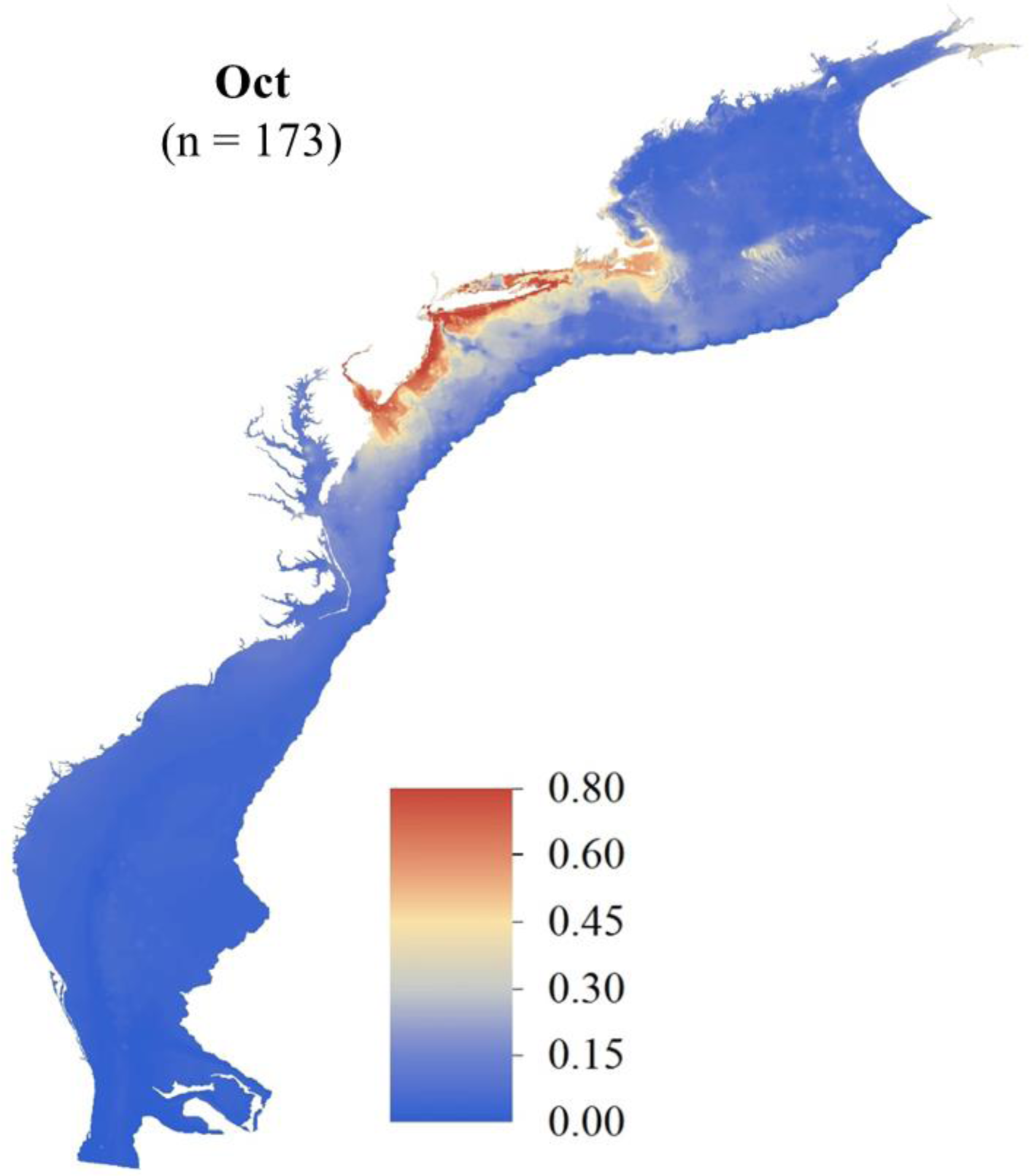
October SDM ensemble map of predicted Atlantic Sturgeon occurrence probabilities in the OCS (raster resolution = 1-km; total area = 626,204 km^2^). Count of total unique detection locations across the OCS is indicated.

**Figure S11.**
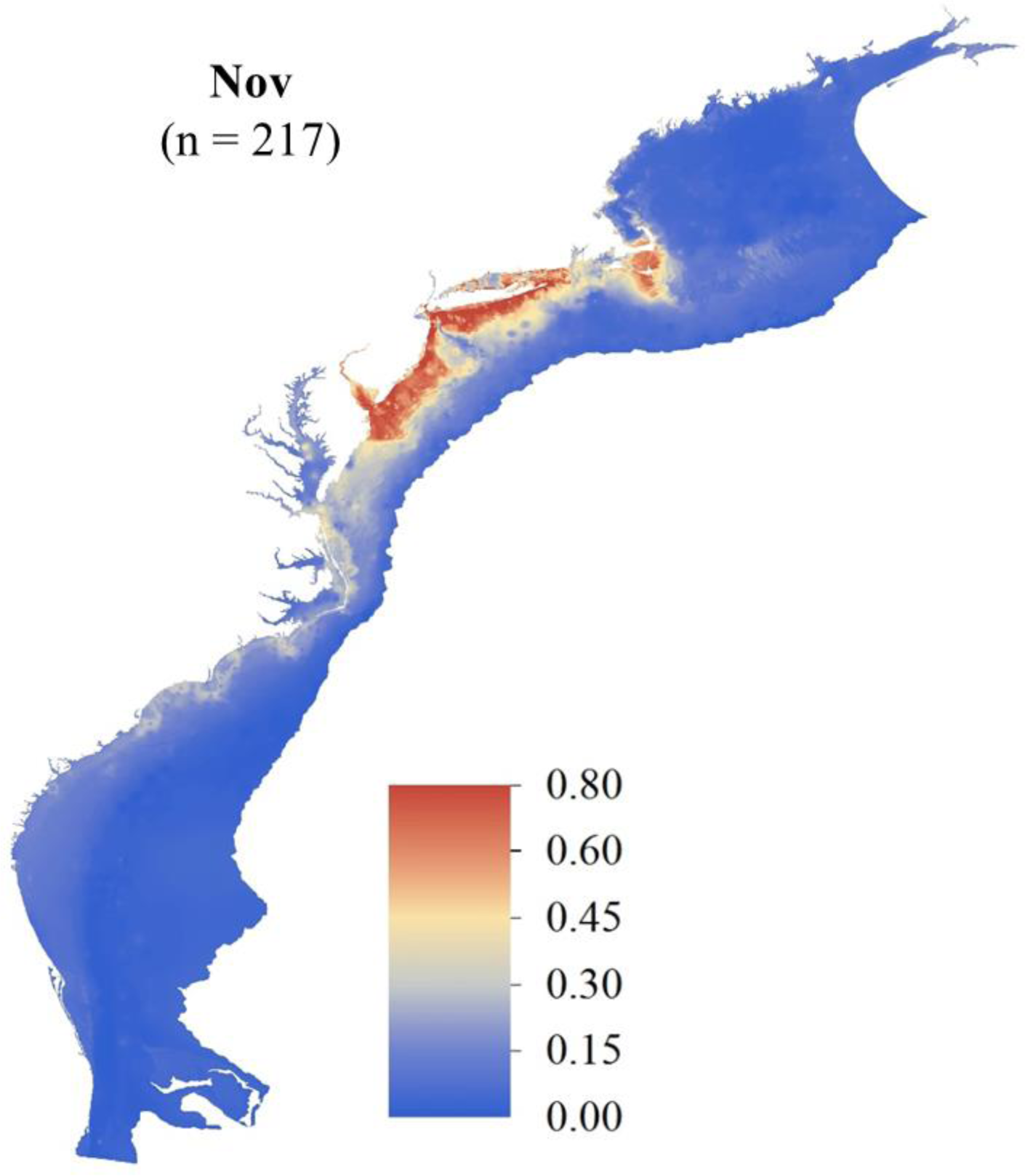
November SDM ensemble map of predicted Atlantic Sturgeon occurrence probabilities in the OCS (raster resolution = 1-km; total area = 626,204 km^2^). Count of total unique detection locations across the OCS is indicated.

**Figure S12.**
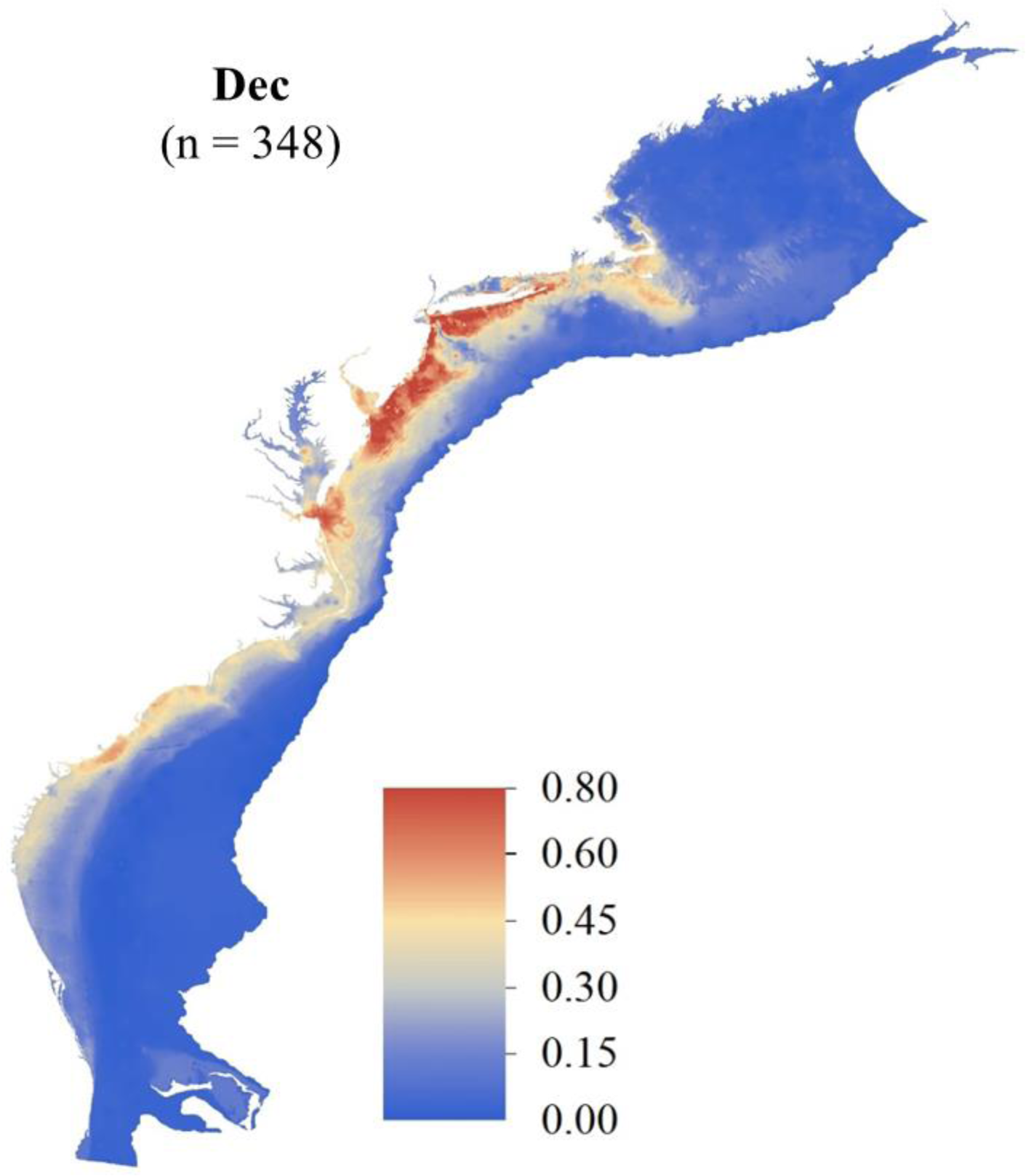
December SDM ensemble map of predicted Atlantic Sturgeon occurrence probabilities in the OCS (raster resolution = 1-km; total area = 626,204 km^2^). Count of total unique detection locations across the OCS is indicated.

**Figure S13.**
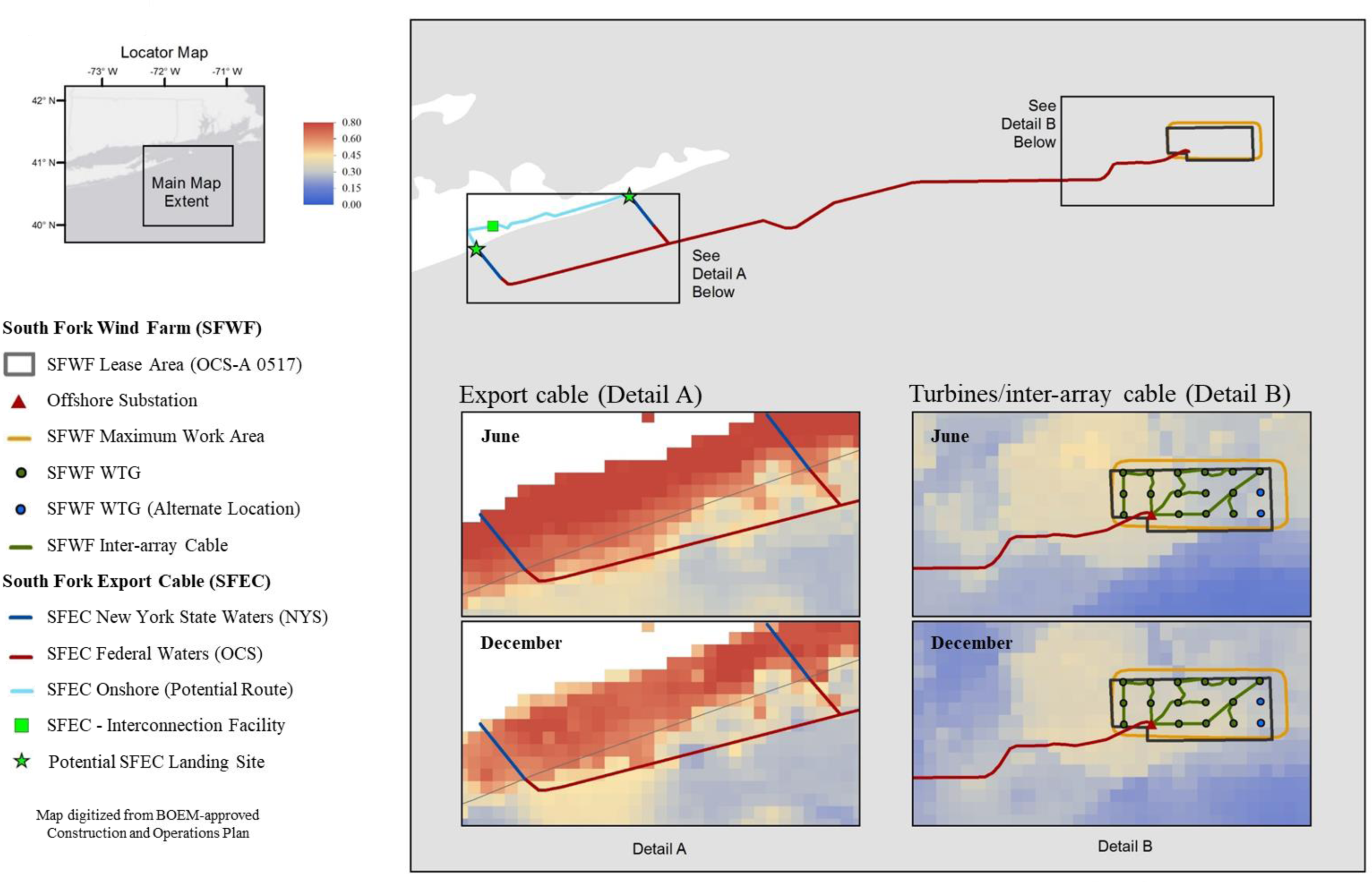
Predicted occurrence of Atlantic Sturgeon in the proposed South Fork Wind Farm (OCS-A 0517) located in the New York Bight off coastal New York and New Jersey. Comparative details for the export cable sites in state waters (detail A) and the turbine/inter-array cable (detail B) are shown for June and December (raster resolution = 1 km).

## Notes

### Competing Interest Statement

The authors have declared no competing interest.

## References

Akhtar N., B. Geyer, and C. Schrum. 2022. Impacts of accelerating deployment of offshore windfarms on near-surface climate. Scientific Reports 12:10.1038/s41598-022-22868-9.

Allouche, O., T. Asaf, and R. Kadmon. 2006. Assessing the accuracy of species distribution models: prevalence, kappa and the true skill statistic (TSS). Journal of Applied Ecology 43:1223–1232.

Alola, A. A., and S. S. Akadiri. 2021. Clean energy development in the United States amidst augmented socioeconomic aspects and country-specific policies. Renewable Energy 169:221 – 230.

Araújo, M. B., R. G. Pearson, W. Thuiller, and M. Erhard. 2005. Validation of species-climate impact models under climate change. Global Change Biology 11:1504–1513.

Araújo, M. B., and M. New. 2007. Ensemble forecasting of species distributions. Trends in Ecology and Evolution 22:42–47.

Backstrom, J. T., and N. M. Warden. 2023. Optimizing offshore wind export cable routing using GIS-based environmental heat maps, Wind Energy Science Discussions in review: 10.5194/wes-2023-146.

Bailey, H., K. L. Brookes, and P. M. Thompson. 2014. Assessing environmental impacts of offshore wind farms: lessons learned and recommendations for the future. Aquatic Biosystems 10:8.

Bangley, C. W., F. G. Whoriskey, J. M. Young, and M. B. Ogburn. 2020. Networked animal telemetry in the Northwest Atlantic and Caribbean waters. Marine and Coastal Fisheries 12:339– 347.

Barbet-Massin, M., F. Jiguet, C. H. Albert, and W. Thuiller. 2012. Selecting pseudo-absences for species distribution models: How, where and how many? Methods in Ecology and Evolution 3:327–338.

Bergstrom, L., F. Sundqvist, and U. Bergstrom. 2013. Effects of an offshore wind farm on temporal and spatial patterns in the demersal fish community. Marine Ecology Progress Series 485:199–210.

Bergstrom, L., L. Kautsky, T. Malm, R. Rosenberg, M. Wahlberg, N. A. Capetillo, and D. Wilhelmsson. 2014. Effects of offshore wind farms on marine wildlife—a generalized impact assessment. Environmental Research Letters 9:034012.

Boakes, E. H., P. J. K. McGowan, R. A. Fuller, D. Chang-qing, N. E. Clark, K. O’Connor K, G. M. Mace. 2010. Distorted views of biodiversity: spatial and temporal bias in species occurrence data. PLoS Biology 8:e1000385.

Boehlert, G. W., and A. B. 2010. Environmental and ecological effects of ocean renewable energy development. Oceanography 23:68–81.

Boria, R. A., E. L. Olson, S. M. Goodman, and R. P. Anderson. 2014. Spatial filtering to reduce sampling bias can improve the performance of ecological niche models. Ecological Modelling 275:73–77.

Breece, M. W., D. A. Fox, D. E. Haulsee, I. I. Wirgin, and M. J. Oliver. 2017. Satellite driven distribution models of endangered Atlantic Sturgeon occurrence in the mid-Atlantic Bight. ICES Journal of Marine Science 75:10.1093/icesjms/fsx187.

Butler, L., and R. A. Sanderson. 2022. National-scale predictions of plant assemblages via community distribution models: leveraging published data to guide future surveys. Journal of Applied Ecology 59:1559–1571.

Cech Jr., J. J., and S. I. Doroshov. 2004. Environmental requirements, preferences, and tolerance limits of North American sturgeons. In: Sturgeons and Paddlefish of North America (eds LeBreton, G. T. O., F. W. H. Beamish, and S. R. McKinley). Springer, pp 73 –86.

Chatterjee, S., and A. S. Hadi. 2015. Regression analysis by example. John Wiley and Sons.

Cohen J. 1960. A coefficient of agreement for nominal scales. Education and Psychological Measurement 20:37–46.

Dingle, H., and V. A. Drake. 2007. What is migration? BioScience 57:113 –121.

Dorman C. F., J. M. McPherson, M. B. Araújo, R. Bivand, J. Bolliger, G. Carl, R. G. Davies, A. Hirzel, W. Jetz, W D. Kissling, I. Kühn, R. Ohlemüller, P. R. Peres-Neto, B. Reineking, B. Schröder, F. M. Schurr, and R. Wilson, 2007. Methods to account for spatial autocorrelation in the analysis of species distributional data: a review. Ecography 30:609 –628.

Drewitt, A. L., and R. H. W. Langston. 2006. Assessing the impacts of wind farms on birds. Ibis 148:29–42.

Dunton, K. J., A. Jordaan, K. A. McKown, D. O. Conover, and M. G. Frisk. 2012. Abundance and distribution of Atlantic Sturgeon (*Acipenser oxyrinchus*) within the Northwest Atlantic Ocean, determined from five fishery-independent surveys. Fishery Bulletin 108:450 –465.

Dunton K. J., D. Chapman, A. Jordaan, K. Feldheim, S. J. O’Leary, K. A. McKown, M. G. Frisk. 2012. Genetic mixed-stock analysis of Atlantic sturgeon *Acipenser oxyrinchus oxyrinchus* in a heavily exploited marine habitat indicates the need for routine genetic monitoring. Journal of Fish Biology 80:207–217.

Elith, J., and J. Leathwick. 2009. Species distribution models: ecological explanation and prediction across space and time. Annual Review of Ecology, Evolution, and Systematics 40:677–697.

Elith, J., S. J. Phillips, T. Hastie, M. Dudík, E. C. Yung, and C, J. Yates. 2011. A statistical explanation of MaxEnt for ecologists. Diversity and Distributions 17:43 –57.

Erickson, D. L., A. Kahnle, M. J. Millard, E. A. Mora, M. Bryja, A. Higgs, J. Mohler, M. DuFour, G. Kenney, J. Sweka, and E. K. Pikitch. 2011. Use of pop-up satellite tags to identify oceanic-migratory patterns for adult Atlantic Sturgeon, *Acipenser oxyrinchus oxyrinchus* Mitchell, 1815. Journal of Applied Ichthyology 27:356 –365.

ESRI. 2018. ArcGIS Desktop: Release 10. Redlands, CA: Environmental Systems Research Institute.

Evans, J. S., M. A. Murphy, Z. A. Holden, and S. A. Cushman. 2011. Modeling species distribution and change using random forest. In: Predictive Species and Habitat Modeling in Landscape Ecology. Springer Science, pp. 139 –159.

Franklin J. 2010. Mapping and species distributions: spatial inference and prediction. Cambridge University Press.

Fielding, A. H., and J. F. Bell. 1997. A review of methods for the assessment of prediction errors in conservation presence/absence models. Environmental Conservation 24:38–49.

Fourcade, Y., J. O. Engler, D. Rödder, and J. Secondi. 2014. Mapping species distributions with MaxEnt using a geographically biased sample of presence data: a performance assessment of methods for correcting sampling bias. PLoS ONE 9:e97122.

Galparsoro, I., I. Menchaca, J. M. Garmendia, A. Borja, A. D. Maldonado, G. Iglesias, and J. Bald. 2022. Reviewing the ecological impacts of offshore wind farms. Ocean Sustainability 1: 10.1038/s44183-022-00003-5.

Gaul, W., D. Sadykova, H. J. White, L. Leon-Sanchez, P. Caplat, M. C. Emmerson, and J. M. Yearsley. 2020. Data quantity is more important than its spatial bias for predictive species distribution modelling. PeerJ 8:e10411.

Gill, A. B. 2005. Offshore renewable energy: ecological implications of generating electricity in the coastal zone. Journal of Applied Ecology 42:605 –615.

Gross, M. R., R. M. Coleman, and R. M. McDowall. 1988. Aquatic productivity and the evolution of diadromous fish migration. Science 239:1291 –1293.

Guisan A., and W. Thuiller. 2005. Predicting species distribution: offering more than simple habitat models. Ecology Letters 8:993–1009.

Gușatu, L. F., S. Menegon, D. Depellegrin, C. Zuidema, A. Faaji, and C. Yamu. 2021. Spatial and temporal analysis of cumulative environmental effects of offshore wind farms in the North Sea basin. Scientific Reports 11:10125.

Hastie, T., R. Tibshirani, and J. Friedman. 2009. The elements of statistical learning: data mining, inference, and prediction. (2nd ed). Springer Science and Business Media.

Hutchison, Z. L., D. H. Secor, and A. B. Gill. 2020. The interaction between resource species and electromagnetic fields associated with electricity production by offshore wind farms. Oceanography 33:96–107.

Ingram, E. C., R. M. Cerrato, K. J. Dunton, and M. G. Frisk. 2019. Endangered Atlantic Sturgeon in the New York Wind Energy Area: implications of future development in an offshore wind energy site. Scientific Reports 9:12432.

Jiménez-Valverde, A., P. Acevedo, A. M. Barbosa, J. M. Lobo, and R. Real. 2013. Discrimination capacity in species distribution models depends on the representativeness of the environmental domain. Global Ecology and Biogeography 22: 508 –516.

Johnson, J. H., D. S. Dropkin, B. E. Warkentine, J. W. Rachlin, and W. D. Andrews. 1997. Food habits of Atlantic Sturgeon off the central New Jersey coast. Transactions of the American Fisheries Society 126:166–170.

Leathwick, J. R., D. Rowe, J. Richardson, J. Elith, and T. Hastie. 2005. Using multivariate adaptive regression splines to predict the distributions of New Zealand’s freshwater diadromous fish. Freshwater Biology 50:2034–2052.

Marmion, M., M. Luoto, R. K. Heikkinen, and W. Thuiller. 2009a. The performance of state-of-the-art modelling techniques depends on geographical distribution of species. Ecological Modelling 220:3512–3520.

Marmion, M., M. Parviainen, M. Luoto, R. K. Heikkinen, and W. Thuiller. 2009b. Evaluation of consensus methods in predictive species distribution modelling. Diversity and Distributions 15:59–69.

Marquardt, D. W. 1970. Generalized inverses, ridge regression, biased linear estimation and nonlinear estimation. Technometrics 12:591–612.

Melnychuk, M. C., K. J. Dunton, A. Jordaan, K. A. McKown, and M. G. Frisk. 2017. Informing conservation strategies for the endangered Atlantic Sturgeon using acoustic telemetry and multi-state mark–recapture models. Journal of Applied Ecology 54:914 –925.

Mi, C., F. Huettmann, Y. Guo, X. Han, and L. Wen. 2017. Why choose Random Forest to predict rare species distribution with few samples in large undersampled areas? Three Asian crane species models provide supporting evidence. PeerJ 5:e2849.

Miller, J., and J. Rogan. 2007. Using GIS and remote sensing for ecological modleing and monitoring. In: Mesev, V. (Ed.). Integration of GIS and Remote Sensing. Wiley, pp 233 –268.

Miller, J., J. Franklin, and R. Aspinall. 2007. Incorporating spatial dependence in predictive vegetation models. Ecological Modelling 202:225–242.

Miller, J. 2010. Species distribution modeling. Geography Compass 4:490 –509.

Naimi, B., and M. B. Araújo. 2016. sdm: a reproducible and extensible R platform for species distribution modelling. Ecography 39:368–375.

Nazir, M. S., Y. Wang, B. Muhammad, and A. N. Abdalla. 2021. Wind energy, its application, challenges, and potential environmental impact. In: Lackner, M., B. Sajjadi, and W. Y. Chen (eds). Handbook of Climate Change Mitigation and Adaptation. Springer, New York, NY.

Papoulious, D. M., A. J. DeLonay, M. L. Annis, M. L. Wildhaber, and D. E. Tillitt. 2011. Characterization of environmental cues for initiation of reproductive cycling and spawning in shovelnose sturgeon *Scaphirhynchus platorynchus* in the Lower Missouri River, USA. Journal of Applied Ichthyology 27:335–342.

Pebesma, E. 2018. Simple features for R: standardized support for spatial vector data. The R Journal:10439–446.

Phillips, S. J., and M. Dudík. 2008. Modeling of species distributions with MaxEnt: new extensions and a comprehensive evaluation. Ecography 31:161–175.

R Core Team. 2018. R: a language and environment for statistical computing. R Foundation for Statistical Computing. Vienna, Austria.

Rothermel, E. R., M. T. Balazik, J. E. Best, M. W. Breece, D. A. Fox, B. I. Gahagan, D. E. Haulsee, A. L. Higgs, M. H. P. O’Brien, M. J. Oliver, I. A. Park, and D. H. Secor. 2020. Comparative migration ecology of striped bass and Atlantic Sturgeon in the US Southern mid-Atlantic bight flyway. PLoS One 15:e0234442.

Shang, N., C. Weng, and G. A. Hripcsak. 2018. conceptual framework for evaluating data suitability for observational studies. Journal of the American Medical Informatics Association 25:248–258.

Stein, A. B., K. D. Friedland, and M. Sutherland. 2004a. Atlantic Sturgeon marine distribution and habitat use along the Northeastern Coast of the United States. Transactions of the American Fisheries Society 133:527–537.

Stein, A. B., K. D. Friedland, and M. Sutherland. 2004b. Atlantic Sturgeon marine bycatch and mortality on the continental shelf of the Northeast United States. North American Journal of Fisheries Management 24:171–183

Sun K., H. Xiao, S. Liu, S. You, F. Yang, Y. Dong, W. Wang, and Y. Liu. 2020. A review of clean electricity policies—from countries to utilities. Sustainability 12(19):7946.

Thuiller W., B. Lafourcade, R. Engler, and M. B. Araújo. 2009. BIOMOD – a platform for ensemble forecasting of species distribution. Ecography 32:369 –373.

Thuiller, W., M. Guéguen, J. Renaud, D. N. Karger, and N. Zimmerman. 2019. Uncertainty in ensembles of global biodiversity scenarios. Nature Communications 10:1446.

USOFR (United States Office of the Federal Register). 2012. Endangered and threatened wildlife and plants; threatened and endangered status for distinct population segments of Atlantic Sturgeon in the northeast region. Federal Register 77:5880 –5912.

Wirgin, I., L. Maceda, C. Grunwald, and T. L. King. 2015. Population origin of Atlantic Sturgeon *Acipenser oxyrinchus oxyrinchus* by-catch in U.S. Atlantic coast fisheries. Journal of Fish Biology 86:1251–1270.

Woodman, S. M., K. A. Forney, E. A. Becker, M. L. DeAngelis, E. L. Hazen, D. M. Palacios, and J. V. Redfern. 2019. esdm: a tool for creating and exploring ensembles of predictions from species distribution and abundance models. Methods in Ecology and Evolution 10:1923–1933.

Zurell, D., N. E. Zimmermann, H. Gross, A. Baltensweiler, T. Sattler, and R. O. Wüest. 2020. Testing species assemblage predictions from stacked and joint species distribution models. Journal of Biogeography 47:101–113.

